# Domesticated cannabinoid synthases amid a wild mosaic cannabis pangenome

**DOI:** 10.1101/2024.05.21.595196

**Authors:** Ryan C. Lynch, Lillian K. Padgitt-Cobb, Andrea R. Garfinkel, Brian J. Knaus, Nolan T. Hartwick, Nicholas Allsing, Anthony Aylward, Allen Mamerto, Justine K. Kitony, Kelly Colt, Emily R. Murray, Tiffany Duong, Aaron Trippe, Seth Crawford, Kelly Vining, Todd P. Michael

**Author notes:** Correspondence: **Todd P. Michael,**,**, Ryan C. Lynch****, Lillian K. Padgitt-Cobb**.

## Abstract

*Cannabis sativa* is a globally significant seed-oil, fiber, and drug-producing plant species. However, a century of prohibition has severely restricted legal breeding and germplasm resource development, leaving potential hemp-based nutritional and fiber applications unrealized. Existing cultivars are highly heterozygous and lack competitiveness in the overall fiber and grain markets, relegating hemp to less than 200,000 hectares globally^1^. The relaxation of drug laws in recent decades has generated widespread interest in expanding and reincorporating cannabis into agricultural systems, but progress has been impeded by the limited understanding of genomics and breeding potential. No studies to date have examined the genomic diversity and evolution of cannabis populations using haplotype-resolved, chromosome-scale assemblies from publicly available germplasm. Here we present a cannabis pangenome, constructed with 181 new and 12 previously released genomes from a total of 156 biological samples from both male (XY) and female (XX) plants, including 42 trio phased and 36 haplotype-resolved, chromosome-scale assemblies. We discovered widespread regions of the cannabis pangenome that are surprisingly diverse for a single species, with high levels of genetic and structural variation, and propose a novel population structure and hybridization history. Conversely, the cannabinoid synthase genes contain very low levels of diversity, despite being embedded within a variable region containing multiple pseudogenized paralogs and distinct transposable element arrangements. Additionally, we identified variants of *acyl-lipid thioesterase* (*ALT*) genes^2^ that are associated with fatty acid chain length variation and the production of the rare cannabinoids, tetrahydrocannabinol varin (THCV) and cannabidiol varin (CBDV). We conclude the *Cannabis sativa* gene pool has only been partially characterized, and that the existence of wild relatives in Asia remains likely, while its potential as a crop species remains largely unrealized.

## Main

Cannabis (*Cannabis sativa* L., cannabis) is an ancient domesticated plant with archaeological evidence for seed (achene) and fiber utilization dating to about 8,000 years before the present in Japan ^3^. Cannabis was originally a multipurpose crop in Asia, where the same plants were utilized as a significant source of fiber, food, and drugs ^4,5^. Through time, cannabis spread globally and single or dual use type cultivars were developed, eventually giving rise to divergent hemp and drug-type populations of the 20th century ^6^.

Throughout history and around the world, cannabis has been subjected to cycles of “cultivation, consumption, and crackdown” ^7^. Modern prohibition originated in the United States during the early 20th century ^8^, but by 1961 had spread to a majority of countries ^9^. This prohibition eliminated the fiber and food uses of cannabis for decades, but gave rise to a high-value black market for phytocannabinoid based drugs, which are derived from glandular trichomes. While over 100 phytocannabinoids have been identified, only a limited number are produced in significant quantities, which are used to classify plants by chemotype: Delta-9-tetrahydrocannabinolic acid (THCA; type I), cannabidiolic acid (CBDA; type III), balanced CBDA and THCA (type II), cannabigerolic acid (CBGA; type IV), and cannabinoid-free (type V) ^10^. Although tetrahydrocannabinol (THC), the primary intoxicant, remains a controlled substance, a majority of U.S. states, and many countries, now allow medical or adult-use of cannabis products. Separately, the 2014 and 2018 Farm Bills facilitated hemp production and research for plants producing less than 0.3% THC once again on U.S. soil, generating opportunity for improved non-THC drug, grain, and fiber applications.

Cannabis is often classified as a monospecific genus (*Cannabis sativa* L.) ^11^, although debate remains regarding the current and historical status of *Cannabis indica* Lam., and *Cannabis ruderalis*, the latter of which is thought to be the source of the “day-neutral” flowering type ^12,13^.

Since the first cannabis genome release in 2011, progress has been made to bring this valuable plant into the scope of modern scientific study and breeding ^14–19^. Although the CBDRx (cs10) reference genome ^14^ resolved the cannabinoid gene cassette structure and revealed the predominantly marijuana (MJ) background of modern high cannabinoid (hc) yielding hemp, questions remain regarding the extent of global diversity, and how hybridization shapes genome architecture and allele transmission.

Single reference genomes are now being superseded by pangenomes, providing the opportunity to understand diversity and conservation status in a species at the population genome scale. Here we uncover genome architecture of *Cannabis sativa* with a pangenome encompassing 193 genomes and haplotypes spanning 156 samples (including 12 existing public assemblies) from both male (XY) and female (XX) plants of European and Asian fiber and seed hemp, feral populations, North American MJ, hc yielding hemp, and wild Asian populations. We catalog the genetic, transposable element, and structural variations among a diverse panel of 78 haplotype-resolved, chromosome-scale assemblies (42 of which are trio phased), and examine their relationships among populations. These analyses reveal megabase (Mb) scale areas of the genome with divergent structural variations (SVs), which contrast with low diversity regions, such as the functional cannabinoid synthase genes. Additionally, we examine variation in terpene and disease resistance genes, as well as fatty acid biosynthesis genes, which impact the production of minor cannabinoid propyl-homologs (tetrahydrocannabinol varin [THCV] and cannabidiol varin [CBDV]).

### Haplotype-resolved, chromosome-scale “anchor” genomes

We took a trio approach to generate a fully phased diploid (haplotype-resolved) chromosome-scale “anchor” genome that captures both chemotypic as well as agronomic trait variation. In the pangenome context, an “anchor” genome achieves similar high quality and is used as a reference for cross-genome comparisons; a pangenome can include multiple anchor genomes (see pangenome section). While we generated 21 trio phased, haplotype-resolved, chromosome-scale cannabis genomes, we focused on one specific F1 hybrid between two phenotypically and genetically divergent parents to clarify features of the genome missed in other studies to date (Fig. 1a; Supplemental Table 1; Supplemental Fig. 1). EH23 (ERBxHO40_23) is a cross between HO40 (EH23a) and Early Resin Berry (ERB; EH23b).

**Fig. 1.**
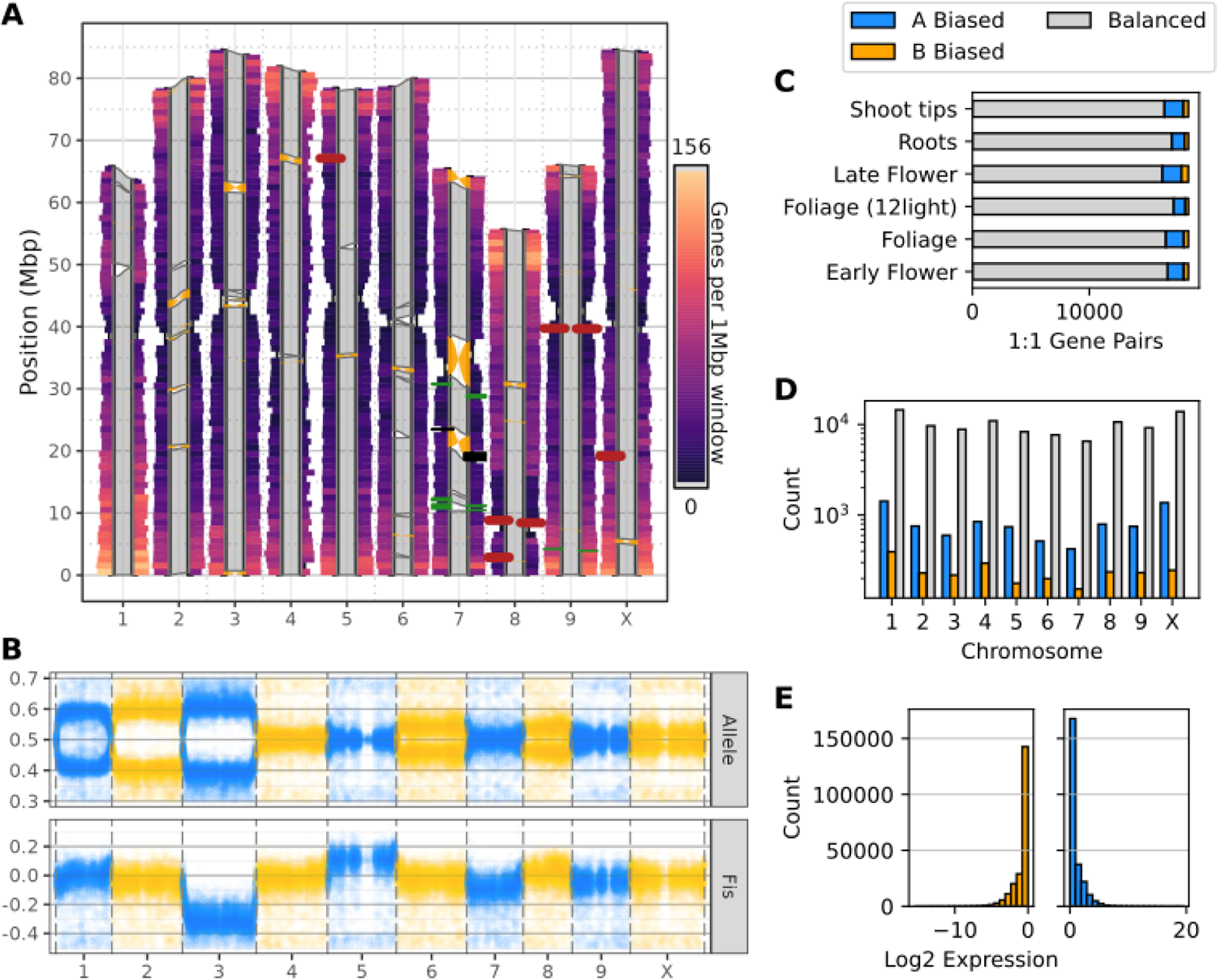
Genomic landscape of *Cannabis sativa*. A) Chromosomal features of nine pairs of autosomes and one pair of sex chromosomes (XX). One million base pair rectangular windows extend outward from each pair of haplotypes at a width proportional to the absence of the CpG motif. Each rectangular window is colored by gene density with warm colors indicating high gene density and cool colors indicating low gene density. Each pair of haplotypes is connected by polygons indicating structural arrangement, with gray for syntenic regions and orange connecting inversions. Rectangles along each haplotype indicate select loci, including 45S (26S, 5.8S, 18S) RNA arrays (firebrick red), 5S RNA arrays (black), and cannabinoid synthases (forest green; CBCAS, CBDAS, THCAS, and olivetolic acid cyclase [OAC]). B) Inheritance of alleles across the genome from the F2 population. The upper panel presents the frequency of each allele and the lower panel shows *F_IS_* or the deviation from our evolutionarily neutral expectation of heterozygosity. C) Haplotype specific expression by tissue and sample type. D) Haplotype specific expression by chromosome. E) Global haplotype specific expression analysis.

HO40 is type I propyl cannabinoid (THCVA and THCA)-producing, short day (SD) flowering responsive, and is part of the drug-type group with a closer affinity to Asian hemp (MJ), while ERB is a type III pentyl cannabinoid (CBDA)-producing, day-neutral (DN) flowering, and is part of the drug-type group more closely related to European hemp (hc hemp) (Supplemental Fig. 2).

**Fig. 2.**
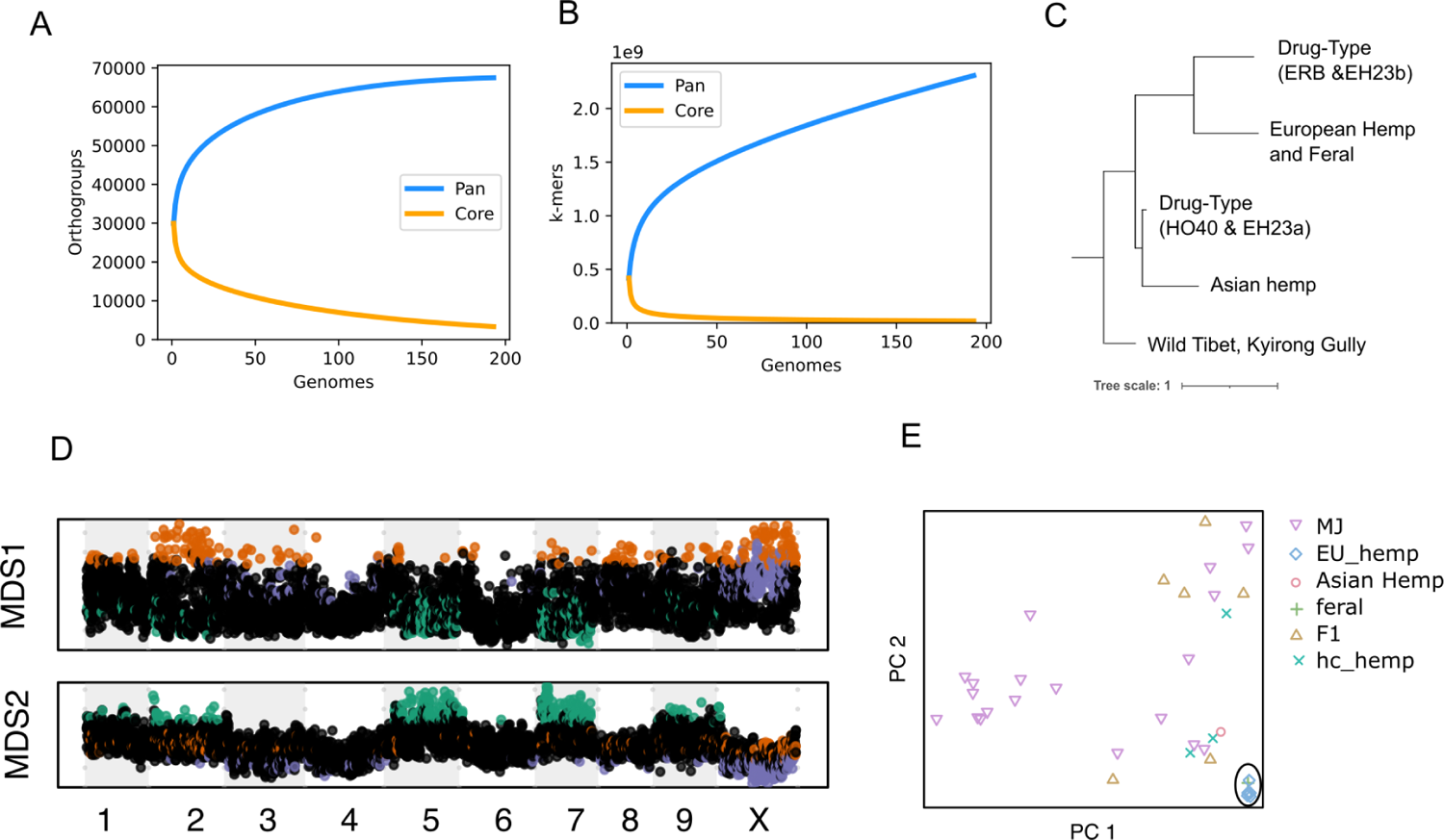
Cannabis pangenome reveals at least five distinct populations. A) Collector’s curve using shared gene orthogroup membership, showing pangenome orthogroup diversity plateaus around 125 samples. B) Collector’s curve using shared 31-mers, suggesting additional Cannabis pangenome diversity remains unsampled. C) Hierarchical clustering of Jaccard similarity scores based on 31-mers reveals a structure of at least major five groups across the 193 pangenome samples. Each drug type group contains both MJ and hc hemp samples, which are fully detailed in Supplemental Fig. 6d) D) Local PCA of phased SNPs from the 78 haplotype-resolved, chromosome-scale assemblies. Two multidimensional scaling (MDS) axes of the PCA distances for 200 SNP genomic windows plotted along the EH23a chromosomes. E) PCA of green outlier MDS SNP windows (from D) contain regions of the genome that show similar ancestry across all European hemp and feral North American samples (inside of ellipse, lower right), with widespread occurrence across chromosomes 2, 5, 7, and 9. The Asian hemp sample (red circle, lower right quadrant) appears closer to marijuana (red triangles pointing down; MJ) and high cannabinoid (blue crosses; hc hemp) hemp samples. Population designations are derived from initial genome wide PCA and known use type of samples (Supplemental Fig. 7).

The EH23 anchor genome revealed differences in chromosome lengths, putative centromere regions, structural variation (SV), distinct locations of THCAS and CBDAS clusters, and gene densities (Fig. 1a; Supplemental Materials), which we also visualized across the other haplotype-resolved, chromosome-scale cannabis genomes (Data Availability). The abundance of the motif ‘CpG’ (cytosine, phosphate, guanine), which is correlated with methylation and centromere formation, varied throughout the chromosomes yet showed clear concentrations along the chromosomes in gene poor regions consistent with the centromere. We identified high copy number repeats consistent with centromere arrays, yet one was not found consistently across the chromosomes and the other was the sub-telomeric repeat^20^ (Supplemental Fig. 3).

**Fig. 3.**
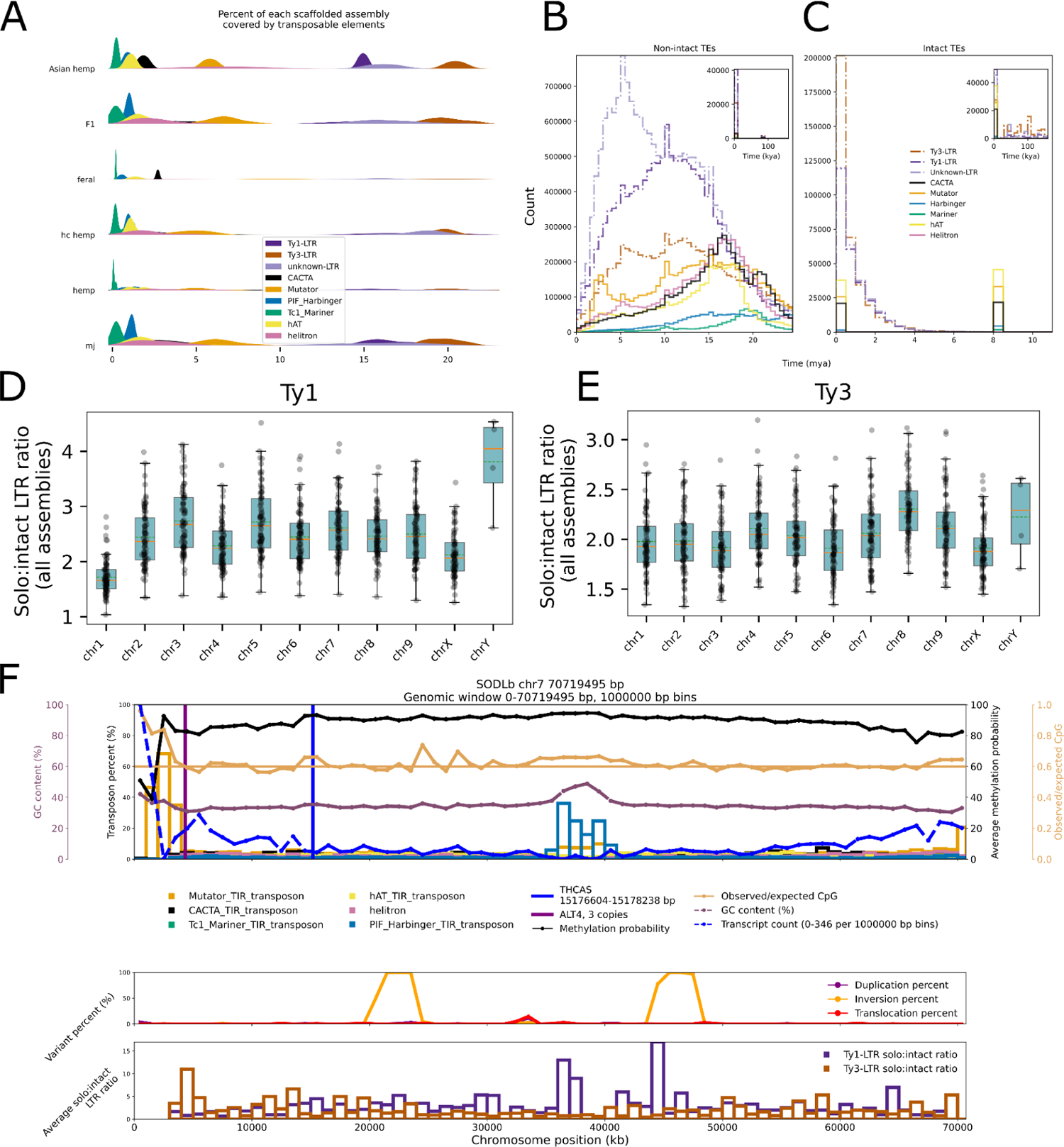
Transposable elements shape the cannabis pangenome. A) Percent of each haplotype-resolved, chromosome-scale assembly covered by TEs, grouped by population ID. The y-axis is a Gaussian kernel density estimation (KDE). B) Age distribution of fragmented TEs (million years ago [mya]), with inset showing distribution within the last 100,000 years (thousand years ago [kya]). In the inset, the highest density occurs within 10 kya. C) Age distribution of intact TEs (million years ago [mya]), with inset showing distribution within the last 100,000 years (thousand years ago [kya]). In the inset, the highest density occurs within 10 kya. D) Solo:intact ratio for Ty1-LTR elements. E) Solo:intact ratio for Ty3-LTR elements. F) SODLb chromosome 7 (Sour Diesel, type I, population type “MJ”) view of DNA TEs, average methylation probability, GC content, and transcript count. Each bin spans 1 Mb, except for the last bin, which depicts the remainder length of the chromosome (719,495 bp).

Long telomere sequences (∼50 Kb; AAACCCT) were found on all chromosome ends and the sub-telomeric repeat^20^ was found on all but four chromosome ends (4/20) and was also found in the putative (high CpG) centromere regions in half (10/20) of the chromosomes (Fig. 1a; Supplemental Fig. 3).

An F2 population derived from selfing EH23 (ERBxHO40_23) exhibits mixed patterns of allele frequencies and F_is_ (deviation of heterozygosity from Hardy-Weinberg expectation) across the genome, likely the consequence of segregation distortion as seen in other cannabis crosses^19^ (Fig. 1b). Segregation distortion patterns observed across many regions of the genome were noted in a prior study of this population ^21^, and may be caused by differences in seed germination rates. A study done by Beutler and Der Marderosian reported a delay in seed germination in crosses between DN and SD plants; the authors speculate this trait is a carryover from the hypothesized species “*Cannabis ruderalis*,” or DN, parent^22^. Alternatively, prezygotic selection such as meiotic drive in the female germline or pollen competition could drive deviations from Hardy-Weinberg equilibrium ^23^. In *Arabidopsis* and rice many tested populations show segregation distortion, which may be due to seed dormancy and lethal epistatic interactions ^24,25^. However, the segregation distortion could be the result of the SV noted across the pangenome, and consistent with this chromosome 1, where DN alleles have been mapped ^21,26^, displays the largest size variation (Supplemental Fig. 4).

**Fig. 4.**
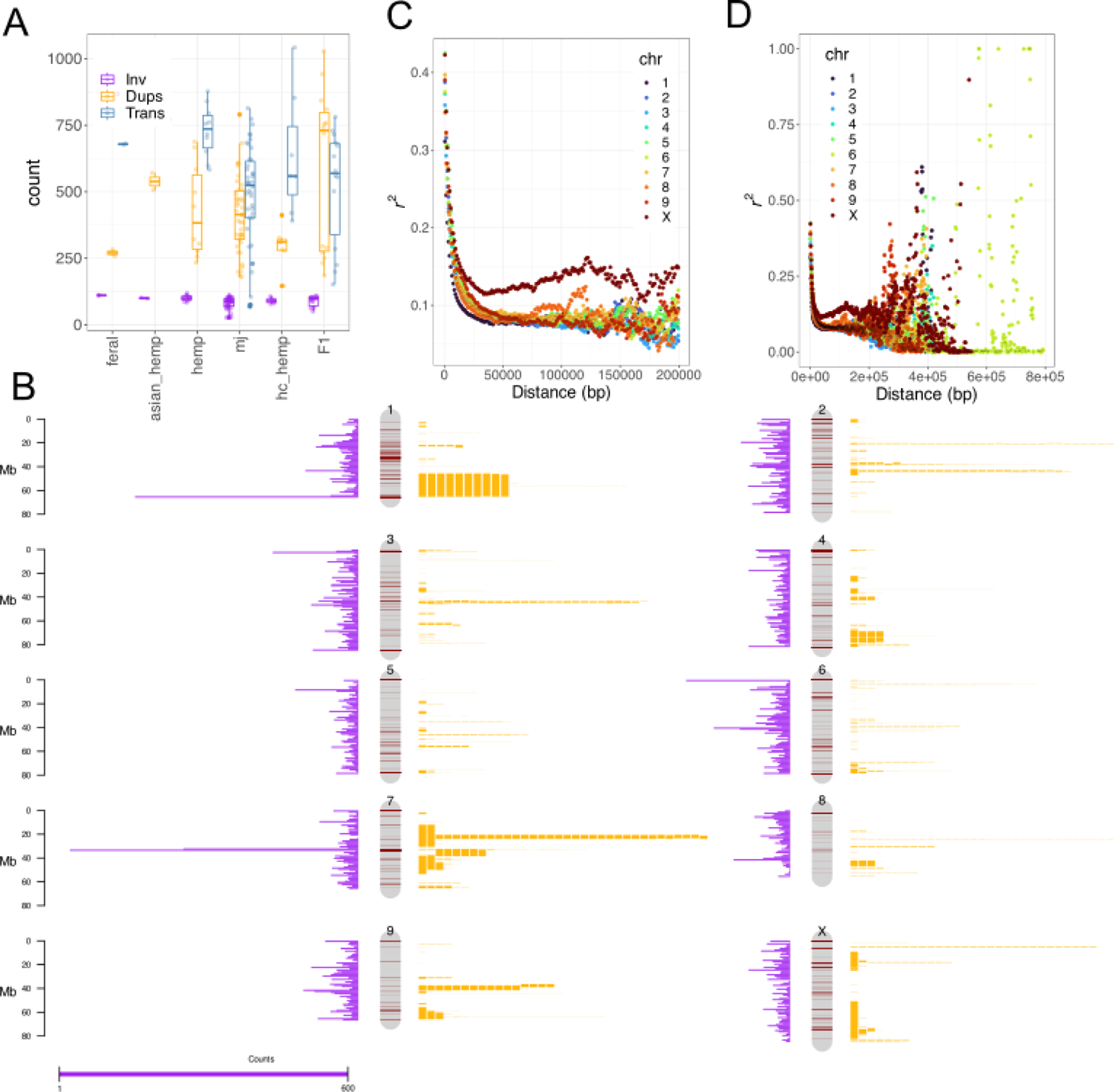
Structural variants occur at different frequencies in various populations and are non-randomly distributed across the genome. A) Frequencies of inversions, duplications, and translocations by population. European hemp, Asian hemp, and marijuana populations differ significantly in average translocation and duplication counts, but not inversions. B) Non-random genomic distribution of translocations (purple histograms), duplications (dark red bands) and inversions (mapped as length-scaled yellow bars on the right side of chromosomes, with each bar equal to one inversion). C) Linkage disequilibrium (LD) plot limited to 200 Kb interactions highlighting the general decay curves, with the X chromosome exhibiting a markedly reduced decay rate. D) LD decay plots extended to 800 Kb.

The EH23 trio phased diploid F1 assembly provides the opportunity to evaluate haplotype specific expression patterns in cannabis. We sampled six different tissue types from EH23 (shoot tips, roots, late flower, leaf under short day, leaf under long day, and early flower) to evaluate haplotype specific expression (Supplemental Table 2). Globally, we found that haplotype EH23a alleles had increased expression compared to those of haplotype EH23b (Fig. 1e). Comparisons between tissue types show that ∼7% of expressed transcripts represent EH23a alleles and ∼3% represent those of EH23b (Fig. 1c), while most (90%) of the genes have balanced allele expression based on our criteria. Additionally, we found about 2 - 3x total increased expression between EH23a and EH23b across the chromosomes (Fig. 1D), by merging the tissue expression data sets. In sum, we used the F1 hybrid EH23 trio phased haplotype-resolved, chromosome-scale assembly to characterize haplotype specific transcription patterns, which show an absence of widespread genome silencing or dominance.

### Pangenome uncovers at least five populations

Single reference genomes are now being superseded by pangenomes, providing the opportunity to understand diversity and conservation status in a species at the population scale^27^. The pangenome samples were selected from multiple sources to maximize the genetic diversity, history, sex expression, and agronomic value. A large portion of the pangenome comes from the Oregon CBD (OCBD) breeding program that includes elite cultivars (Fig. 1; EH23a, EH23b); foundational MJ lines potentially originating from the 1970s, 80s, 90s to present; and elite trios (see varin section). A pedigree of the OCBD lines shows how these lines are related and nested (Supplemental Fig. 1). The remaining cultivars come from the United States Department of Agriculture (USDA) Germplasm Resource Information Network (GRIN) and German federal genebank (IPK Gatersleben) repositories to ensure researchers would have access to plants for experimentation. The pangenome includes European and Asian fiber and seed hemp, feral populations, North American marijuana (type I), and hc yielding (CBDA or CBGA) hemp (type III and IV). Additional cannabinoid diversity is represented with chemotypes presenting high expression of pentyl or propyl (varin) homologs of CBDA or THCA, and cannabinoid free (type V) plants. Flowering time variation is also captured with the inclusion of both regular SD and DN (aka autoflowering) phenotypes. Finally, four male (XY) genomes are included in the pangenome to elucidate the sex chromosomes in cannabis (Supplemental Table 1).

We used two methods to achieve phased chromosome scale genomes to serve as potential anchors; the first and simplest relied on HiC data for both phasing and scaffolding (Supplemental Table 1 and Methods). In total 24 such haploid genomes from 12 samples were produced (Hifiasm_HiC) and used as scaffolding references for 42 genomes from 21 samples (Hifiasm_Trio_RagTag) resulting in trio phased haploid assemblies. These 78 genomes form our haplotype-resolved, chromosome-scale assemblies that were used in many of the analyses presented in this pangenome. In addition, we generated 20 haplotype-resolved contig level assemblies (Hifiasm), as well as 83 contig level assemblies based on older PacBio continuous long reads (CLR; 23 assemblies) and circular consensus sequencing (CCS; 60 assemblies) (Supplemental Table 1). All of the genomes are high quality with a median N50 length of 4 Mb and genome Benchmarking Universal Single-Copy Orthologs (BUSCO) ^28^ complete scores around 97% on average (Supplemental Table 1; Supplemental Fig. 4).

We calculated the “collector’s curve” (pangenome rarefication) using shared gene-based orthogroups and shared k-mers (31-mers) (Fig. 2a,b). These reveal that we likely sampled the majority of cannabis orthogroup genetic diversity by around 100 - 125 genomes (Fig. 2a), but that much global genomic variation remains uncharacterized (Fig. 2b). Collector’s curves of the 78 haplotype-resolved, chromosome-scale assemblies show similar, although more attenuated, relationships between diversity and sample numbers (Supplemental Fig. 5). These findings are similar to other super pangenome projects that include multiple species within a genus, such as tomato ^29^ and strawberry ^30^. Additionally, some of the pairwise average F_st_ values based on phased SNPs among these cannabis populations (Supplemental Table 3) are within the range of some interspecies comparisons such as strawberry ^30^, suggesting the long debated taxonomy within the *Cannabis* genus remains unresolved.

**Fig. 5.**
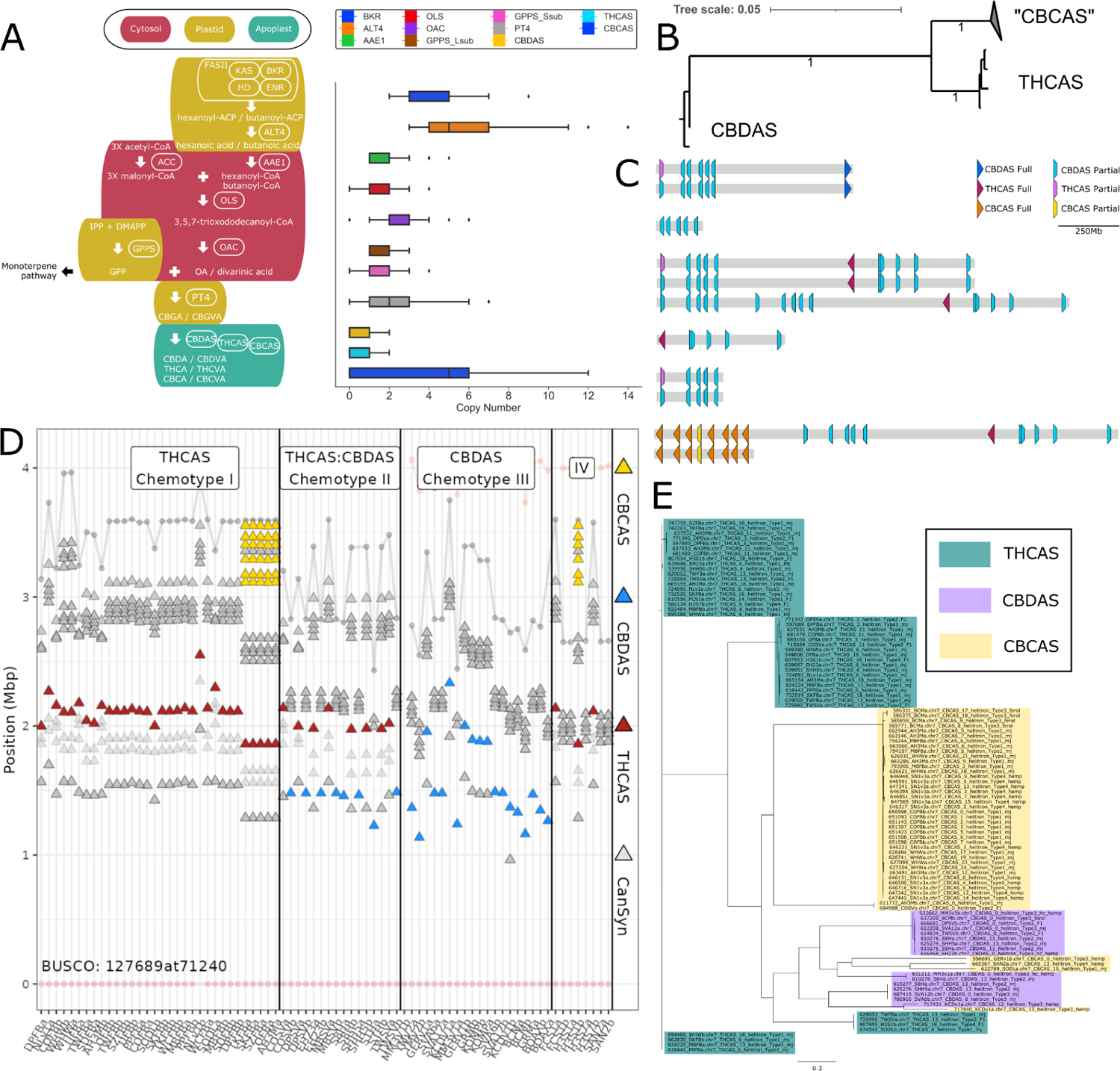
The cannabinoid pathway is domesticated, but shows contrasting patterns of genetic diversity and synteny. A) Cannabinoid biosynthesis pathway and gene copy numbers across the pangenome, per assembly. B) Consensus maximum likelihood phylogeny of aligned coding sequences from cannabinoid synthases, with the proportion of 100 bootstrap replicates shown on branches where values are > 0.75. Each branch tip represents a distinct cluster of synthases within > 99% identity of 859 total synthases from across all 193 pangenome samples. C) Summary of common cannabinoid synthase cassette arrangements. D) Synthase cassettes exhibit variation in synteny as seen in BUSCO anchored local alignment of chromosome 7. Red triangle = THCAS cassettes; Blue triangles = CBDAS cassettes; Yellow triangle = CBCAS cassettes; gray triangles = low stringency synthase-like genes and pseudogenes; gray and pink circles = BUSCOs. E) Maximum likelihood tree of helitron DNA TE sequences flanking (2 Kb upstream or downstream) cannabinoid synthases in the 78 haplotype-resolved, chromosome-scale assemblies.

A broad split of drug-type (high cannabinoid content; hc) samples into two groups, one aligned with Asian hemp and one with European hemp, is suggested by the k-mer based hierarchical clustering (Fig. 2c; Supplemental Fig. 6). Both groups contain MJ and hc hemp samples, which are thought to have largely marijuana ancestry with a recent history of introgression breeding for CBDAS genes, perhaps from European hemp origins ^14^. However, using a phased SNP-based structure with all MJ samples treated as a single population, the TreeMix model inferred a most likely phylogeny that includes six gene flow (migration) events between Asian hemp, hc hemp, and European hemp, as well as MJ and hc hemp samples (Supplemental Fig. 7). These results may partially explain the European and Asian groupings of drug-type samples found by our k-mer clustering analysis, and reflect the effects of historical hybridization breeding between Asian and European hemp that is documented in the breeding literature ^31^.

**Fig. 6.**
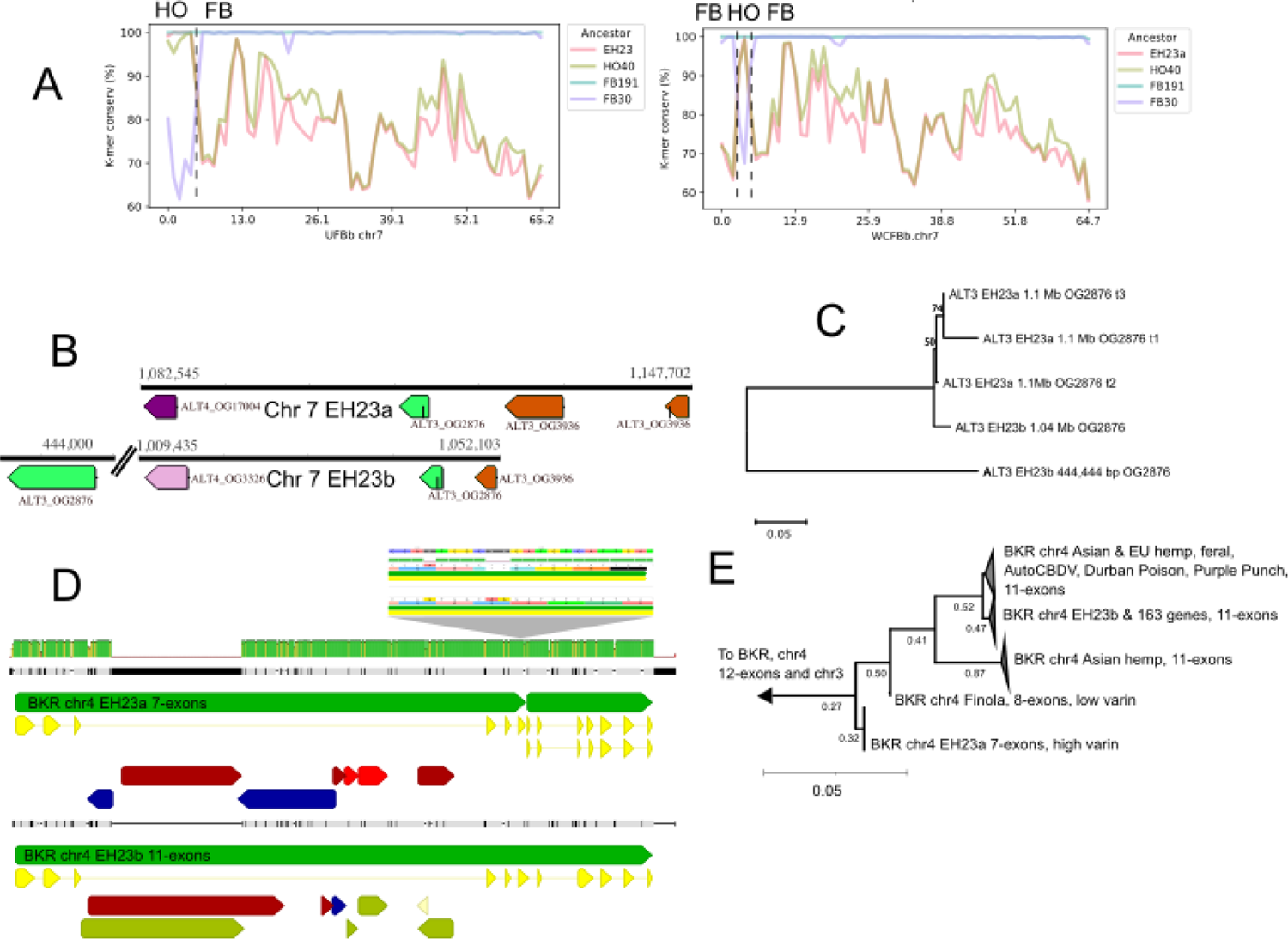
*acyl-lipid thioesterase* (ALT) gene transduplication (TD), and diversification explains varin cannabinoid phenotype in cannabis. A) PanKmer crossover analysis identifies the specific breakpoints on chromosome 7 (vertical dashed lines) for the ALT gene haplotypes in relationship to cannabinoid synthases. UFBb has one crossover at 5 Mb that breaks the linkage between HO40 (HO) THCAS and the varin haplotype ALT genes, while WCFBb has two crossovers, which result in an absence of the HO40 ALT alleles. B) ALT3 and ALT4 arrangements on chr7 of EH23a and EH23b C) Protein based neighbor joining phylogeny showing relationships between the three ALT3 orthogroup OG2876 members on chromosome 7, including the three alternative splice variants (t1, t2, t3) from the EH23a gene model, with the proportion of 100 bootstrap replicates shown on branches where values are > 0.50. D) BKR 7-exon and 11-exon gene models and local nucleic acid alignment for EH23a and EH23b, with close up of 2 bp deletion that truncates the 7 exon model. Green arrows = gene models, yellow arrows = CDS, Red, blue, white, and olive arrows = TEs. Green vertical bars represent percent identity for the alignment. E) BKR protein based neighbor joining phylogeny from 772 pangenome gene models, with the proportion of 100 bootstrap replicates shown on branches where values are > 0.25.

Drug-type populations in North America are thought to have originated from regions of south east and central Asia, and were brought to the western hemisphere via the Caribbean and South America; however, most of what is known about these ancestral populations is based on limited historical accounts and speculation^32^. Earlier demographic models in which hemp-type samples from Europe and Asia were treated as a single population also proposed gene flow events between hemp and drug-type samples ^4^. However, our results reinforce the genomic distinction between European and Asian hemp despite broad phenotypic similarities ^33^.

Importantly, the deep k-mer-based divergence between the single available wild Tibetan assembly ^16^ (Fig. 2c; Supplemental Fig. 6) and all other domesticated and feral lines supports that wild *Cannabis sativa* relatives still exist in remote regions of Asia, despite other reports unable to find support for this idea ^4^. Refining hypotheses regarding domestication, biogeography and the history of various usetypes will require further work with wider samplings of Asian and historical samples and careful consideration of model and polymorphism selection.

The population structure and hybridization models suggest there are regions of the genome driving these differences among populations. We employed a technique called Local Principal Component Analysis (PCA) to look at relatedness along the chromosomes using phased SNPs (Fig. 2d)^34^, which shows close relatedness of many regions across chromosomes 2, 5, 7, and 9 (Fig. 2e, green MDS2 outliers). Furthermore, there are separate regions of distinct relatedness across chromosome 2 and the pseudoautosomal region (PAR) of the X chromosome (shared with the Y chromosome, Fig. 2e, orange MDS1 outliers). A PCA analysis of the green MDS2 outlier windows by population type shows a tight cluster of European hemp and North American feral samples, with a large spread of MJ and other samples (Fig. 2e). In this analysis of genomic regions identified with the local PCA method, the Asian hemp samples appear more closely related to some of the MJ and hc hemp samples than to the European hemp samples. This suggests a distinct ancestry of these regions was woven into the modern pangenome through a complex history of hybridization and selection.

Many plant species have recent whole genome duplications (WGD) that can drive speciation and the evolution of genetic novelty. In cannabis, our Ka/Ks analysis of syntenic orthologs with *Amborella* and *Vitis* imply extant cannabis lineages have not experienced a WGD since the lambda event ∼100 million years ago (mya) (Supplemental Fig. 8), which is consistent with the lack of subgenome dominance found in our F1 haplotype expression analyses and gene copy number variation (Fig. 1, Supplemental Table 2, Supplemental Fig. 4). While cannabis is typically considered diploid ^20^, polyploid (triploid) cultivars have been reported ^35^; however, these appear to be uncommon in North American and European gene pools. We found all 78 haplotype-resolved chromosome-scale assemblies were diploids (2n = 20), similar to prior karyotyping work on European hemp ^36^. Although one report of a natural population of tetraploids was described from a cold high altitude desert in the Himalayas ^37^, it appears that high levels of genetic diversity in modern cannabis (Fig. 2a) have evolved without the occurrence of recent WGDs or hybridization driven alloploidy.

### Transposable element evolution and function

Transposable elements (TEs) contribute to genome evolution, size, and diversity across all domains of life ^38^. While TEs tend to be methylated and epigenetically silenced ^39,40^, some TEs in plants are involved in regulation of gene expression responses to environmental stress ^41^ and the evolution of important phenotypes ^42–45^. Long terminal repeat retrotransposons (LTR-RTs) are found at a moderate to high frequency across the genome (Supplemental Table 4), and specific types of DNA TEs are prevalent near genes, including mutator and helitron, with TE abundance highest within 500 bp upstream of predicted coding genes (Supplemental Fig. 9).

Overall, genes tend to be located near TEs, with an average distance of 443-613 bp (Supplemental Table 5). Methylation is higher in TEs than across the genome on average among the subset of genomes we assessed, which likely suppresses most TE activity. Intriguingly, the MJ samples have consistently higher average TE methylation than European hemp and F1 samples (Supplemental Table 6, Supplemental Fig. 10).

Recent and lineage specific DNA TE expansions suggest ongoing diversification could be driven by hybridization and sources of stress, including clonal propagation, which is a common practice for modern MJ production, but not often used in hemp field production (Fig. 3a). Combined with the recent estimated divergence date (Fig. 3b), these notable patterns suggest that TEs are involved in differentiation of regulatory networks and gene evolution between populations and perhaps specific lineages. For example, in the EH23 hybrid, expressed TEs from our RNA dataset include CACTA, Harbinger, and hAT, in addition to LTR-RTs (Supplemental Fig. 11), pointing to transposition as an active and ongoing feature in at least some cannabis populations and hybrids.

Despite around 4 million years of sustained activity and a recent burst of LTR proliferation (Fig. 3b,c), the cannabis genome has maintained a smaller haploid genome size (∼820 Mb), than its sister genus, *Humulus*, with genome sizes of 1.7 Gb (*Humulus japonicus*) and 2.7 Gb (*Humulus lupulus*, “hop”) ^46^. *Humulus* experienced a large-scale duplication event less than 5 mya, which could explain its larger size ^47^. Additionally, a high solo-intact LTR-RT ratio in cannabis may also help to maintain its smaller genome size ^48^. Solo-LTRs can be created as a result of ectopic recombination that occurs in the internal sequence between two LTR-RTs, which are part of a complete LTR-RT element ^49^. We found that Ty1-*copia* LTR-RTs in particular occur in a high solo:intact ratio, meaning these are being purged from the genome, a finding similar to that in other plants ^50–52^ (Fig. 3d,e). Specifically, the Y chromosome, which shares ∼20 Mb of PAR with the X chromosome, is the longest chromosome and displays the highest level of solo:intact ratio compared to the other chromosomes suggesting it is actively being shaped by TE activity (Fig. 3d; Supplemental Fig. 3, 12, 13).

While DNA TEs make up a small overall percentage of the genome (Supplemental Table 4), the distribution of TEs across the pangenome is highly variable. TEs typically accumulate in regions of infrequent recombination and high levels of cytosine methylation (and CpG content), especially in centromeres and pericentromeric regions (Supplemental Fig. 3 and 14). We find Harbinger TE abundance is especially high in the centromeric region of chromosome 7 (Fig. 3 F), but there is no trace of this peak in the homologous chromosome in hop (Supplemental Fig. 15). The divergence date of this Harbinger block is ∼20 mya (Fig. 3f, Supplemental Fig. 16), corresponding to the approximate time of speciation from *Humulus* ^47,53,54^. This implies the cannabis-hop speciation event involved divergent centromeric evolution and genome rearrangements mediated by Harbinger TE proliferation (Supplemental Fig. 14-16).

### High levels of structural variation

Small and medium length translocations, duplications, and inversions are often directly caused by TE activity, while larger inversions are formed at breakpoints that tend to be enriched for segmental duplications and inverted repeats, which are often signatures of prior TE proliferation (Supplemental Fig. 17) ^55^. SV counts among populations exhibit the highest variation for translocations and duplications, which parallels the high variation in DNA TE abundances across populations (Fig. 3a), while inversion counts have the least variance (Fig. 4a). However, inversion counts do not take into consideration length, which ranges from 200 bp to 25 Mb, in a multimodal distribution (Supplemental Fig. 17), suggesting multiple evolutionary processes act on inversions of differing lengths ^56^. Overall, cannabis inversion statistics such as counts and percent of genome exhibiting inversion are as high as were found in some multiple species comparisons (Supplemental Fig. 17), although sequencing technology likely limits cross-study comparisons ^57^.

Long inversions, like one found on chromosome 1 (19.5 Mb; Fig. 4b), may function as a supergene, perhaps maintained as a balanced polymorphism through associative overdominance, resulting in length variability (Supplemental Fig. 4). Indeed, the 17 instances of this inversion were found to be heterozygous in 15 samples and homozygous in one. This inverted region contains 1,203 genes, spanning many functions, including the core circadian and flowering time gene *PSEUDO RESPONSE REGULATOR 3* (*PRR3*) implicated in the DN behavior of “autoflower” cannabis ^21,26,58^, but are significantly enriched for catalytic activity, transport activity, and binding gene ontology (GO) categories. Pairwise SNP r^2^ values and local PCA plots of this area suggest some level of haplotype formation and elevated linkage disequilibrium (LD) across this region, especially at the interior breakpoint (Supplemental Fig. 18). However, these are not obvious signals of complete differentiation or recombination suppression as has been shown in other systems, such as between ecologically differentiated sunflower species ^59^. We also identified a large ∼25 Mb inversion on the non-PAR region of the Y chromosome in the hc hemp sample Golden Redwood (GRMa; type III) that may provide some insight into sex chromosome evolution (Supplemental Fig. 12). Elucidating the exact function and history of large inverted regions in cannabis requires more targeted investigation, but poses potential challenges for future cultivar improvement projects. Collectively, across the haplotype-resolved, chromosome-scale assemblies, LD decay plots show decay to half of the maximum r^2^ value around 10 Kb (Fig. 4c), which is similar to wild outcrossing soy and rice populations ^60^. However, the elevated long range (100s of Kb to Mb) LD patterns found in certain SNP pairs further underscores the importance of using accurately phased genome assemblies and consideration of SVs for mapping and improvement efforts (Fig. 4d).

The relationship among SVs and TEs is complex. Several hot spots are found on chromosomes 1, 4, and 7, where duplications and translocations are enriched within or near common inversion breakpoints (Fig. 4b). Additionally, some of these regions are associated with specific TE abundances (Fig. 3f). A pangenome wide examination of TE content within SV breakpoints (500 bp window flanking upstream and downstream of the breakpoint [1 Kb total]) shows population specific patterns of enrichment. For example, duplications in MJ samples that contain the same TE at both breakpoints are enriched for Ty3 LTR-RTs (and three classes of DNA TEs), consistent with LTR-RT replication occurring via copy-and-paste mechanism (Supplemental Table 7; *p* < 0.05, Welch’s two-sided *t*-test). Yet across the other five populations, only Harbinger and Mutator DNA TEs are significantly enriched at duplication breakpoints. In some cases, such as feral hemp duplications, no significant TE enrichment of any type was detected; this could reflect the recency of TE activity, or perhaps other mechanisms that give rise to SVs.

Considering lineage and population specific TE and SV patterns, and the frequent occurrence of TEs near genes, suggests a host of mechanisms by which cannabis has diversified and adapted, which were largely cryptic using prior resequencing approaches.

### Pan-gene analyses

In addition to population-level and chromosome-level pangenome analyses, we performed gene-level analyses for cannabinoid synthases, terpene synthases, and disease resistance gene analogs. We developed a web portal that provides Orthofinder based search and extraction functions to make consistent gene based analyses readily available to the research community (Data Availability). Protein coding regions were predicted based on homology and expression data (Supplementary methods; Supplemental Table 8), resulting in 35,000 genes on average for the scaffolded genomes and a linear increase of predicted genes with the duplication status of the contig assemblies (Supplemental Fig. 4). We assessed the quality of the protein coding gene predictions with Benchmarking Universal Single-Copy Orthologue (BUSCO) ^28^, which were around 95% complete on average, matching closely with genome BUSCO scores that averaged above 97% as well (Supplemental Table 1; Supplemental Fig. 4).

### Terpene synthase genes

Terpenes are involved in a complex interplay of communication and defense against herbivores ^61^, as well as potentially modulating cannabinoid activity ^62^. Terpenes in cannabis have been investigated previously ^63–68^, although the genomic organization and extent of variation of terpene synthases have not yet been fully mapped. Chromosomes 5 and 6 have terpene synthase “hotspots,” where the majority of copies are located in a multi-megabase region on one end of the chromosome (Supplemental Fig. 19a,b and Data Availability); this is consistent with previous findings that chromosomes 5 and 6 have QTLs associated with consumer preference for “sweet” or “earthy” aroma profiles ^64^. The majority of terpene synthase genes have 7 exons, consistent with monoterpene or sesquiterpene biosynthesis ^65^.

### Disease resistance gene analogs

In addition to primary gene annotations, we developed plant disease resistance gene analog (“R gene”) maps for the 78 scaffolded, haplotype-phased genomes using Drago2 (Data Availability). Specific R genes encode protein functional domains that determine pathogen specificity and subcellular localization. In cannabis, the *MILDEW LOCUS O* (*MLO*) disease resistance gene family has received the most attention because of the widespread presence of powdery mildew disease caused by *Golvinomyces* spp. in commercial operations ^69^. However, as the legal cannabis industry has expanded, new disease and pest challenges have emerged, including fungal pathogens such as *Fusarium spp*., *Botrytis cinerea* and *Pythium* spp. ^70^. Therefore, knowledge of the diversity of disease resistance genes and alleles has relevance to cannabis breeding. Receptor-like kinases (RLK) are the most abundant R-gene type and overall, R-genes are distributed across all chromosomes. The highest concentration and variation of R-genes are found in hotspots at the top of chromosome 1 (Supplemental Fig. 19c). Most Coiled-Coil NBS-LRR genes (CNLs) are found on chromosomes 3 and 6. Together, these results indicate genomic regions where specific disease resistance breeding targets could be developed (Supplemental Fig. 19d,e).

### Partially domesticated cannabinoid pathway

While key proteins in the cannabinoid pathway have been identified (Fig. 5a, Supplemental Fig. 20), the genomic locations and organization were not resolved until the release of chromosome resolved assemblies ^14,19^. Cannabinoid synthases duplicated and neofunctionalized from the BBE-like family of genes that are expressed on chromosome 7 (Supplemental Fig. 21), then were ultimately reduced to a limited set of functional THCAS and CBDAS alleles through the domestication process (Fig. 5b,c) ^14,71^. The presence of numerous pseudogenized paralogs that are only found in a limited number of cassette arrangements (Fig. 5c, Supplemental Fig. 22) suggests selection has linked a small range of functional alleles to pseudogene cassette haplotypes. These THCAS and CBDAS haplotypes are often, but not always, located in non-syntenic regions of chromosome 7 (Fig. 1a and 5b; Supplemental Fig. 23).

Conversely, the CBCAS paralogs, which are typically located 15-20 Mb closer to the chromosome 7 centromere, appear not to have been under strong selection, with most genomes hosting only pseudogenes (Fig. 5d). Perhaps human preference for THC and CBD, or low expression driven by a weak or disrupted promoter motif, has resulted in limited evidence for CBCAS function. While it was shown that CBCAS genes can produce CBCA *in vivo* when transformed into yeast ^19^, an analysis of over 59,000 cannabis samples found effectively no evidence for CBCA content, although limited commercial testing for CBCA may have led to an underestimation ^72^. Helitron DNA TEs and LTR-RTs are significantly enriched upstream of THCAS, relative to TE content in the full set of genes (Supplemental Table 9), which may underlie gene duplication and evolution of promoters, similar to maize and other plants ^73^. Since helitron and LTR-RT sequences upstream of the cannabinoid synthases cluster largely according to their corresponding synthase type ^14^, but show more diversity than synthase genes (Fig. 5e, Supplemental Fig. 24), improvements to the CBCAS specific promoters could unlock a novel CBCA dominant chemotype.

### Varin cannabinoids and fatty acid genes

*In planta* cannabinoid alkyl side-chain length can vary from one to at least seven carbons, with five carbons being the most common in modern gene pools ^74,75^. Three carbon side-chain cannabinoids (propyl; THCV, CBDV, CBGV) are much less common, but have attracted interest as novel therapeutics ^76,77^. Prior studies have characterized the polygenic nature of this trait, and associated the *β-keto acyl carrier protein (ACP) reductase* (*BKR*) gene with varin cannabinoid production, but left open at least one step needed for a complete biosynthetic hypothesis^78^. In this work, we extend the model for varin cannabinoid production by identifying a complex of *acyl-lipid thioesterase* (*ALT3* and *ALT4)* genes located near the beginning of chromosome 7 that are associated with varin production in our F2 mapping population (Supplemental Fig. 25 and Supplemental Tables 10, 11), and are contained within a common haplotype in our breeding crossover analysis (Fig. 6a,b). *ALT* gene copy number variation is striking in cannabis, ranging from 2–14 copies (considering both phased and unphased assemblies) found across 4 chromosomes (Fig. 5 a). Most plant genomes contain 4–5 *ALT* homologues, and some only a single homologue (e.g. *Brassica rapa, Glycine max*) ^2^. Additionally, *ALT* protein sequence variation in cannabis is notable, with distinct orthogroup membership of each *ALT4* in EH23a and EH23b genomes (Fig. 6b,c), despite these genes being located at similar positions. Since the shortest known fatty acid product of a plant fatty acyl-thioesterase is a 6:0 fatty acid generated by the *Arabidopsis ALT4*, the EH23a *ALT4* allele is a lead candidate for further experimentation. However, given the crossover locations (Fig. 6 a), potential for linkage disequilibrium, and short read mapping issues in this region, any of these *ALT3* and *ALT4* genes (or splice variants) could be causal for varin cannabinoid production. Alternatively, they may have overlapping sub-functionalized substrate specificities, which would pose challenges for further mapping and improvement efforts ^79^.

Although the *BKR* gene on chromosome 4 has been identified previously in a genome wide association study (GWAS), our pangenome findings suggest that a 2-bp deletion produces a 6-exon loss-of-function gene model, which lacks catalytic active site residues (Fig. 6d). Thus, reduction or loss of function in this gene is likely required to increase the butyryl-ACP pool, which one of the *ALT3* or *ALT4* gene enzymes then hydrolyzes to butyric-acid, leading to varin cannabinoid biosynthesis (Fig. 5a). Since cannabis hosts *BKR* genes on both chromosomes 3 and 4, loss of catalytic function of one copy likely does not fully terminate iterative fatty acid chain synthesis. This concept could also explain why varin cannabinoids are only found in certain ratios with pentyl cannabinoids (Supplemental Fig. 25)^75,78^.

Across the pangenome, the EH23a 6-exon *BKR* variant was exclusively found in HO40 pedigree samples (high varin); all other samples, except one 8-exon version of *BKR* in the seed oil cultivar Finola (a low varin producer) are 11 or 12 exon models. The phylogenetic relationships of the predicted *BKR* proteins shows the 6-exon gene may be closer to certain Asian hemp, European hemp and feral variants (Fig. 6e). However, one of the 11-exon gene clades contains the varin producing AutoCBDV genome, and potential varin producer Durban Poison, which could be reduced function variants. Some reports of the varin chemical phenotype suggest that there is no defined geographic origin associated with this trait ^80^.

However, other studies report plants containing high amounts of varin cannabinoids from the southern regions of Africa and certain regions of Asia ^75,81^. Since this is a polygenic trait with genes on different chromosomes it is not possible to fully assess the origins and range of each potential gene involved. Nonetheless, the *BKR* gene phylogeny, and whole genome k-mer based clustering analysis suggest an Asian origin for varin cannabinoid genes used in this breeding project (Fig. 2c).

## Conclusions

After conducting this significant pangenome analysis of 193 genome assemblies, we conclude that the cannabis gene pool still remains under sampled, and Asian germplasm is underrepresented in both genome and germplasm resources. Additionally, extant wild cannabis relatives likely remain in Asia, but little is known about their range or conservation status, which leaves questions about taxonomy and population structure unresolved in the *Cannabis* genus. Asian hemp, which generally resembles European hemp phenotypically, has many divergent genomic regions, some of which are more closely related to North American drug-type samples. We find that lineage and population specific TE activity, in conjunction with hybridization, drives genomic evolution in cannabis rather than whole genome duplication and ploidy variation.

Furthermore, SVs pose challenges to improvement efforts, and contain diversity that was previously cryptic using short read resequencing techniques. Despite a century of prohibition and limited germplasm conservation efforts, overall genetic diversity in cannabis remains high. Although the well-studied cannabinoid synthase genes have low levels of sequence variation, other classes such as disease resistance, terpenes, growth regulators, and fatty acid biosynthesis genes contain high levels of sequence diversity and copy number variation. We hope that the development and release of 78 chromosome-level, haplotype-resolved assemblies, including the phased EH23 used to anchor many of the analyses presented here, will pave the way for accelerated conservation and improvement efforts. Additionally, we are currently developing a cannabinoid-free, autoflower (DN), and monecious model cannabis plant to stimulate research activities.

In this work we have expanded previous genetic and inheritance models of varin cannabinoid production using the first chromosome-scale haplotype-resolved cannabis genomes. While cannabinoids will continue to provide opportunities for drug development, our findings of high levels of fatty acid biosynthesis gene variation (*ALT*, *BKR*, and others) suggest the *Cannabis* genus harbors untapped lipid metabolism traits. Given the partial overlap between cannabinoid biosynthesis and seed oil anabolism, hybridizing diverse parental lines outside of the typical Northern European hemp seed oil gene pool could produce novel lipid profiles and characteristics. In particular, utilization and conservation of Asian hemp and Asian wild cannabis remain an important next step toward normalizing and modernizing cannabis science and agriculture.

## Conflict of Interest

S.C. was a co-founder of Oregon CBD. A.R.G and A.T. were employees of Oregon CBD. R.C.L is a stakeholder in Saint Vrain Research LLC, which manufactures hemp-based products. T.P.M is a founder of the carbon sequestration company CQuesta.

## Author Contributions

T.P.M., R.C.L., S.C., A.R.G., and K.V. conceived and organized research efforts. R.C.L., L.K.P.-C., T.P.M., B.J.K., and K.V. wrote and edited the manuscript. R.C.L., L.K.P.-C., T.P.M., B.J.K., N.T.H., N.A., A.A., A.M., J.K., A.R.G., A.T., and K.W. analyzed data. R.C.L., L.K.P.-C., A.R.G., K.C., E.M., T.D., and S.C. conducted greenhouse, field and laboratory experiments.

## Acknowledgments

We thank the entire Michael Lab for discussion on this work. We thank Tyler Gordon and Zach Stansell from the USDA for sending leaf material from lines from the GRIN collection. This work was funded in part by a National Science Foundation Plant Genome Postdoctoral Research Fellowship to L.K.P.-C. (NSF-IOS PRFB 2209290), the Tang genomics fund to T.P.M., and the pangenome tools development in the Michael Lab was supported by Bill and Melinda Gates Foundation (INV-040541) to T.P.M.

## Data Availability

The complete set of pangenome assemblies, gene and TE annotations, DNA read libraries (PacBio and Illumina), and RNA reads are available from NCBI (pending). A subset of 44 genomes and annotations, as well as links to corresponding U.S. National Plant Germplasm System accessions are available from: <u>resources.michael.salk.edu</u>. Methylation data, and specific annotations for R-genes, terpenes and cannabinoid synthases and additional genome visualizations are available from: <u>figshare.com/projects/Cannabis_Pangenome/205555</u>. Orthobrowser and Genome Jbrowse instances are hosted at: <u>resources.michael.salk.edu/root/home.html</u>.

## Methods

### Plant material

*Cannabis sativa* pangenome samples were selected from multiple sources to maximize the genetic diversity, history and agronomic value. A large portion of the pangenome comes from the Oregon CBD (OCBD) breeding program that includes elite cultivars; foundational marjuana lines potentially originating from the 1970s, 80s, 90s to present; and elite trios used for different aspects of the breeding program. The remaining cultivars come from the United States Department of Agriculture (USDA) Germplasm Resource Information Network (GRIN) and German federal genebank (IPK Gatersleben) repositories, as well as collections made by the Salk Institute from various breeders. The pangenome includes European and Asian fiber and seed hemp, feral populations, North American marijuana (type I), and North American high cannabinoid yielding (CBD or CBG) hemp (type III and IV). Additional cannabinoid diversity is represented with chemotypes presenting high expression of pentyl or propyl (varin) homologs of CBD or THC, and cannabinoid free (type V) plants. Flowering time variation is also captured with the inclusion of both regular short day and day neutral (autoflowering) phenotypes (Supplemental Table 1).

### EH23 phased, haplotype-resolved, chromosome-scale anchor genome

EH23a (HO40) and EH23b (ERB) are haplotype-resolved assemblies for ‘ERBxHO40_23,’ an F1 resulting from a cross between parents, ERB and HO40, both being proprietary female inbred lines from OCBD. ERB is a day-neutral (DN; autoflower), type III (CBDA-dominant) plant and HO40 is a high-varin (propyl), type I (THCA-dominant) plant. The genetically female (XX) ERB plant was induced to produce male flowers by treatment with silver thiosulfate (STS) and used to pollinate HO40. One individual from the F1 populations (ERBxHO40_23) was selected for genome sequencing. Initial genome size estimates of ‘ERB x HO40_23’ using flow cytometry estimated a diploid genome size of 1445.6 Mbp (722.8 Mbp haploid genome size). High molecular weight (HMW) DNA was extracted from leaf tissue. Following DNA extraction and library preparation (see HMW DNA Isolation and Genome sequencing section) HiFi reads were generanted on the PacBio Sequel II. Hifiasm v0.16.1^1^ was then used in conjunction with HiC reads to produce initial assemblies. After assembly, HiC reads were aligned to the Hifiasm_HiC contigs using the Juicer v1.6.2 pipeline^2^ followed by ordering and orientation utilizing version 180922 of the 3D-DNA pipeline^3^. The scaffolded assemblies were then manually corrected using Juicebox v1.11.08^4^.

### EH23 F2 population

In addition to the whole genome sequencing data described above, ERBxHO40_23 was self-pollinated using STS induced masculinization of select flowers, to create an F2 mapping population. From this F2 population, individuals were scored for autoflower, varin content, and sequenced using Illumina 100 bp reads by NRGene (Nrgene Technologies Ltd, Israel). Illumina WGS genotyping runs were performed on 288 plants from this population, plus the ERBxHO40_23 parent. Trim_galore was used to trim sequences using: --2 colour 20, resulting in 271 individuals for analysis ^5^. On average samples had 8.5x coverage. Minimap was used to align each sample to EH23b.softmasked.fasta. Freebayes was used to call variants: -g 4500 -0-n 4 --trim-complex-tail --min-alternate-count 3^6^. Bcftools was used to filter on QUAL>20 scores (99% chance variant exists) ^7^. Finally, Vcftools^8^ tools was then used to further filter SNPs: --remove-indels --minGQ 20 --maf 0.25 --max-missing 1 --min-alleles 2 --max-alleles 2 –stdout --recode^8^; only sites that were scored as heterozygous (0/1) in ERBxHO40_23 sample were retained, resulting in 93,251 SNPs.

### EH23 F2 cannabinoid HPLC methods

High performance liquid chromatography (HPLC) was conducted according to the protocol thoroughly described in Garfinkel et al. 2021^9^ to determine relative propyl and pentyl cannabinoid content in all the plants used in this study, including F2 progeny. In short, mature flower tissue was collected from each individual, frozen at −80 C and homogenized, before cannabinoids were extracted in methanol.

### EH23 RNASeq

Total RNA extraction was done using the QIAGEN RNeasy Plus Kit following manufacturer protocols. Total RNA was quantified using Qubit RNA Assay and TapeStation 4200. Prior to library prep, we performed DNase treatment followed by AMPure bead clean up and QIAGEN FastSelect HMR rRNA depletion. Library preparation was done with the NEBNext Ultra II RNA Library Prep Kit following manufacturer protocols. Then these libraries were run on the NovaSeq6000 platform in 2 x 150 bp configuration.

### EH23 haplotype expression analysis

We measured gene expression levels using Salmon v1.6.0^10^. In brief, the raw paired end short-reads from sequencing were mapped to the coding sequences (CDS) from both haplotypes (EH23a and EH23b), and the abundance was estimated in transcripts per million (TPM) for downstream analysis. Mapping rates were calculated with samtools flagstat ^7^. The minimum transcript per million (TPM) threshold for a given gene was >= 0.1. Visualization was performed using a combination of Matplotlib ^11^, SciPy ^12^, and NumPy ^13^, and expression values were shown in heatmaps as log_2_ TPM to represent log-fold change.

### HiC library preparation and sequencing

For the Dovetail Omni-C library, chromatin was fixed in place with formaldehyde in the nucleus and then extracted. Fixed chromatin was digested with DNAse I, chromatin ends were repaired and ligated to a biotinylated bridge adapter followed by proximity ligation of adapter containing ends. After proximity ligation, crosslinks were reversed and the DNA purified. Purified DNA was treated to remove biotin that was not internal to ligated fragments. Sequencing libraries were generated using NEBNext Ultra enzymes and Illumina-compatible adapters. Biotin-containing fragments were isolated using streptavidin beads before PCR enrichment of each library. The library was sequenced on an Illumina HiSeqX platform to produce ∼ 30x sequence coverage. Then HiRise used (See read-pair above) MQ>50 reads for scaffolding. Additional HiC libraries were generated using Phase Genomics Proximo HiC Kit (Plant) version 4.

### High Molecular Weight DNA Isolation and Genome sequencing

All samples were sequenced on a Pacific Biosciences Sequel II. For samples sourced from “Michael” (Supplemental Table 1), HMW DNA was isolated using Carlson Lysis buffer and Qiagen Genomic tips as described in Oxford Nanopore Technologies (ONT) Protocol “Plant leaf gDNA” Arabidopsis method. The DNA was further size-selected for fragments longer than 10-25 kb using the ONT Short Fragment Eliminator Kit (EXP-SFE001). HMW DNA was then confirmed by Tapestation Genomic DNA ScreenTape (Agilent cat# 5067-5365) or Femto Pulse Genomic DNA 165 kb Kit (Agilent cat# FP-1002-0275). For samples sourced from “OCBD” (Supplemental Table 1) HMW DNA was isolated using a modified protocol ^14^. Briefly, samples were ground in a mortar and pestle with liquid nitrogen, two chloroform:isoamyl wash cycles were performed, and ‘Total Pure NGS’ beads from Omega Biotek were used as a substitute from the original protocol. gDNA quality/purity was then assessed using a NanoDrop One (ThermoFisher) prior to starting library preparation. CLR libraries were made using the Pacbio protocol PN 101-693-800 V1.

Size selections on gDNA were made using the Blue Pippin U1 High Pass 30-40 kb cassette with a 30-40 kb base pair starting threshold to produce fragment distributions of 60-90 kb. HiFi CCS libraries were prepared according to the PacBio protocol (PN 101-853-100 V5). Sheared gDNA fragment distributions with a modal peak ∼18kb were produced using g-Tubes from Covaris and Blue Pippin S1 High Pass 6-10kb cassettes to remove everything under 10kb in size.

### Pangenome assembly and scaffolding

All genomes labeled Hifiasm_HiC, Hifiasm_Trio_RagTag, Hifiasm_RagTag, and Hifiasm (Supplementary Table 1) were assembled using Hifiasm v0.16.1^1^. When available, HiC data and HiFi parental trio data were also incorporated into the assembly process defining the Hifiasm_HiC and Hifiasm_Trio_RagTag types respectively. CLR (continuous long reads) assemblies were generated using FALCON Unzip from PacBio SMRT Tools 9.0 Suite ^15^ and CCS (circular consensus sequencing) labeled genomes were assembled with HiCanu v2.2 ^16^. After assembly, HiC reads were aligned to the Hifiasm_HiC contigs using the Juicer v1.6.2 pipeline^2^ followed by ordering and orientation utilizing version 180922 of the 3D-DNA pipeline^3^. The scaffolded assemblies were then manually corrected using Juicebox v1.11.08^4^.

Hifiasm_RagTag and Hifiasm_Trio_RagTag assemblies were scaffolded using the split chromosomes of the 24 HiC scaffolded genomes and error checked with yak-0.1 (github.com/lh3/yak). Sourmash v4.6.1^17^ was used to generate a Jaccard similarity matrix between the chromosomes and each un-scaffolded assembly, and the most similar version of chromosome 1 through X was concatenated to generate a reference for scaffolding via RagTag v2.1.0^18^. If the similarity matrix identified the Y chromosome as the best match, the assembly remained un-scaffolded. BUSCO v5.4.3 ^21^ with the eudicots_odb10 dataset and assembly-stats v1.0.1 (https://github.com/sanger-pathogens/assembly-stats) were used on all assemblies to measure completeness and contiguity.

### Basecalling methylated cytosines

Genomic reads from the raw Oxford Nanopore (ONT) FAST5s generated from *C. sativa* deep sequencing samples were used for methylation calling. Genome assemblies generated for the same individuals were used as references for alignment. FAST5 data were converted to POD5 format using the pod5 software package (https://github.com/nanoporetech/pod5-file-format).

Methylation calling was performed with ONT base calling software Dorado version 0.3.4 (https://github.com/nanoporetech/dorado/). Dorado uses the raw POD5 data and a reference to identify methylated cytosines. This was performed with the super high accuracy (SUP) base calling model trained for R9.4.1 or R10.4.1 pore type and 400bps translocation speed, according to the sequencing conditions for each line. The assembled genomes generated from each sample were used as references to generate an aligned BAM file with MM/ML tags containing 5mC and 5hmC methylation calls. These were then piled up with modkit (https://github.com/nanoporetech/modkit), and the piled-up calls (aggregating 5mC with 5hmC) were used for calculating genome-wide methylation frequencies across all CG sites.

### Gene and repeat prediction

Gene model prediction involved a multi step pipeline and was applied to all assemblies.

1. Curation of a repeat library using RepeatModeler ^19^ on a small number of high quality cannabis assemblies and pre-existing repeat libraries. We used OrthoFinder (v2.5.4) ^20^ to group repeats for deduplication. The final repeat library included 10% of the sequences from each orthogroup (minimum 1) for a total of 6262 sequences from 5793 groups.

a. Finola (GCA_003417725.2)
b. CBDRx (GCF_900626175.2)
c. Purple_Kush (GCA_000230575.5)
d. ERBxHO40_23
e. ERBxHO40_23
f. I3
g. JL (GCA_013030365.1)
h. ERB_F3
i. Cannbio-2 (GCA_016165845.1)
j. W103
k. JL_Mother (GCA_012923435.1)
l. FB30
m. TS1_3_v1
n. HO40
2. Mask repeats identified in our library with RepeatMasker (v4.1.2) ^21^.
3. Gene prediction with the TSEBRA pipeline (using Braker v2.1.6) ^22^ using a variety of pre-existing protein libraries from cannabis and other organisms:

a. *Arabidopsis thaliana*
b. *Theobroma cacao*
c. *Glycine max*
d. *Rhamnella rubrinervis*
e. *Ziziphus jujuba*
f. *Trema orientale*
g. *Vitis vinifera*
h. *Prunus persica*
i. *Morus notabilis*
j. *Cannabis sativa*
k. *Humulus lupulus*
4. RNAseq libraries (Supplemental Table 8) were aligned with either hisat2 (v2.2.1) ^23^ for short read mapping, or minimap2 (v2.24) ^24^ for full-length cDNA.

i. Short-read Illumina data trimmed with fastp ^25^.
5. Putative functional annotations of genes using eggnog-mapper (v2.0.1) ^26^.
6. Validation of gene models by comparing genome BUSCO (v5.4.3) ^27^ scores to proteome BUSCO scores on the eudicots_ocdb10 dataset (Supplemental Table 1).

### Ideogram methods (Figure 1)

Ideograms for each pair of chromosomes were created using ggplot2 [https://ggplot2.tidyverse.org] in R (www.R-project.org). The length of each chromosome was determined using ‘nuccomp.py’ (github.com/knausb/nuccomp) and used with ‘ggplot::geom_rect()’ to initialize the plot. One million base pair windows were created for each chromosome where the number of ‘CpG’ motifs were counted for each window with the program ‘motif_counter.py’ (github.com/knausb/nuccomp). The CpG count was converted into a rate by dividing by the window size; this also accommodated the last window of each chromosome which was less than one million base pairs in size. These rates were scaled by subtracting the minimum rate and then dividing by the maximum rate (the maximum rate after subtracting the minimum rate), on a per chromosome basis. In order to visually emphasize the enrichment of the ‘CpG’ motif in the centromeric region, an inverse of the CpG rate was taken by taking one and subtracting the ‘CpG’ rate for each window. This scaled, inverse ‘CpG’ rate was used for the width of each one mbp window and colored based on gene density using the viridis magma palette (doi.org/10.5281/zenodo.4679424).

Structural variation among each pair of chromosomes was determined using minimap2 ^24^ alignments. The minimap2 comparisons were annotated using SyRI ^28^. The syntenous and inverted regions were plotted using ggplot2::geom_polygon() in a manner inspired by plotsr ^29^ but implemented in R (github.com/ViningLab/CannabisPangenome).

The location of candidate loci within EH23 haplotypes A and B were determined using BLASTN^30^. Query sequences were as follows: cannabichromenic acid synthase (LY658671.1), cannabidiolic acid synthase (AB292682, AB292683, AB292684), tetrahydrocannabinolic acid synthase (AB212829, AB212830), and olivetolic acid cyclase (NC_044376.1:c4279947-4279296, NC_044376.1:c4272107-4271242). These sequences were combined with centromeric, telomeric and rRNA sequences in the file ‘blastn_queries_rrna_cann.fasta’ (github.com/ViningLab/CannabisPangenome). BLASTN was called with the following options: -task megablast -evalue 0.001 -perc_identity 90-qcov_hsp_perc 90. Tabular results (subject chromosome, subject start of alignment, subject end of alignment) from BLASTN were read into R and plotted on ideograms with ggplot2::geom_rect() (https://ggplot2.tidyverse.org).

### Centromere and telomere analysis

ONT and PacBio based long read-based genome assemblies enable the assembly of some of the highly repetitive centromeres and telomeres sequences ^31^. Centromeres were identified by searching genomes using tandem repeat finder (TRF; v4.09) using modified settings (1 1 2 80 5 200 2000 -d -h) ^32^. TR were reformatted, summed and plotted to find the highest copy number TR per our previous methods to identify centromeres (Supplemental Figure 3) ^31^.

Telomeres were estimated using two different methods. First, the TRF output was queried for repeats with the period of seven (7) for the 14 different version of the canonical telomere base repeat: AAACCCT, AACCCTA, ACCCTAA, CCCTAAA, CCTAAAC, CTAAACC, TAAACCC, TTTAGGG, TTAGGGT, TAGGGTT, AGGGTTT, GGGTTTA, GGTTTAG, GTTTAGG. (grep -a ‘PeriodSize=7’ *.genome.fasta.1.1.2.80.5.200.2000.dat.gff | grep -a ‘Consensus=AAACCCT\|Consensus=AACCCTA\|Consensus=ACCCTAA\|Consensus=CCCTAA A\|Consensus=CCTAAAC\|Consensus=CTAAACC\|Consensus=TAAACCC\|Consensus=TTTAG GG\|Consensus=TTAGGGT\|Consensus=TAGGGTT\|Consensus=AGGGTTT\|Consensus=GG GTTTA\|Consensus=GGTTTAG\|Consensus=GTTTAGG’ -). Second, we searched raw ONT and PacBio reads for telomere sequences using our TeloNum algorithm ^33^. While the results were variable across the pangenome assemblies, in general, telomere sequence was found at the end of the chromosome with an average length of 16 kb for PacBio assemblies and 60 kb for ONT assemblies. The differences between ONT and PacBio telomere length most likely reflected the input read length of >100 kb and 15-20 kb respectively. TeloNum analysis of the raw reads supported the distributions from the assemblies consistent with most chromosomes having telomere sequence while being shorter than the actual size. Cannabis telomeres are on the longer side for a eudicot and could be explained by its predominantly clonal propagation for medicinal uses ^34^.

Centromere sequence was identified based on the hypothesis that it will be the most abundant repeat in the genomes that also has a higher order repeat (HOR) structure ^31,35^. Two different repeats with HOR were identified in the PacBio HiFiasm assemblies, while only one was found in the ONT assemblies and the previous CBDRx assembly, which is based on ONT sequence ^36^. The highest copy number repeat was 370 bp that varied between 20-30 Mb (2-4% of the total genome) with HOR at 740 and 1,110 bp (Supplemental Figure 3). The second highest, and the only one found in the ONT assemblies, was a 237 bp repeat that varied between 3-5 Mb (0.4-1.0% of the total genome) and had HOR at 474 and 711 bp (Supplemental Figure 3). Mapping of the 370 bp repeat to the chromosome-resolved genomes revealed that this repeat was primarily located at the end of the chromosomes next to the telomere sequence, which suggested that it may be related to the CS-1 subtelomeric repeat ^37^. Comparison of the putative 370 bp centromeric repeat and the CS-1 subtelomeric repeat showed they are the same repeat element. In contrast, the putative 237 bp centromeric repeat predominantly was found on Chr06 and Chr08 in the predicted centromere region (Figure 1). However, smaller 237 bp arrays were found on all chromosomes across the assemblies in the predicted centromere region (based on CpG, methylation, gene content and TEs) with most assemblies having small arrays on Chr06 and Chr08.

### Ribosomal DNA detection and quantification

Ribosomal DNA (rDNA) 45S (18S, 5.8S, 26S) and 5S sequences were identified in the CBDRx/CS10 assembly (LOC115701787 5.8S, LOC115701759 18S, LOC115701762 26S, and LOC115721558 5S) and used to BLAST against the pangenome assemblies (Figure 1; Supplemental Figure 3). Across the scaffolded genomes the 45S array was predominantly located on the acrocentric end of Chr08, and the 5S was located exclusively on Chr07 between the cannabinoid synthase cassette array, consistent with published results with FISH (fluorescence in situ hybridization) ^37^. However, partial arrays were found in some assemblies on all of the chromosomes (Supplemental Figure 3). The distribution of the partial arrays on different chromosomes could reflect variability across the genomes since some share similar locations across assemblies. Most arrays are found on the un-scaffolded contigs, suggesting that these variable arrays across different chromosomes could be the result of mis-assemblies. In general there are on average 1,000 45S and 2,000 5S arrays in the cannabis genome; some assemblies have the 5S array completely assembled on Chr07.

### Allele frequency methods

Genotype data in the VCF format ^7^ was input into R (www.R-project.org/) using vcfR ^38^. Allele and heterozygote counts were made with vcfR. Wright’s F_IS_ ^39^ was calculated to provide the deviation in heterozygosity from our random, Hardy-Weinberg, expectation. Wright’s FIS was calculated as (HS - HO)/HS where HO is the observed number of heterozygotes divided by their number and HS is the number of heterozygotes we expect based on the allele frequencies, calculated as the frequency of the first allele multiplied by the frequency of the second multiplied by two and divided by their number. Scatterplots were generated using ggplot2 (ggplot2.tidyverse.org). Graphical panels were assembled into a single graphic using ggpubr (<u>CRAN.R-project.org/package=ggpubr</u>).

### PanKmer genome analysis

Using PanKmer, we constructed two 31-mer indexes: a “full” index of 193 Cannabis assemblies and a “scaffolded-only” index of 78 scaffolded assemblies, using the “pankmer index” command with default parameters. We calculated and plotted pairwise Jaccard similarities for all assemblies in the full index using “pankmer adj-matrix” followed by “pankmer clustermap--metric jaccard”. We calculated and plotted a collector’s curves for both the full and scaffolded-only indexes using the “pankmer collect” command with default parameters. Mathematical details of the collector’s curve are provided in Supplementary Note 1. All scripts used for this analysis can be found on GitHub.

### Analysis of gene-based pangenome

We define the gene-based pangenome as the set of all gene families (orthogroups) with a representative in at least one genome of the pangenome. For each of 193 (78 scaffolded) *C. sativa* genomes, the primary transcript of each high-confidence gene prediction was chosen as a representative. The proteins corresponding to each primary transcript were clustered into orthogroups using Orthofinder (v.2.5.4, see Orthofinder and synteny analysis section below).^20^ The set of primary transcript CDS were merged into a single FASTA file, and exact duplicates were removed with SeqKit (2.7.0) ^40^. Among primary transcripts, likely contaminants were determined by identifying transcripts predicted on contigs where fewer than 90% of predictions were annotated as either “viridiplantae” or “eukaryote” according to eggNOG-mapper (v2.1.12) ^26^, and were removed. To mitigate the problem of unannotated genes, we aligned coding sequences of all primary transcripts to each of the 193 (78) Cannabis genomes using minimap2 (v2.26) ^24^ with parameters “minimap2 -c -x splice” to generate a PAF file with CIGAR strings for each genome. For each genome, if an aligned CDS sequence had a mapping quality of at least 60, had a number of CIGAR matches at least 80% of the query length, and did not overlap a directly annotated gene, it was considered an unannotated gene and its orthogroup was marked as present in the target genome. The set of orthogroups that had at least one representative present in all 193 (78) genomes were considered to be the core genome, the remaining orthogroups were considered to be the variable genome. The presence or absence of each orthogroup in each genome was recorded in a table (Data Availability). All scripts for this analysis are available from GitHub.

### Calculating collector’s curves: Haplotypes, orthogroups, and scores

In pangenomics, collector’s curves (pangenome rarefication) show the relationship of the number of haplotypes (here *H*) to the number of gene families or orthogroups (here *X*).

Given the *X* orthogroups distributed across *H* haplotypes, let the score *s_x_* ∈ [0, *H*] of an orthogroup *x* be the number of haplotypes in which *x* is present. For any score s let *P*(*s*) be the number of orthogroups with score equal to *s*.

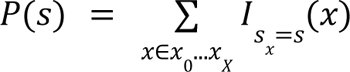

Where *I_s_x_*:{*x*_0_…*x_X_*} → {0,1} is the indicator function on {*x* ∈ *x*_0_…*x_X_*: *s_x_*=*s*}.

### The collector’s curves

The collector’s curve *C*(*h*): [1, *H*] → [0, *X*] is the expected number of orthogroups that will be present in a subset of *h* haplotypes randomly drawn from the total set of *H*. It can be calculated by:

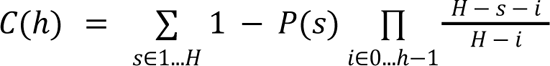

The expected number of *core* orthogroups *Ĉ*(*h*) can be estimated by

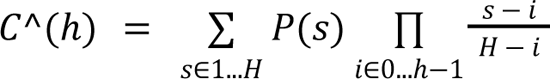

Each of these is a special case of a general formula for the expected number of orthogroups with a score of at least *n*, based on the hypergeometric survival function:

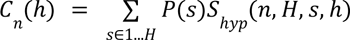

Where *S*_hyp_ is the hypergeometric survival function or the hypergeometric cumulative distribution function subtracted from 1:

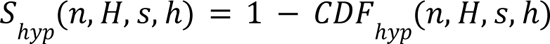

And, conventionally, the cumulative distribution function is:

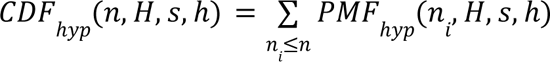

So defined, we can see that the pan-genome collector’s curve *C*(*h*) is equivalent to *C_1_*(*h*), while the core genome collector’s curve *Ĉ*(*h*) is equivalent to *C_h_*(*h*): *k*-mer based collector’s curves

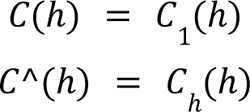

The definition of the collector’s curve is agnostic to the unit of genomic sequence, so the calculation of a *k*-mer based curve is identical to the orthogroup based curve, excepting that *X* will be the number of *k*-mers and *x* will represent a *k*-mer, rather than an orthogroup.

### Repeat analysis

#### Identification of solo-LTRs and intact LTR-retrotransposons (LTR-RTs)

To identify solo-LTRs and intact LTR-retrotransposons (LTR-RTs), we used the EDTA pipeline ^41^. We identified solo-LTRs by first collecting the set of LTRs that were not assigned as intact LTR-RTs, which are retrieved on the basis of “method=homology” in the attribute column of the TEanno.gff3 file. We applied thresholds to isolate solo-LTRs from truncated and intact LTRs, as well as internal sequences of LTR-RTs. These thresholds include a minimum sequence length of 100 bp, 0.8 identity relative to the reference LTR, and a minimum alignment score of 300 ^42^. We also required that the four adjacent LTR-RT annotations did not have the same LTR-RT ID ^43^. Further, we required a minimum distance of 5,000 bp to the nearest adjacent solo-LTR, intact LTR, or internal sequence ^44^. Last, we kept solo-LTR sequences that fell within the 95th percentile for LTR lengths ^45^. Overall, this method represents a modified approach based on the solo_finder.pl script from LTR_retriever ^42^ and the LTR_MINER script ^44^ with guidance from the github page for LTR_retriever (github.com/oushujun/LTR_retriever/issues/41).

#### Enrichment of transposable elements flanking genomic features

The method presented as part of PlanTEnrichment ^46^ was adapted for the *Cannabis* pangenome to assess TE enrichment both upstream and downstream of different genomic features, including cannabinoid synthase genes. The goal of the analysis was to identify TEs that are significantly associated with a specific category of genomic feature. Briefly, “X” represents a specific type of TE and “Y” encompasses all TEs. The total number of X located upstream or downstream of a specific genomic feature (for example, cannabinoid synthases) is denoted as “a”; the total number of X located upstream or downstream of all genomic features (for example, all genes) is “b”; the total number of Y located upstream or downstream of a specific genomic feature (cannabinoid synthases) is “c”; and the total number of Y located upstream or downstream of all genomic features (all genes) is “d”. An enrichment score (ES) is defined as *ES* = (*a*/*b*)/(*c*/*d*), and the p-value is defined as *p* = (*a* + *b*)! (*c* + *d*)! (*a* + *c*)! (*b* + *d*)! /(*a*! *b*! *c*! *d*! *N*!), where N is the sum of a, b, c, and d. A multiple test correction ^47^ was performed on the p-values using the Python library statsmodels ^48^. Significance threshold cut-offs included an FDR < 0.05 and ES >= 2.

We used bedtools intersect ^49^ to collect and survey the set of TEs located 1 kb upstream or downstream of the genomic feature category of interest. An example command: bedtools intersect -a assemblyID_genomic_feature_coord_file.txt -b assemblyID.TE.gff3 -wo > assemblyID_intersect_results.txt.

#### Distance between genes and TEs

The median and mean distances between genes and each of the TE categories was calculated using bedtools sort (`bedtools sort -i genome.TEs.bed > genome.sorted.TEs.bed`) and bedops closest-features (command: `closest-features--closest --header --dist genome.sorted.genes.bed genome.sorted.TEs.bed > genome.closest_features.bed`) ^50^. To obtain the initial pre-sorted BED file for genes, the following command was used: `cat genes.gff3 | grep mRNA | grep ‘.chr’ | awk ‘{print $1“\t”$4“\t”$5“\t”$7“\t”$3“\t”$9}’ > genome.genes.bed`. For TEs, the following command was used: `cat genome.EDTA.TEanno.gff3 | grep ‘.chr’ | awk ‘{print $1“\t”$4“\t”$5“\t”$7“\t”$3“\t”$9}’ > genome.TEs.bed`. To calculate mean and median values, the built-in Python statistics module was used.

#### Phylogeny of TEs surrounding cannabinoid synthases

The genomic coordinates for the 2 kb flanking distance surrounding copies of CBCAS, CBDAS, and THCAS for the 78 scaffolded assemblies were retrieved with bedtools flank (bedtools flank -i assemblyID_synthase_coords.bed -g chromSizes.txt -l 2000 -r 2000 > assemblyID_flanking_2000.bed). Next, the TEs contained in this flanking region were retrieved using bedtools intersect (bedtools intersect -a assemblyID_flanking_2000.bed -b assemblyID.EDTA.TEanno.gff3 -wo > assemblyID_intersect_2000.bed) ^49^. The genomic sequences for each of the TE types identified with bedtools intersect were collected in a fasta file and aligned with mafft (mafft --auto helitron.fasta > helitron_aln.fasta) ^51^. A maximum likelihood tree was constructed with FastTree (FastTree -nt -gtr -gamma helitron_aln.fasta > helitron_aln.tree) ^52^. The tree was visualized with FigTree ^53^. To reduce redundancy in the full set of LTRs, CD-HIT was applied to the set of sequences, prior to multiple sequence alignment (cd-hit-est -i Ty1_LTRs.fasta -o Ty1_LTRs.cdhit.fasta -c 1) ^54^.

#### Expression analysis of active TEs

The non-redundant TE sequence library from EDTA was provided as the “transcriptome” to salmon. Each of the EH23 RNA-seq samples was mapped to the TE transcriptome. Similar to the gene expression analysis, the minimum transcript per million (TPM) threshold for a given TE was >= 0.1 TPM in >= 20% of samples ^55^. The top 50 expressed TEs were visualized as a heatmap, showing log_2_ TPM to represent log-fold change.

#### Observed/expected CpG

“CpG islands” are defined as unmethylated regions spanning >200bp, GC content >50%, and observed/expected CpG ratio >0.6. Cytosine methylation over time results in a loss of CpG dinucleotides after cytosine is deaminated to thymine. With cytosine methylation, the expectation is that CpG dinucleotides (CG, CHG, CHH [where H=A, T, or C]) will have greater methylation activity. The observed/expected CpG ratio calculation is: (*CpG dinucleotide count/L*)/(*C count/L * G count/L*) ^56,57^. Observed/expected CpG patterns were visualized in Figure 3 F.

#### Analysis of transposable elements directly flanking structural variants

For each of the structural variant subtypes (inversions [INVS], duplications [DUPS], translocations [TRANS], and inverted translocations [INVTR]), the flanking region 500 bp upstream and downstream of each breakpoint (1 kb total for each breakpoint) was surveyed for TE content, using both intact and fragmented annotations. To compare with the genome at large, a random window was retrieved from the same genome and chromosome, with the same length as each of the SVs with bedtools shuffle, and the flanking windows were retrieved for each of the simulated breakpoints. Only cases where a specific type of TE was associated with both breakpoints of a single SV were further assessed with bedtools intersect. Both fragmented and intact TEs were included in this analysis. Statistical significance was assessed using Welch’s two-sided t-test in SciPy ^12^. All 78 scaffolded, chromosome-level genomes were included, grouped by population.

TEs occur more frequently near SV breakpoints (500 bp upstream and downstream of the breakpoint; 1 kb total) than in randomly selected regions of the same length from the same chromosome and genome. To overcome differences in abundance, the randomly shuffled regions of the genome were bootstrapped (1,000 replicates), with the requirement that each of the simulated, shuffled TE datasets match the number of observed breakpoints in the population.

The TE content of observed and simulated data was assessed for statistical significance with Welch’s two-sided *t* test in scipy ^12^ and Benjamini/Hochberg multiple test correction (alpha=0.5, method=’indep’, is_sorted=False) ^48^. A test statistic and P value was generated for each of the thousand bootstrap replicates. The average test statistic and P value were then calculated (Supplemental Table 7).

#### Orthofinder and synteny analysis

We ran Orthofinder version 2.5.4 to aid in analysis of the hundreds of cannabis proteomes. Two runs were completed. The first was focused on our highest quality cannabis assemblies and only included scaffolded assemblies along with dozens of other plant samples from Plaza and a few samples from NCBI. Another run, including all of our cannabis pangenome assemblies, along with close relatives sourced from Plaza, was also produced to allow for detailed protein level analysis of the remaining assemblies. In all cases, only the primary (longest isoform unless otherwise annotated) protein sequence was used. Orthofinder results were analyzed using a variety of methods, including an as yet published tool, Orthobrowser, capable of generating static web pages that allow for simultaneous visualization of gene tree dendrograms, gene tree multiple sequence alignments, and variably sized local windows around the genes of interest in the genome (resources.michael.salk.edu/root/home.html).

Non-cannabis genomes included in the scaffolded cannabis Orthofinder run:

1. Amborella trichopoda
2. Aquilegia oxysepala
3. Arabidopsis thaliana
4. Cannabis sativa
5. Carpinus fangiana
6. Carya illinoinensis
7. Ceratophyllum demersum
8. Citrullus lanatus
9. Corylus avellana
10. Cucumis melo
11. Cucumis sativus
12. Fragaria vesca
13. Fragaria X
14. Lotus japonicus
15. Magnolia biondii
16. Malus domestica
17. Manihot esculenta
18. Morus notabilis
19. Nelumbo nucifera
20. Oryza sativa
21. Parasponia andersoni
22. Prunus persica
23. Quercus lobata
24. Rosa chinensis
25. Sechium edule
26. Trema orientale
27. Trochodendron aralioides
28. Vaccinium macrocarpon
29. Vitis vinifera
30. Ziziphus jujuba
31. Humulus lupulus

Non-cannabis genomes included in the full cannabis Orthofinder run:

1. Fragaria vesca
2. Lotus japonicus
3. Malus domestica
4. Prunus persica
5. Rosa chinensis

#### Visualization of synteny with genespace

To visually assess gene-level variation in the haplotype-resolved, chromosome-scale genomes with X and Y chromosomes, we used genespace version 0.9.3 ^58^ within R version 4.2.2 (2022-10-31) ^59^. We initially ran OrthoFinder ^20^ outside of the genespace environment and imported the results. To run the analysis, we used the “synteny” function, followed by “plot_riparianHits.”

#### WGD analysis reveals cannabis has the eudicot hexaploidization

The high quality wild cannabis genome analysis provided evidence for a large whole genome duplication (WGD) or whole genome triplication (WGT) in the past, in addition to more recent WGD events ^60^. We leveraged several chromosome-resolved haplotype genomes representing the different classes of cannabis and sexes to estimate the WGD history as well as whether the sex chromosomes were under different selection across the populations. First, we performed all-all protein alignments within and between cannabis genomes representing type I, type III, feral and fiber. All of the alignments resulted in a syntenic depth of 1:1 with off-diagonal patterns more consistent with unmasked transposable elements (TEs) remaining in the genomes (Supplemental Figure 8 A). We estimated the synonymous substitution (ks) rate between syntenic proteins to determine when genomes diverged, and found that self-self and haplotype-haplotype cannabis alignments revealed early peaks around 1 and 10 MYA respectively (Supplemental Figure 8 B). However, consistent with the syntenic depth analysis, the gene pairs driving the early peak were unmasked TEs. When the unmasked TEs were removed, the Ks distribution suggested genes off the diagonal were duplicated around 100 MYA (Supplemental Figure 8 B). Therefore, as seen in the age analysis of the TEs, and now the pair wise analysis of syntenic TEs, there was a recent burst of TEs in the cannabis genome.

The 1:1 syntenic depth suggests that either cannabis has not had a WGD event recently, or it has been aggressively fractionated back to diploid state with few duplicated regions. Alignment with amborella, which is the sister of the eudicots and lacks a WGD ^61^, resulted in a 3:1 syntenic depth consistent with cannabis having a WGD event after the split with amborella. It has been shown that grape (*Vitis vinifera*) has only the lambda WGT (hexaploidization) that is shared by many eudicots ^62^. Cannabis has a 1:1 syntenic depth with grape consistent with it sharing the WGT with grape but not having another polyploidy event. A high quality genome is also available for the sister species hop (*Humulus lupulus*) ^63^. We found that hop and cannabis share a similar 1:1 syntenic depth as we saw with grape, suggesting that both cannabis and hop share the lambda WGT, but do not have any subsequent WGD events (Supplemental Figure 8 B).

When the TEs were removed from the cannabis self-self syntenic pairs, the off-diagonal (WGD remnant pairs) were consistent with the cannabis genome experiencing a polyploidization event around 100 MYA, which coincides with the lambda WGT shared with grape and hop (Supplemental Figure 8 C). The self-self syntenic pairs for grape and amborella were consistent with them separating around 100 and 150 MYA respectively, further supporting the alignment to the cannabis genome (Supplemental Figure 8 C). Since hop and cannabis share the lambda WGT but do not have any other polyploidization events, the hop self-self syntenic pairs should also be around 100 MYA apart. However, what we observe is similar to cannabis with two peaks around 10 and 30 MYA (Supplemental Figure 8 D), suggesting the hop genome also has a large number of TEs that are not masked. Taken together these results showed that cannabis has one WGT event shared with grape and hop, as well as many other eudicots, but neither cannabis nor hop have had a more recent polyploidization event. However, both the cannabis and hop genomes have been shaped by recent bursts of TEs that may have played a role in separating these species.

The genes that were retained in the cannabis genome after the lambda WGT and fractionation were analyzed for gene ontology (GO) enrichment to identify potential pathways evolutionarily conserved. There were roughly 9,000 genes retained in the cannabis genome and these genes resulted in >700 overrepresented GO terms with the most significant terms relating to disease resistance and response to the environment (Supplemental Figure 8 E, F). While the cannabis genome does not have a recent WGD, it does have a number of tandem duplicated (TD) genes that also play a role in the recent ks peaks. GO overrepresentation analysis of the TD genes revealed that much like the retained genes, disease resistance, response to the environment as well as secondary metabolism were enriched.

#### Structural Variation analysis

The 78 fully scaffolded assembly haplotypes were each aligned to the EH23a assembly using minimap2 (Heng Li 2018). Syri was then used to call structural variations on each alignment (Goel et al. 2019) and plotsr was used to visualize alignments and SVs (Goel and Schneeberger 2022). CDS and TE content were analyzed using bedtools intersect (Quinlan and Hall 2010).

Inversion breakpoint (BP) repeats were called using blastn alignments of inversions with a minimum size of 10 kb. 8 kb windows centered around the start and end breakpoint of each inversion, and were aligned self-to-self, as well as to the BP window pair on the opposing side of the inversion (start-to-end). Only one the top scoring alignment (excluding the full length self-self alignment) was counted per breakpoint. Inverted repeats were called as alignments in opposing orientations and segmental duplications were called for alignments in the same orientation.

#### Phased SNPs

SNPs were also called using Syri on the same assemblies and alignments as described above. SNPs from each of the two haplotypes per sample were merged into single phased genotype calls per sample, and sites with an N as the ALT call were removed (github.com/RCLynch414/SYRI_vcf.sh). Finally, vcftools was used to quality filter and thin SNP sites to a minimum of 1000 bp spacing: --remove-indels --minGQ 20 --remove-indv EH23a--min-alleles 2 --max-alleles 2 --thin 1000 --stdout --recode.

#### LD calculations

Phased SNPs from the scaffolded assemblies were first assessed for r2 correlations in with bin using plink ^64^: --double-id --allow-extra-chr --set-missing-var-ids @:# --maf 0.01 --geno 0.1--mind 0.5 --chr 7 --thin 0.1 -r2 gz --ld-window 100 --ld-window-kb 1000 --ld-window-r2 0--make-bed. Then ld_decay.py was used to make decay curves (GitHub - erikrfunk/genomics_tools), which were plotted with ggplot in R. Separately LD heat maps were made using vcftools: --thin 50000 --recode; and plotted in with LDheatmap in R (sfustatgen.github.io/LDheatmap/).

#### GO terms

Gene ontology term enrichment tests were performed with the TopGO package in R, using the all high confidence gene annotations of EH23a as the null distribution and classic Fisher test of significance ^65^.

#### TreeMix

The TreeMix model was run using only SNPs outside of gene models: -seed 69696969 -o out_stem -m 5 -k 50 -noss -root asian_hemp. One to 10 migration scenarios were simulated, and ranked based on the ln(likelihoods). Five migration events (-m = 5) was selected as the most likely final number.

#### Local PCA

The local PCA method was applied to the phased SNPs, with 1000 bp minimum spacing between SNPs, and genome windows of 100 SNPs ^66^

#### Disease resistance gene analog (RGA) analysis

Plant disease resistance gene analogs (RGAs) are defined by the presence of one or more highly-conserved amino acid motifs in their encoded proteins. These motifs encode functional protein domains that determine pathogen specificity and subcellular localization. Depending on the particular pathosystem, RGA proteins can be entirely cytoplasmic, or can span the cell membrane with cytoplasmic functional domains, extracellular domains, or both.

Drago2 ^67^ was used to identify motifs conserved among plant disease resistance gene analogs. Input files were transcript annotation fasta files for each genome. Sets of genes containing both NBS and LRR domains were used as input to MEME to assess and compare amino acid composition in motifs over gene sets.

To identify the specific CNL related to powdery mildew resistance, the proximal gene to a marker mapped to chromosome 2 in CBDRx (Mihalyov and Garfinkel, 2021) was used as a blastn query against the 38 chromosome-scaffolded genomes. The resulting top blast hits did not overlap with any gene annotations; however, 16 of the 38 genomes had blast hits on chromosome 2 with >95% nucleotide identity to the CBDRx gene; of these, nine of these had 99-100% nucleotide identity over all three exons (1745 bp, 1448 bp, and 287 bp), respectively.

Sequences from five of the 16 genomes (H3S7a, OFBa, SZFBa, TKFBa,WCFBa) clustered separately from the rest. These were distinguished by a 1-bp insertion in the first exon, ten small indels (2-8 bp) in exonic space, and a 1280 bp longer second intron. These regions were extracted and aligned with the CBDRx gene sequence, and the alignment was used to produce a maximum-likelihood tree (Supplemental Figure 19 D).

CNLs showed a distinct pattern on chromosomes 3 and 6. There were one to two CNL genes between 400-600 kb; two to four between 1-1.4 Mb; one to two at 6-8 Mb; a single CNL gene near the near the centromeric region of the chromosome at 35-37 Mb, and one to five (COFBa) CNLs between 78-84 Mb. Exceptions to this pattern were OFBa, H3S1a, and MMv31a, which lacked a CNL in the centromeric region. In SDFBa and SN1v3a, the centromeric CNLs were located at 42.8 and 47.5 Mb, respectively. SN1v3a had a CNL at 12.2 Mb, another exception to the overall pattern. Chromosome 3 in this genome was larger than the others, at 90 Mb, compared to the rest at 80-85 Mb. Finally, GERv1a lacked a CNL in the 78-84 Mb region of chromosome 3.

#### Identification of terpene synthase genes

Each of the *Cannabis* proteomes was aligned to a set of 40,926 protein sequences from UniProt (search criteria “Embryophyta” and “reviewed”; accessed on 09/20/2022) with blastp (version blast 2.6.0, build December 7, 2016) ^68^. Alignment thresholds included an E-value threshold of less than 1e-3, at least 20% query coverage, and a percent identity based on the length of the alignment ^69^. Terpene synthases were also identified based on the presence of Pfam domains, PF01397 and/or PF03936 ^70^. To assess domain content, each of the *Cannabis* proteomes was aligned to the Pfam-A.hmm database (last modified 11/15/2021; accessed 09/20/2022) ^71^ with hmmscan (HMMER 3.3.2 November 2020) ^72^ on default settings.

#### Identification of genes in the precursor pathways for terpene and cannabinoid biosynthesis

Terpene biosynthesis proceeds via two pathways: the chloroplastic methyl-D-erythritol phosphate (MEP) pathway, which produces precursors for monoterpene and cannabinoid biosynthesis, and the cytosolic mevalonate pathway, which produces precursors for sesquiterpene biosynthesis. The protein sequences for these pathways ^73–75^ were aligned to each of the *Cannabis* proteomes with diamond version 2.1.4 on default settings ^76^.

#### Synthase cassette analysis

To identify full and partial length cannabinoid synthases in each of the cannabis genomes, the reference cannabinoid synthase sequences were aligned to the genome with blastn. An enriched LTR sequence developed from CBDRx ^36^ was used as a reference to further aid in the identification of synthases. LTR08 is an LTR sequence from the CBDRx genome. Blast hits were then piped into a python script for filtering and visualization.

A Python script was written to take in cannabinoid synthase blast results and LTR08 blast results in table format. Synthase hits with length <500 bp were filtered out. LTR08 hits with bitscore <1250 were filtered out. Synthase and LTR08 hits with mismatches <10 and zero gaps were labeled as “Full” sequences. All other hits were labeled as “Partial” sequences. Hits that shared the same starting position were then filtered to a single sequence and given one of the synthase labels according to the following. Full hits were retained and labeled as the corresponding functional synthase. Partial hits within 60 kb of an LTR08 hit upstream or downstream were labeled as CBDAS and retained. If there were no Full hits or hits with an LTR08 in proximity, the hit with the highest bitscore was labeled as the respective synthase and retained. The filtered and labeled synthases were then plotted onto a track to visualize cannabinoid synthase orientation for each region of a genome. A minimum of 4 synthase hits was required for visualization.

Inkscape was used to visualize synthase cassette tracks. Manual edits were used to correct a few incorrect labels between CBDAS and CBCAS. Synthase cassettes are grouped by overall cassette shape.

#### Cannabinoid synthase gene analysis

First ORFinder was used to remove pseudogenes from the initial list of potential genes described above (ftp.ncbi.nlm.nih.gov/genomes/TOOLS/ORFfinder/linux-i64/). Then we used usearch11.0.667 to cluster synthase coding sequences: -cluster_fast -id 0.997 -sort length-strand both -centroids -clusters. ^77^ TranslatorX was then used to produce protein guided multiple sequence alignments ^78^. Synthase evolutionary history was inferred by using the Maximum Likelihood method and General Time Reversible model in MEGA11^79^.

#### Kmer Crossover analysis

We used the anchoring function of PanKmer to locate crossover events in known trios of Cannabis genotypes. Eleven trios included FB191 as a varin donor parent and 6 trios included SSV as a varin donor parent. The parents of FB191 are HO40 and FB30, while the parents of SSV are HO40 and SSLR; in both cases, HO40 was the varin donor. For each trio, the F1 genome was haplotype-resolved and included one haplotype from a varin-donor parent and one from a non-varin donor parent. In each case, we used PanKmer anchoring to identify the “varin haplotype”. For FB191 trios, we generated a 31-mer index of the FB191 genome using “pankmer index” with default parameters. Using a Python script importing PanKmer’s API functions “pankmer.anchor_region()” and “pankmer.anchor_genome()” ^80^, we anchored the FB191 index in each haplotype of the cross, for example COFBa and COFBb. We identified the varin haplotype as the haplotype with higher 31-mer conservation in the FB191 index. We applied the same procedure to SSV trios using a PanKmer index of SSV. We then sought to trace potential varin alleles from HO40 to the varin haplotype of the cross. To represent HO40, we generated two single-genome 31-mer indexes: one for the HO40 genome and a second for the highly similar EH23a sequence. We also generated single-genome 31-mer indexes of FB30 and SSLR. For each FB191 cross, we anchored the HO40 index, EH23a index, and FB30 index in the varin haplotype. We inferred crossover events at loci with a clear ‘haplotype switch’ indicated by k-mer conservation values. We repeated the same procedure for SSV trios, applying the SSLR index in place of the FB30 index. All scripts for this analysis are available on GitLab.

#### Varin SNP association tests and genetics

First, the BestNormalize package in R was used to select the arcsinh method to transform the varin ratio data, which were initially deemed multimodal. Then the model BLINK from the GAPIT package in R^81^ was used with PCA.total=6 to test associations between SNPs in the F2 population and transformed varin ratio data (Supplemental Table 11). This PCA.total parameter was selected based on visual evaluation of QQ plots for PCA.total values 1-10, where 6 was the smallest number that did not show systemic inflation of p-values ^81^. Next, gene and TE models were manually assessed for windows within 50 kb of the four F.D.R. corrected significant SNPs. Then, Orthofinder groups for *BKR*, *ALT3*, and *ALT4* were extracted, and the three *ALT* 3 and 4 orthogroups were pooled into a single set of *ALT* gene counts. Phylogenies of *BKR* and *ALT* protein sequences were constructed in MEGA with the neighbor joining method from the orthogroups using 100 bootstrap replicates ^79^. The BKR alignment and translation displayed in Figure 6 was made using the Geneious ^82^ alignment algorithm on default settings.

## Supplemental Figures and Table

**Supplemental Figure 1.**
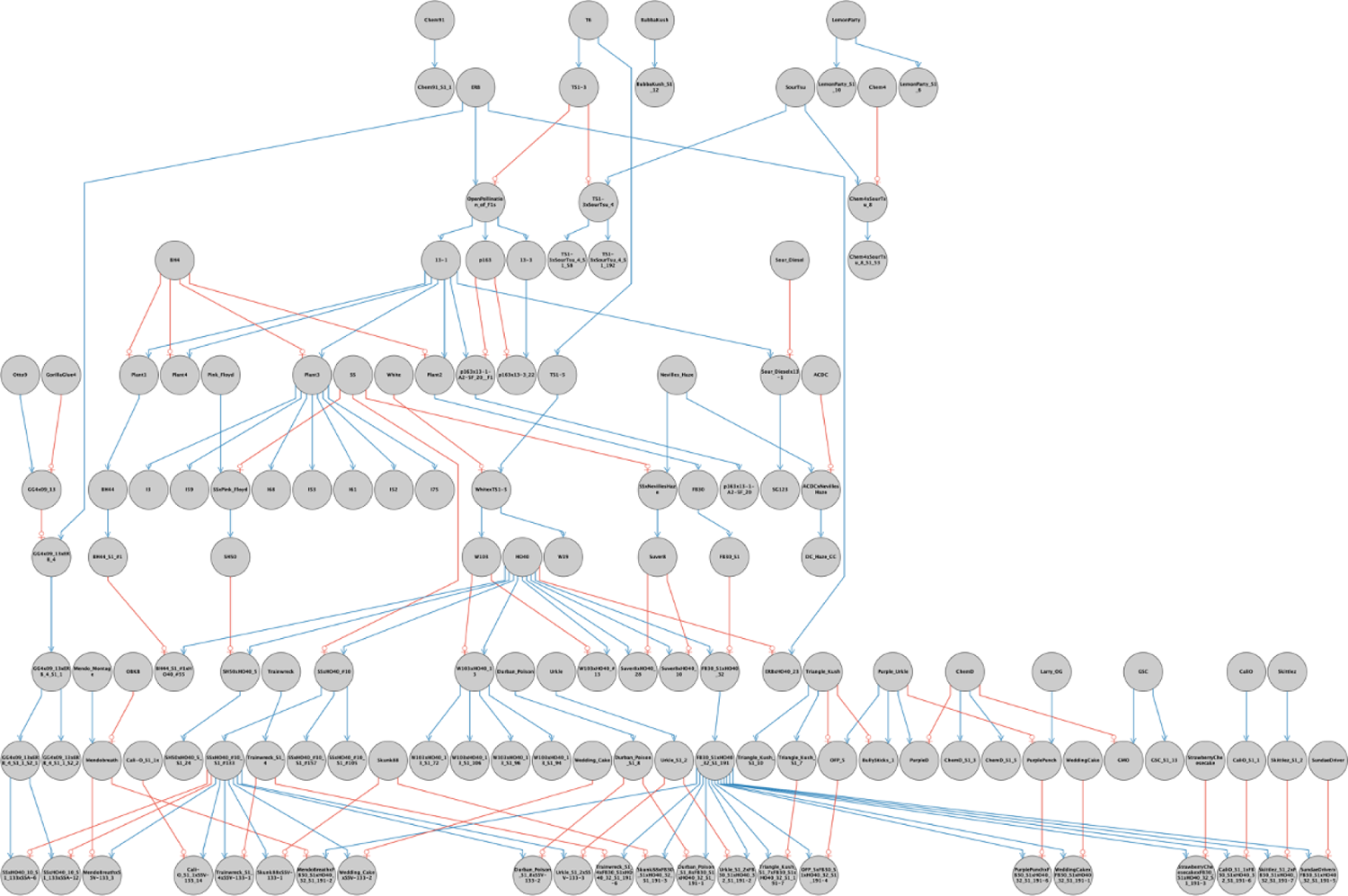
The Pedigree of the OCBD cultivars sequenced for the cannabis pangenome. A high proportion of the pangenome is based on the Oregon CBD (OCBD) breeding program including a specific strategy to produce high varin lines. Pedigree of the OCBD breeding program to generate high varin CBGA plants. HO40, which is a high varin plant, was crossed with the high CBGA plant FB30_S1 (100:1 CBGA:THCA, DN dominant) and selfed. Then, a high varin plant was selected (FB191) for crossing to type I plants for high CBGV production.

**Supplemental Figure 2.**
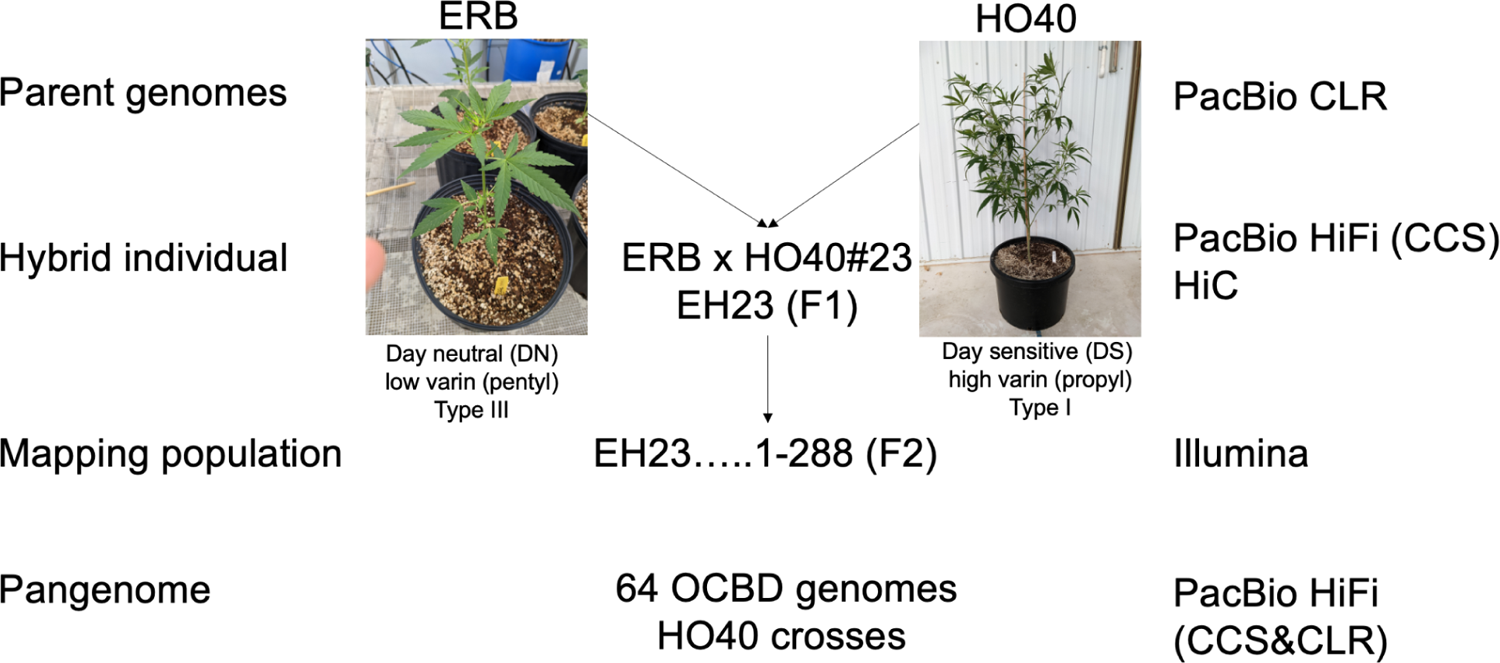
The EH23 anchor genome sequencing strategy and resulting populations. The F1 hybrid EH23 (ERBxHO40#23) was generated by crossing the type III (high CBDA), day neutral (DN), Early Resin Berry (ERB) with the type I (high THC), day sensitive (DS), HO40. Both ERB and HO40 were sequenced with PacBio CLR, while EH23 was sequenced with PacBio HiFi (CCS) and scaffolded with HiC. The F2 mapping population (288 individuals) was sequenced with Illumina short reads. The remaining pangenome samples from OCDB are summarized in Supplemental Table 1 and Supplemental Figure 1 pedigree.

**Supplemental Figure 3.**
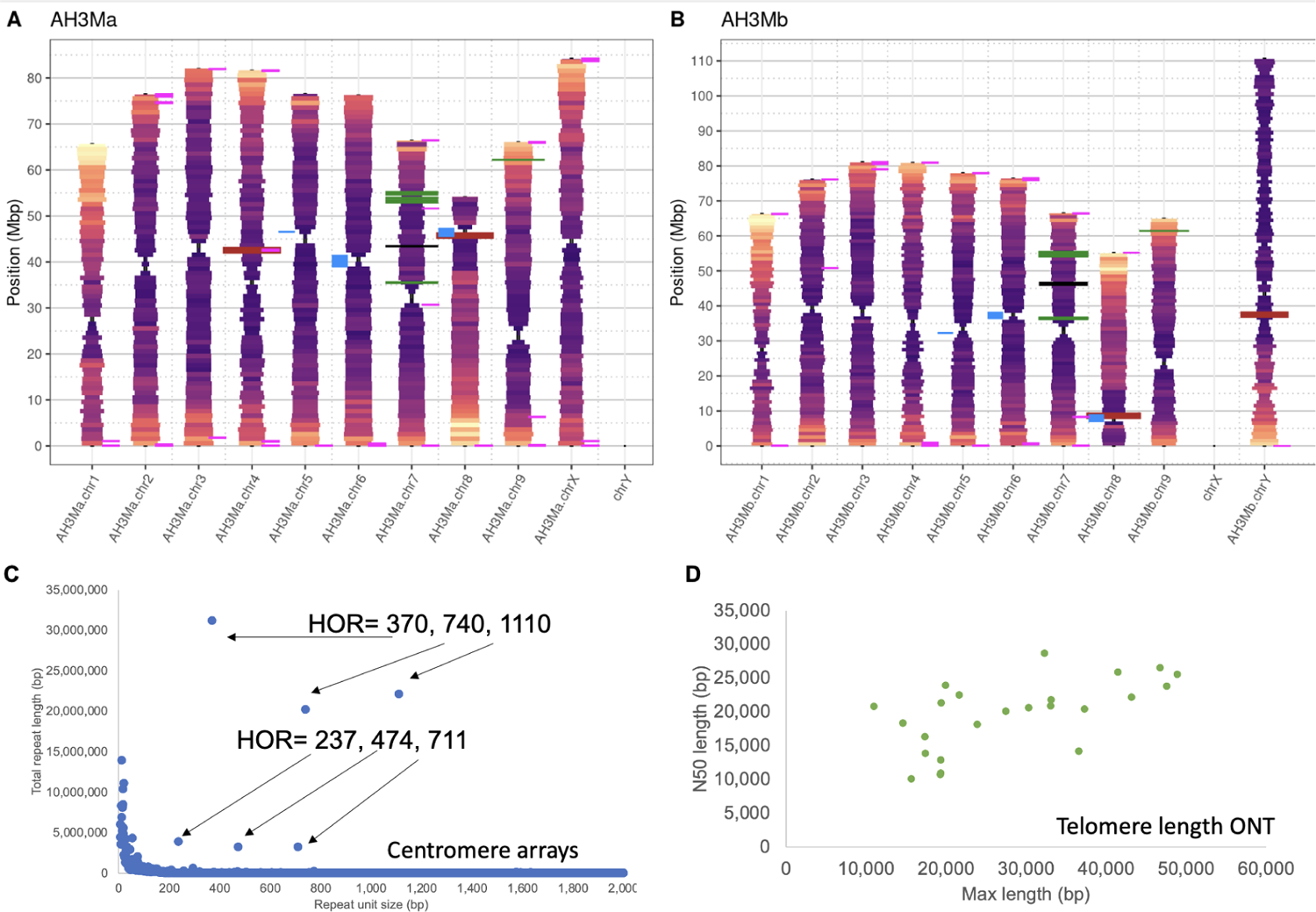
Cannabis centromere and telomere analysis. A-B) The AceHigh3 (AH3M) chromosomal features of nine pairs of autosomes and one pair of sex chromosomes (X and Y). One million base pair rectangular windows extend outward from each pair of haplotypes at a width proportional to the absence of the CpG motif. Each rectangular window is colored by gene density with warm colors indicating high gene density and cool colors indicating low gene density. Each pair of haplotypes is connected by polygons indicating structural arrangement, with gray for syntenic regions and orange connecting inversions. Rectangles along each haplotype indicate select loci, including 45S (26S, 5.8S, 18S) rDNA arrays (firebrick red), 5S RNA arrays (black), 237 bp centromere repeat (blue), 370 bp CS-1 sub-telomeric repeat (pink) and cannabinoid synthases (forest green; CBCAS, CBDAS, THCAS, and OAC). Chromosomal plots for all 78 haplotype-resolved, chromosome-scale genomes show similar trends (Additional data). C) The centromere arrays identified in the AH3M genome (as an exemplar for the pangenome) with Tandem Repeat Finder (TRF). Two high copy number arrays were identified with base repeats of 237 and 370 bp, along with their higher order repeats (HOR). The 237 bp array is sparsely found in the genome (blue, panel A), although usually proximal to the high “CpG” sites putatively demarcating the centromere regions. The 370 bp repeat is the same sequence as the sub-telomeric repeat CS-1 ^37^ and found on the ends of the chromosomes (pink, panel A). D) A subset of the genomes were sequenced on Oxford Nanopore Technologies to estimate the telomere length in cannabis genomes ^33^. The N50 ONT read length is plotted as a function of the max telomere repeat identified using the TeloNum software.

**Supplemental Figure 4.**
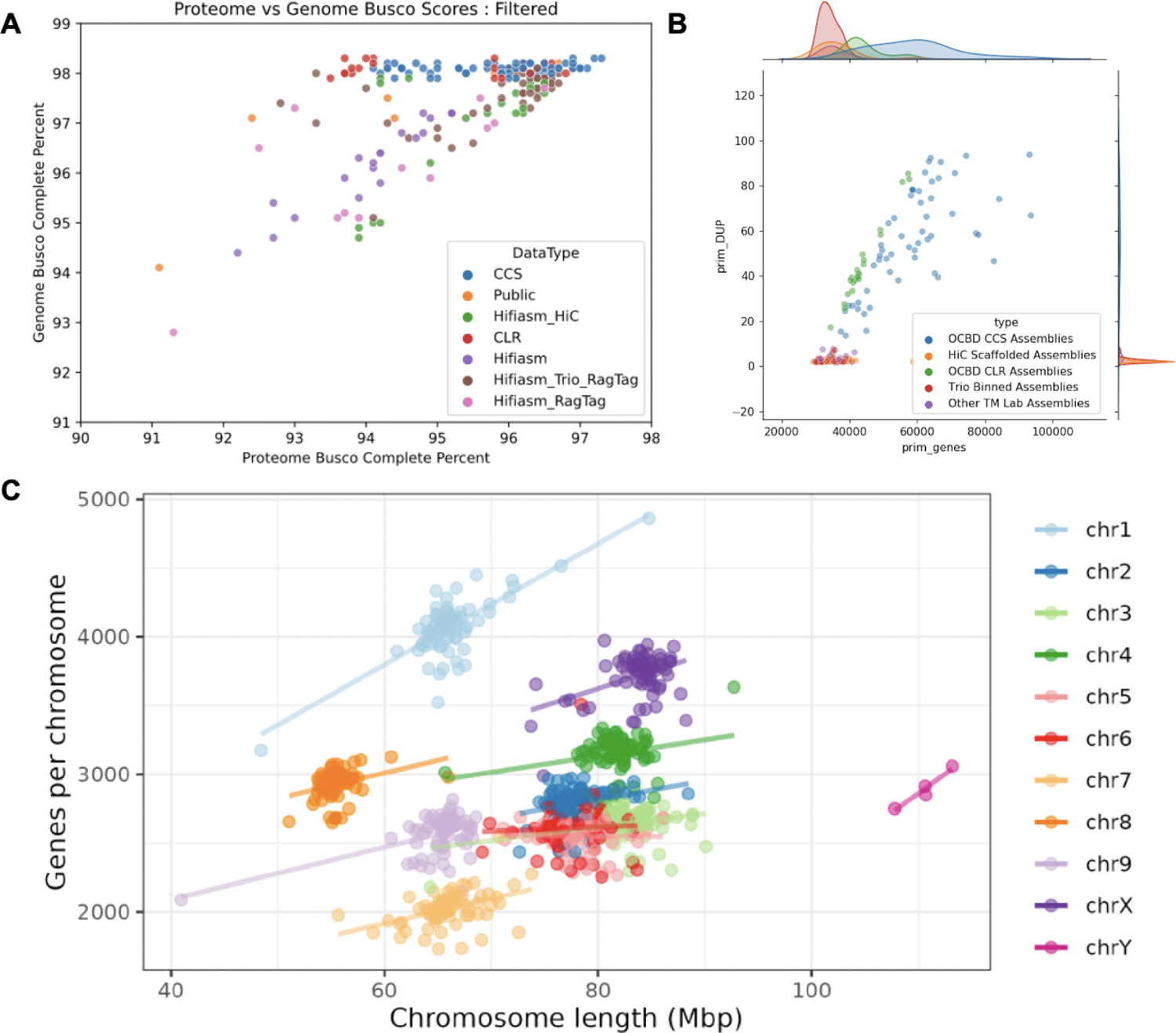
The cannabis pangenome and pan-genes are high quality. A) Benchmarking Universal Single-Copy Orthologs (BUSCO) ^83^ for both the genome and gene predictions suggest that they are both high quality and complete. B) The number of genes predicted (“prim_genes;” primary genes; no splice variants) contrasted with the number of BUSCO duplicate (prim_dup) genes suggests that the CCS and CLR contig based assemblies are retaining significant duplicated sequence due to uncollapsed haplotypes. These haplotypes were not removed to retain the level of variation for downstream analysis. C) Scatter plot of chromosome lengths on the x-axis compared with gene counts per chromosome on the y-axis across the 9 autosomes and both sex chromosomes.

**Supplemental Figure 5.**
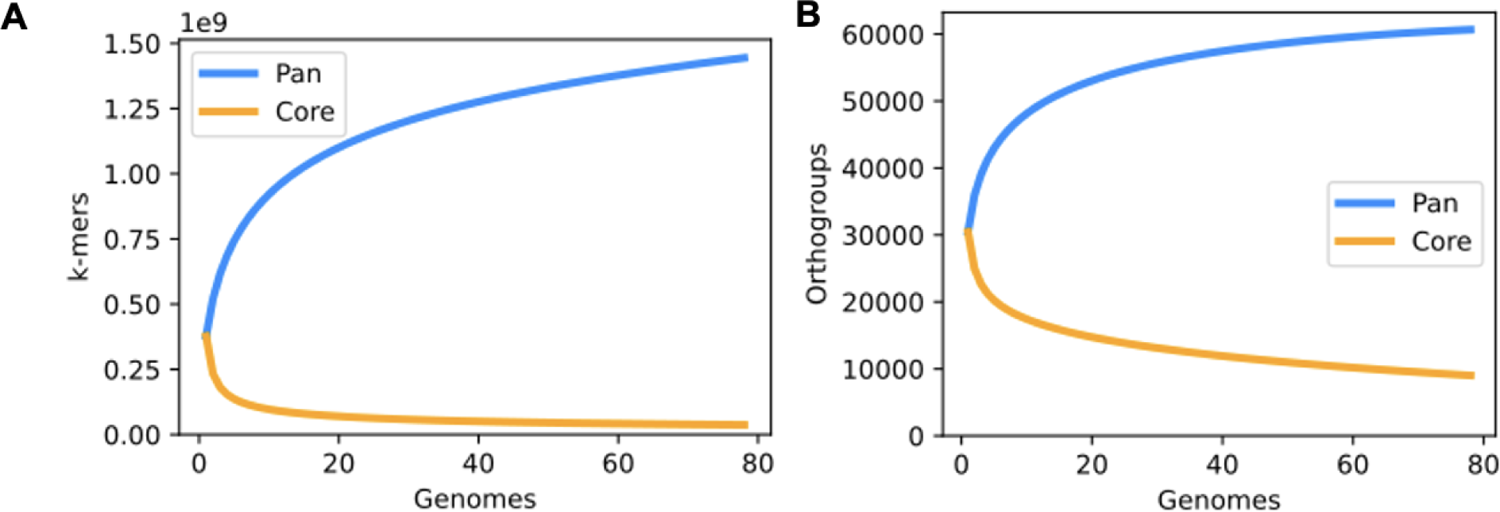
Collector’s curves (pangenome curves) for the 78 haplotype-resolved, chromosome-scale assemblies. A) The collector’s curve is based on PanKmer (31 bp K-mer). B) The collectors curve is based on orthofinder gene families.

**Supplemental Figure 6.**
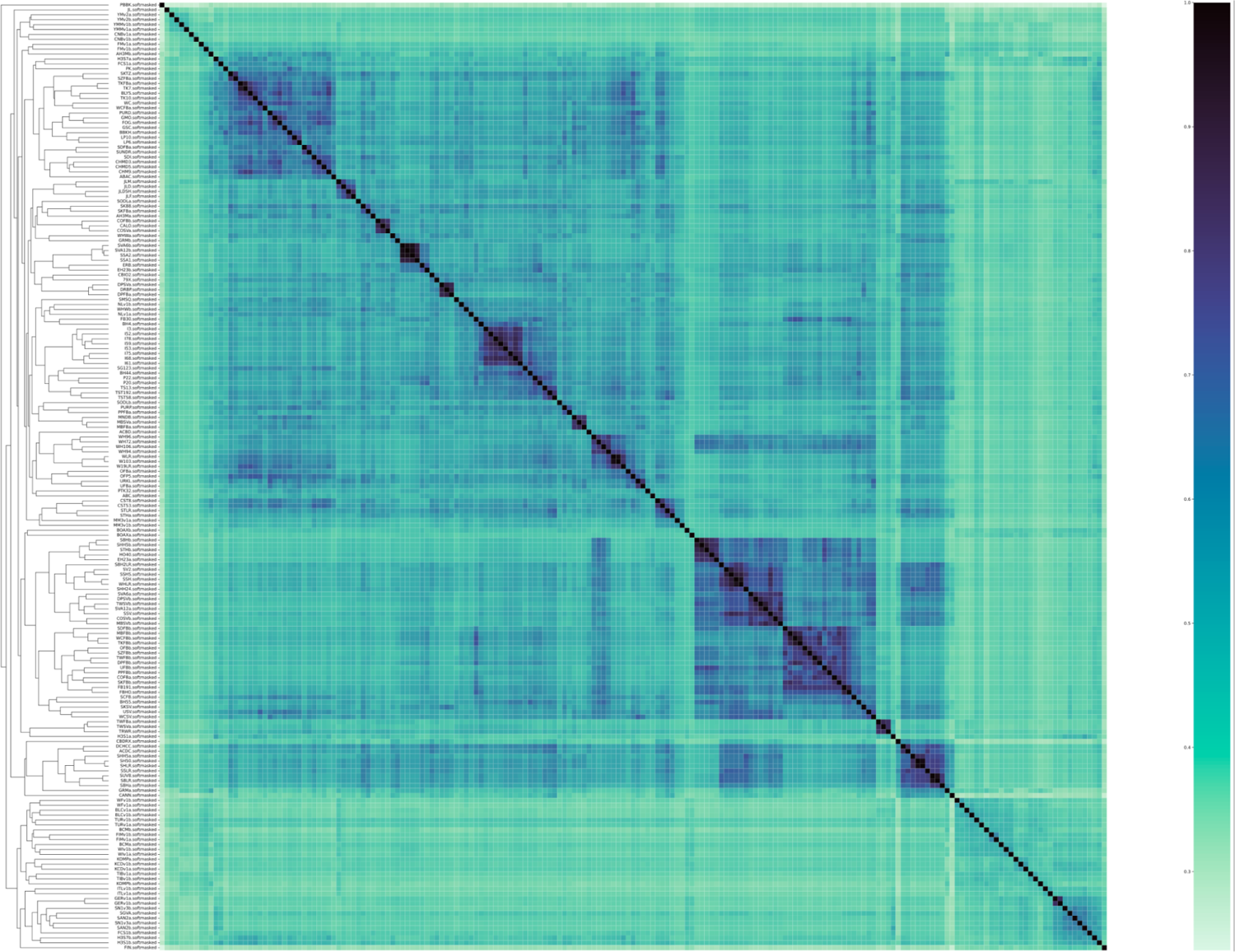
PanKmer Jaccard similarity matrix of 193 Cannabis genomes. PanKmer was used to estimate the relationship between the genomes in the cannabis pangenome. See <u>Figshare for full resolution version.</u>

**Supplemental Figure 7.**
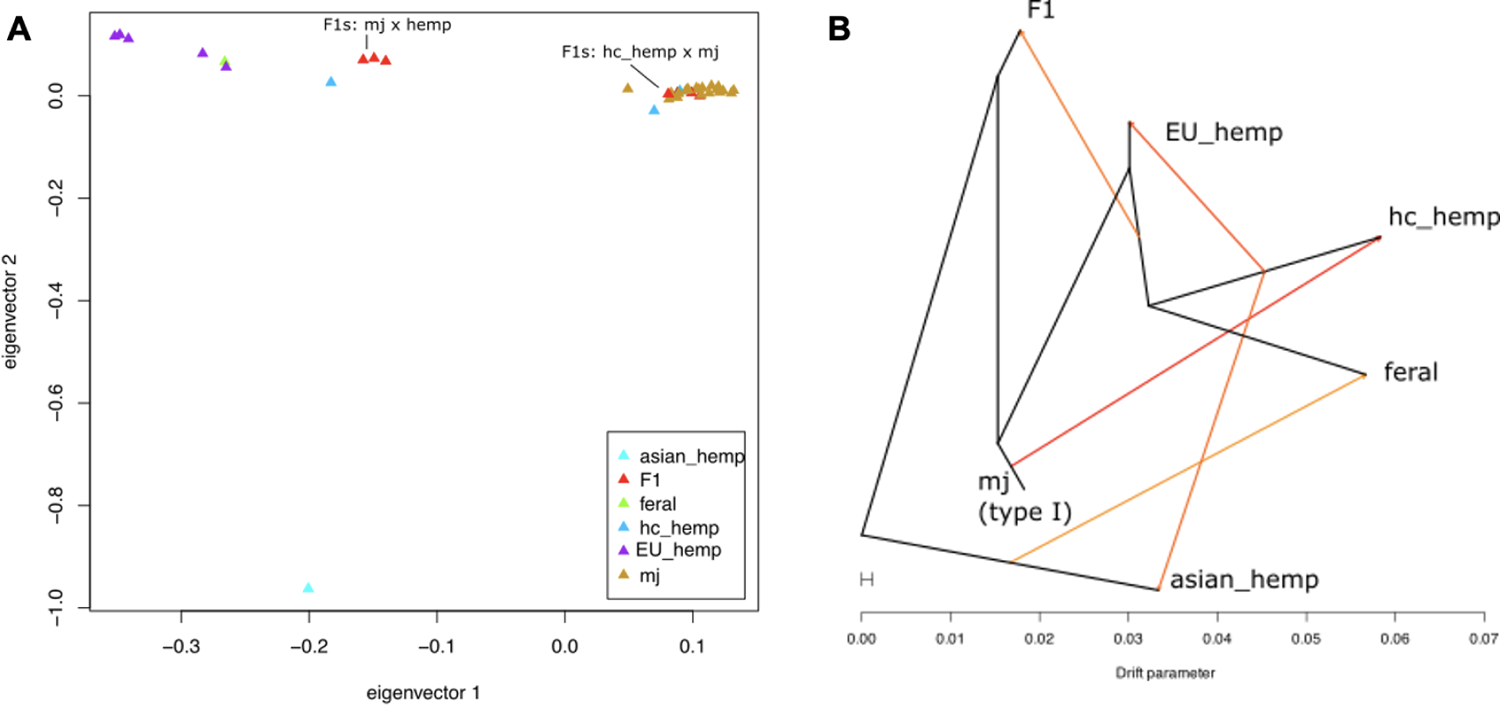
Cannabis pangenome analysis of the 78 haplotype-resolved, chromosome-scale assemblies for breeding history. A) Principal component analysis (PCA) of 494,603 high confidence phased SNPs from 78 haplotype-resolved, chromosome-scale assemblies (39 samples). Population assignments are based on pedigree and provenance, as well as PCA. This analysis appears to reflect differences between European hemp, Asian hemp and drug-type samples, but shows little differentiation among drug-type samples; therefore, we assigned all drug-type samples as either high cannabinoid (hc) hemp or marijuana (MJ) for downstream SNP-based analyses. Samples that are known breeding program hybrids between these major groups were classified as F1s. B) TreeMix SNP based phylogeny shows likely relationships between major populations including 5 hybridization events, based on SNPs from 78 haplotype-resolved, chromosome-scale assemblies.

**Supplemental Figure 8.**
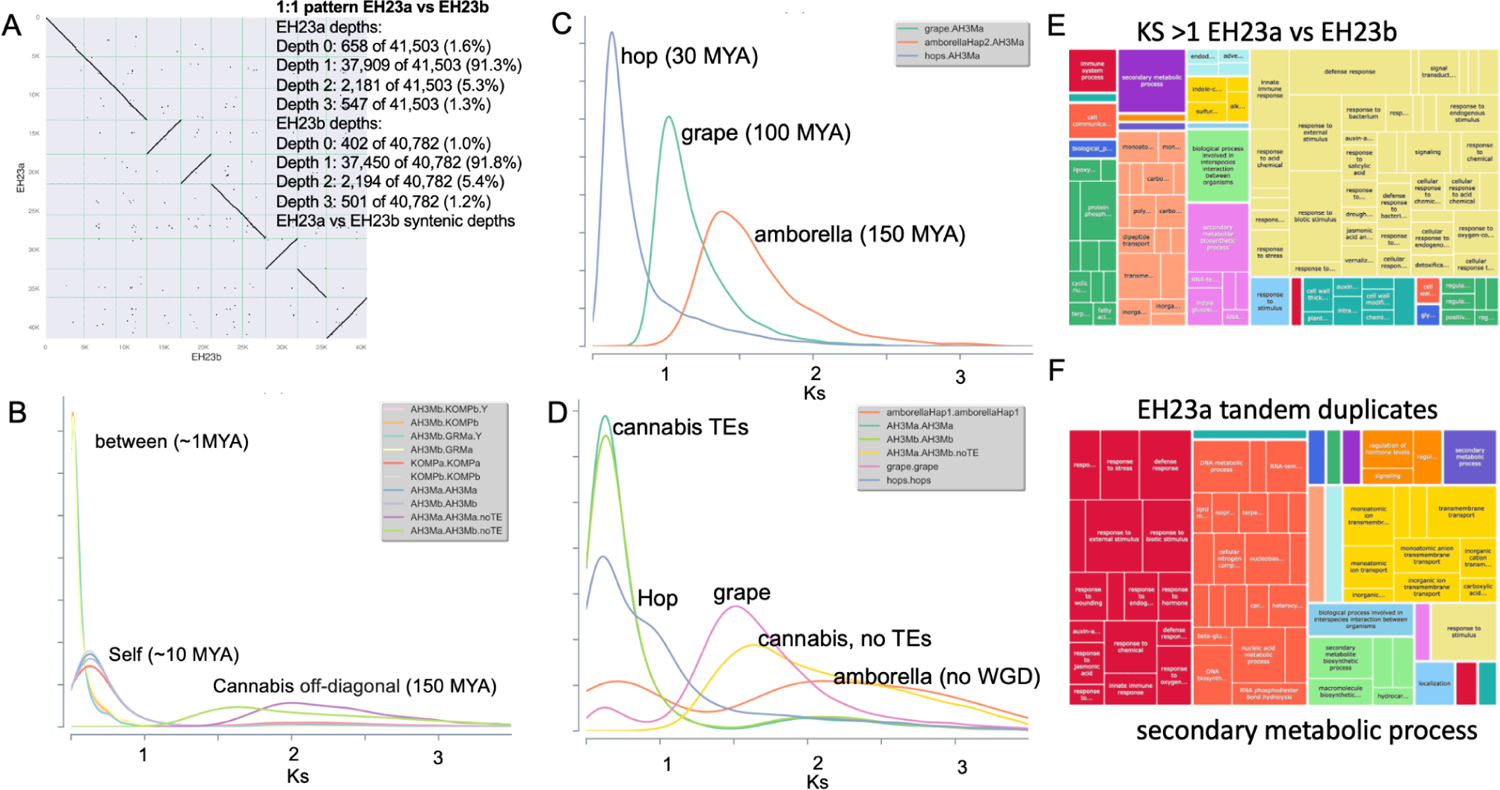
Whole genome duplication (WGD) history of cannabis is consistent with the lambda WGT and a recent burst of TEs and TDs. A) Protein-based dotplot comparing EH23a (HO40) with EH23b (ERB). The inset provides the syntenic pattern numbers showing that both genomes have few unique genes (∼1%) not found in the other genome. 91% of genes are found in 1:1 syntenic relationship between the two haplotypes consistent with very few genes retained after whole genome duplication (WGD). B) The Ka/Ks (non-synonymous/synonymous substitution rate ratio) was estimated across several genomes in the pangenome and Ks was used to estimate the divergence time between genomes. The Ks between cannabis haplotypes shows a peak around 1 million years ago (MYA). In contrast, self-self Ks suggests a peak around 10 MYA. When TE-associated genes are removed, the off-diagonal Ks peak moves in line with the lambda whole genome triplication (WGT) found in many eudicots. C) AH3Ma haplotype was compared to amborella, which is sister to the eudicots and is typically used as a baseline for no WGD, and grape, which only has the lambda WGT. These Ks plots show the divergence between the two species is consistent with the Ks peaks. Cannabis diverged from hop ∼30 MYA, from grape ∼100 MYA around the lambda WGD, and from amborella 150 MYA. D) The timing of the last WGD was refined by looking at the self-self Ks values per species. E) The genes with a Ks > 1 are thought to be under positive selection. Gene ontology (GO) enrichment was performed for the genes with a Ks > 1 and the resulting GO categories were plotted using the tree-map function in Revigo. F) The cannabis genome has a large number of tandemly duplicated (TD) genes. Gene ontology (GO) enrichment was performed for the TD genes and the resulting GO categories were plotted using the tree-map function in Revigo.

**Supplemental Figure 9.**
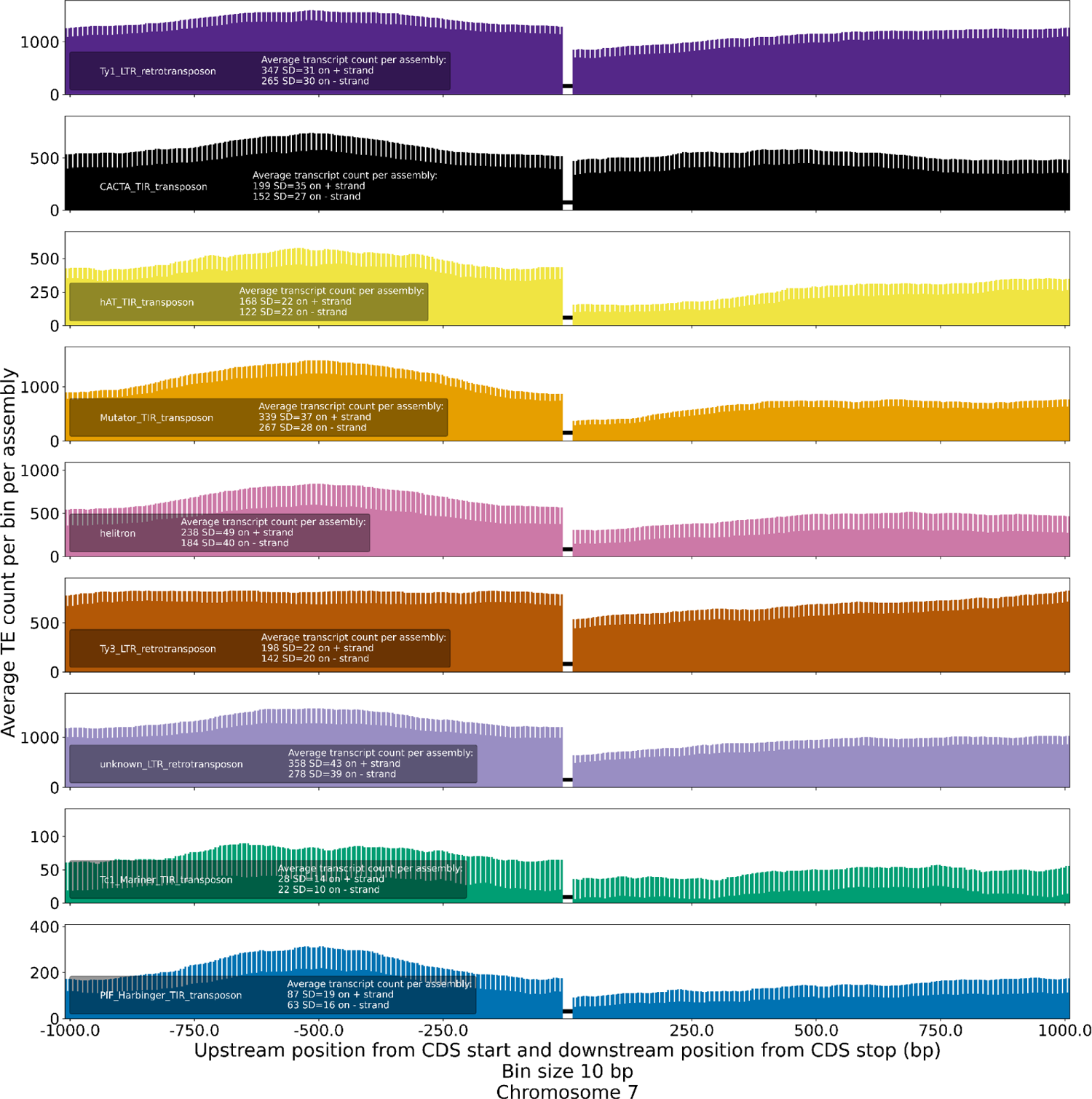
Position-specific density of transposable elements (TEs) upstream (1 kb) and downstream (1 kb) of genes in the cannabis pangenome (78 scaffolded genomes). Each panel corresponds to a different TE, with the histograms showing the average TE count across all scaffolded assemblies in the region upstream and downstream of genes, along with error bars. The horizontal line in the center of each of the panels denotes the gene body.

**Supplemental Figure 10.**
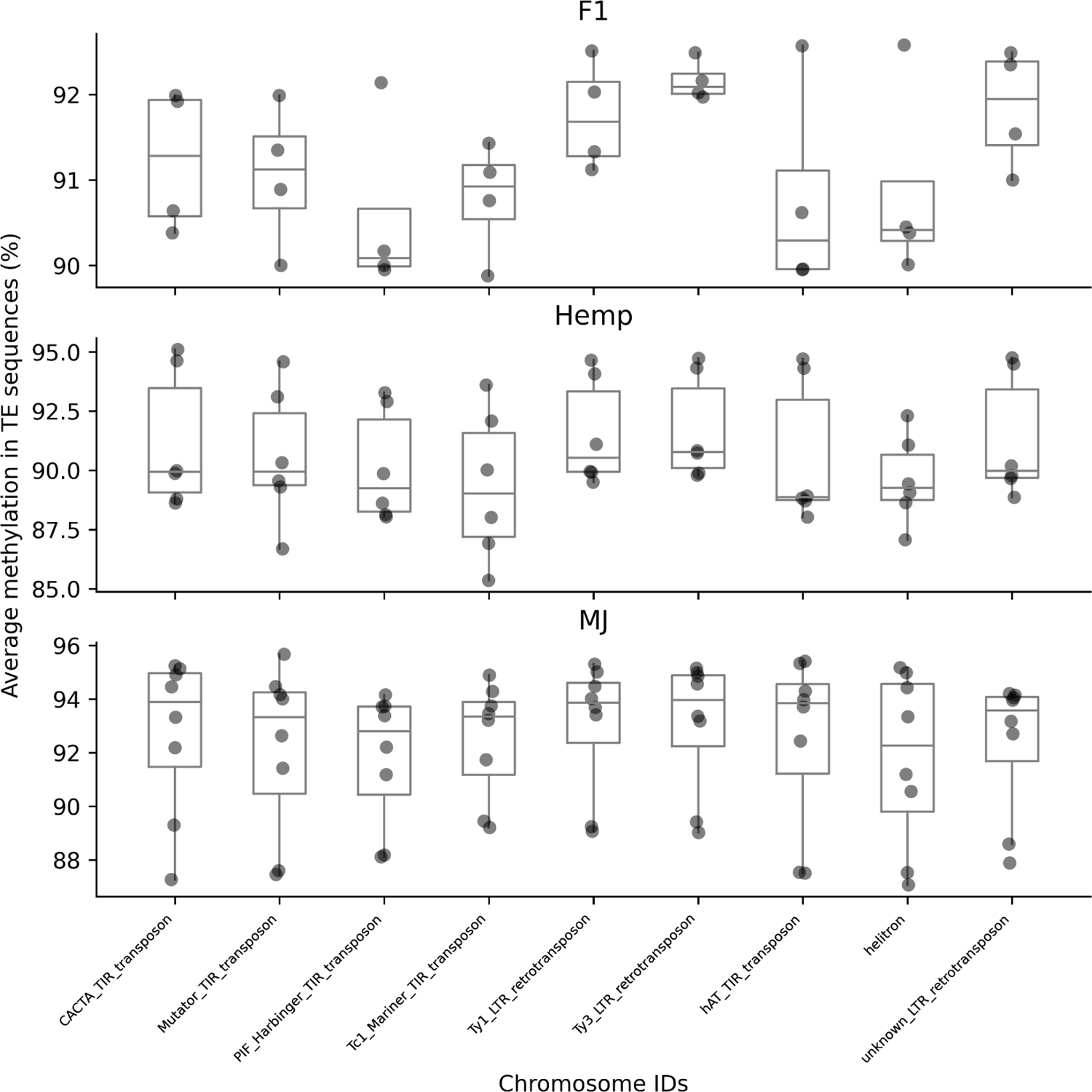
Average methylation (%) in TE sequences. Analysis applies to the subset of genomes with methylation data (see Supplemental Table 6). Each data point in the box plot corresponds to a specific assembly. For each genome, the average methylation value (%) along the length of a TE sequence was averaged for all TE sequences. MJ samples have a slightly higher average methylation value in the TEs than F1 samples.

**Supplemental Figure 11.**
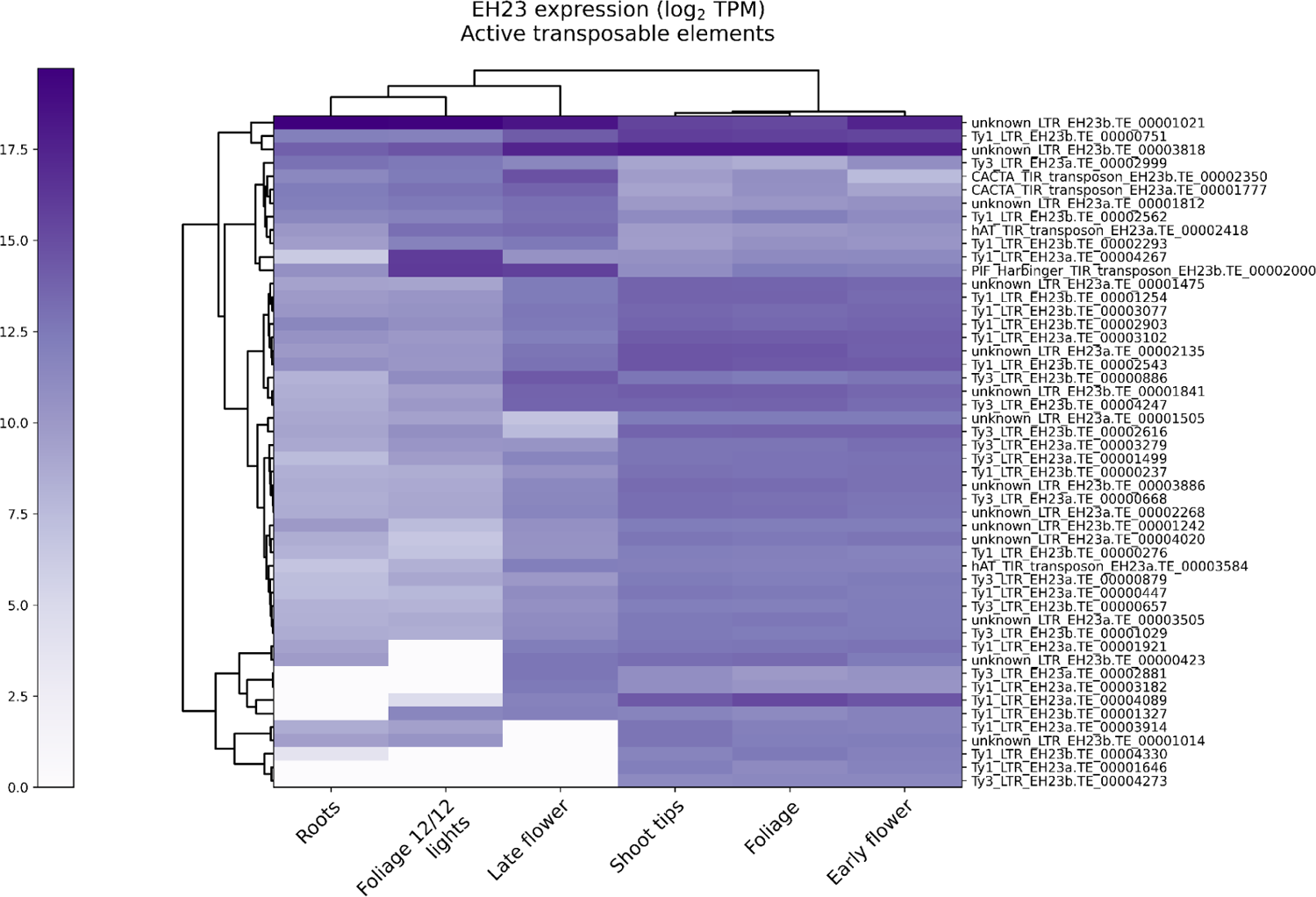
Heatmap showing top 50 expressed TEs in EH23. The majority of active TEs are LTRs, and putative active DNA TEs in EH23 include CACTA, hAT, and Harbinger. Active TEs show tissue-specific patterns, with late-stage flowers showing higher Harbinger activity and lacking activity that is present in shoot tips, foliage, and early-stage flowers.

**Supplemental Figure 12.**
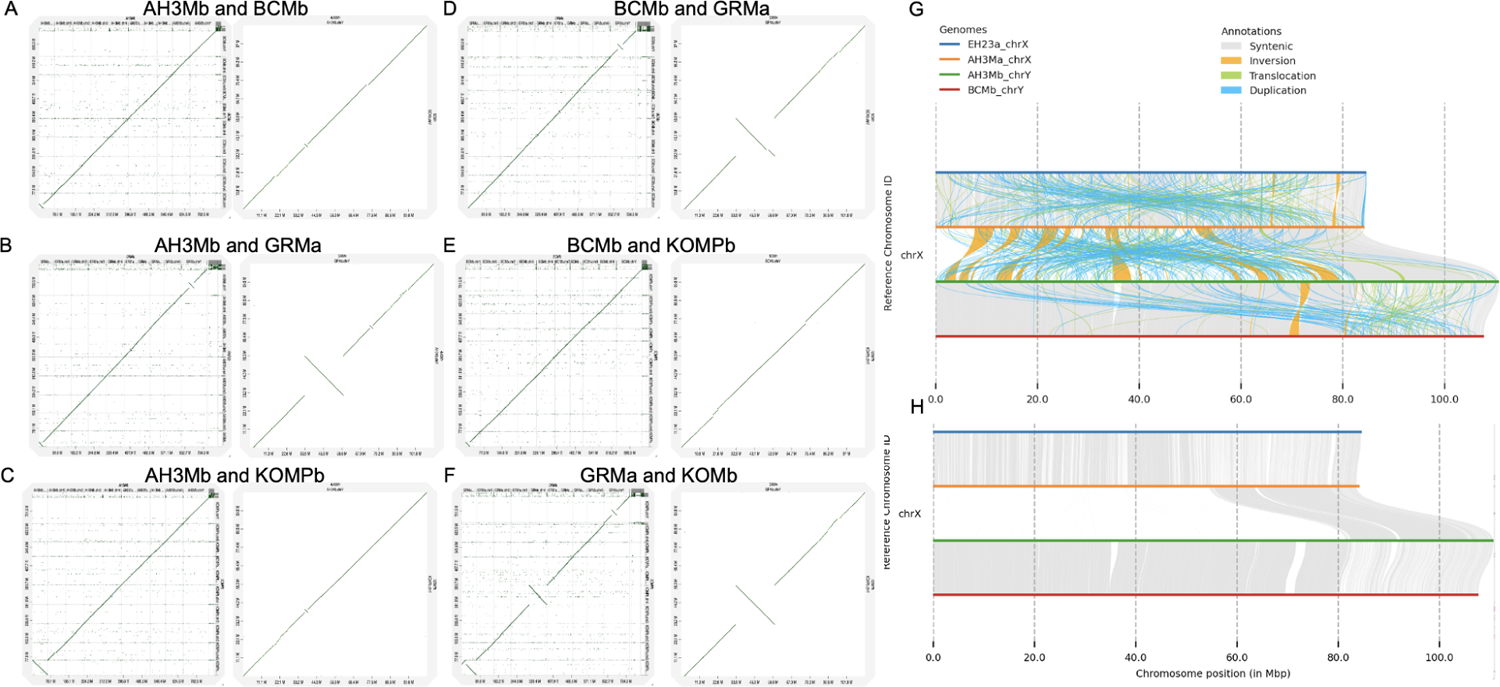
The Y chromosome variation across the cannabis pangenome. A-F) Dotplots between male cannabis genomes that include a Y chromosome. The plot on the left is the whole genome DNA-based dotplot, and the right is just the Y chromosome. G-H) Syntenic plots of the X and Y chromosomes from EH23a (X), AH3Ma (X), AH3Mb (Y), and BCMb (Y) were plotted with syntenic regions (grey), inversions (orange) translocations (green), and duplications (blue), and with just syntenic regions using syri ^28^.

**Supplemental Figure 13.**
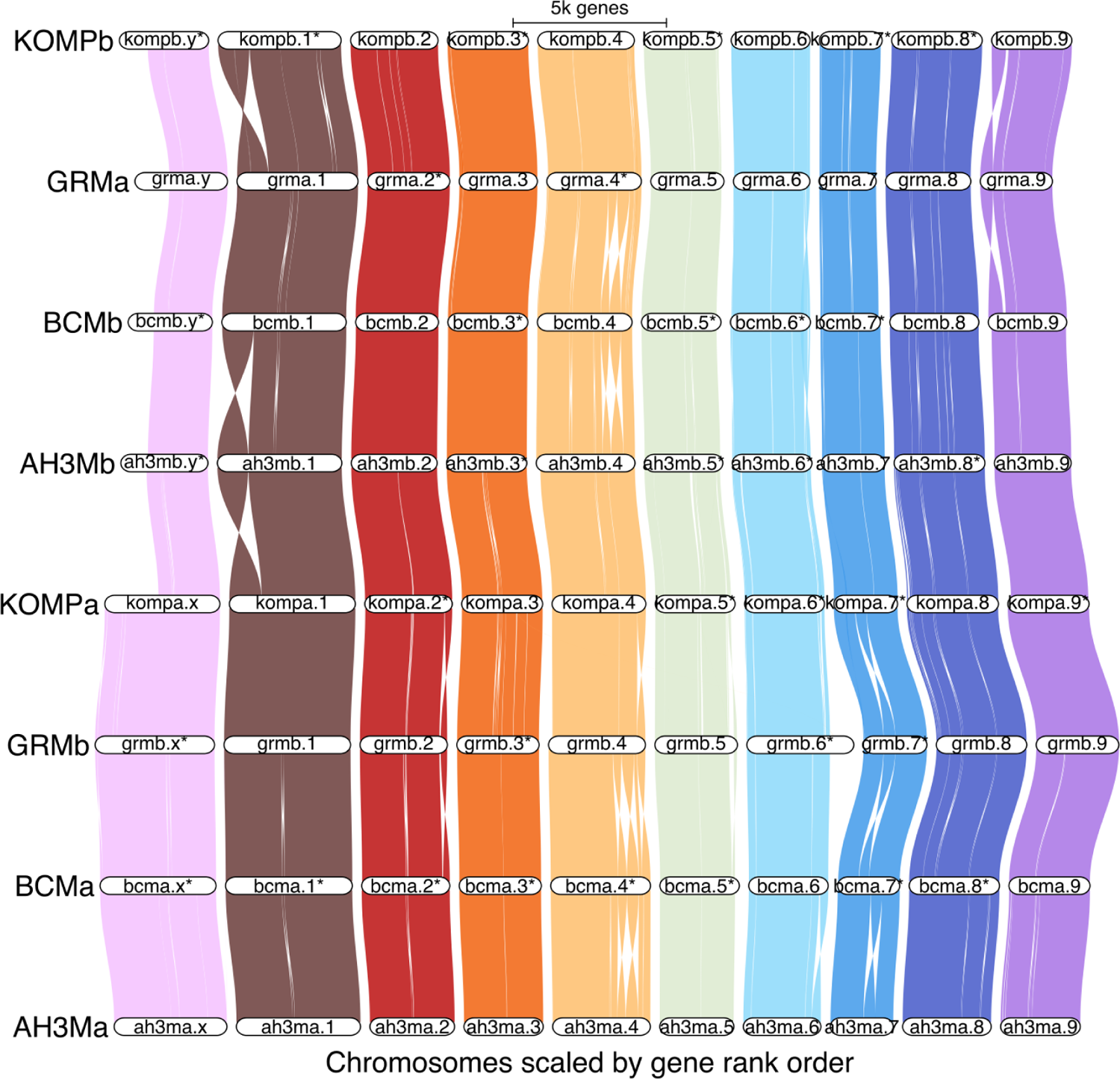
Genespace visualization of haplotype-resolved, chromosome-scale assemblies with X and Y chromosomes, as well as nine autosomes. Chromosomes are not drawn to scale based on physical positions; positions on the x-axis are scaled by gene rank order. The PAR boundary is drawn between KOMPa chromosome X and AH3Mb chromosome Y. Chromosome 1 has a large inversion, and there are smaller inversions in different chromosomes. However, gene-based synteny is broadly consistent across genomes.

**Supplemental Figure 14.**
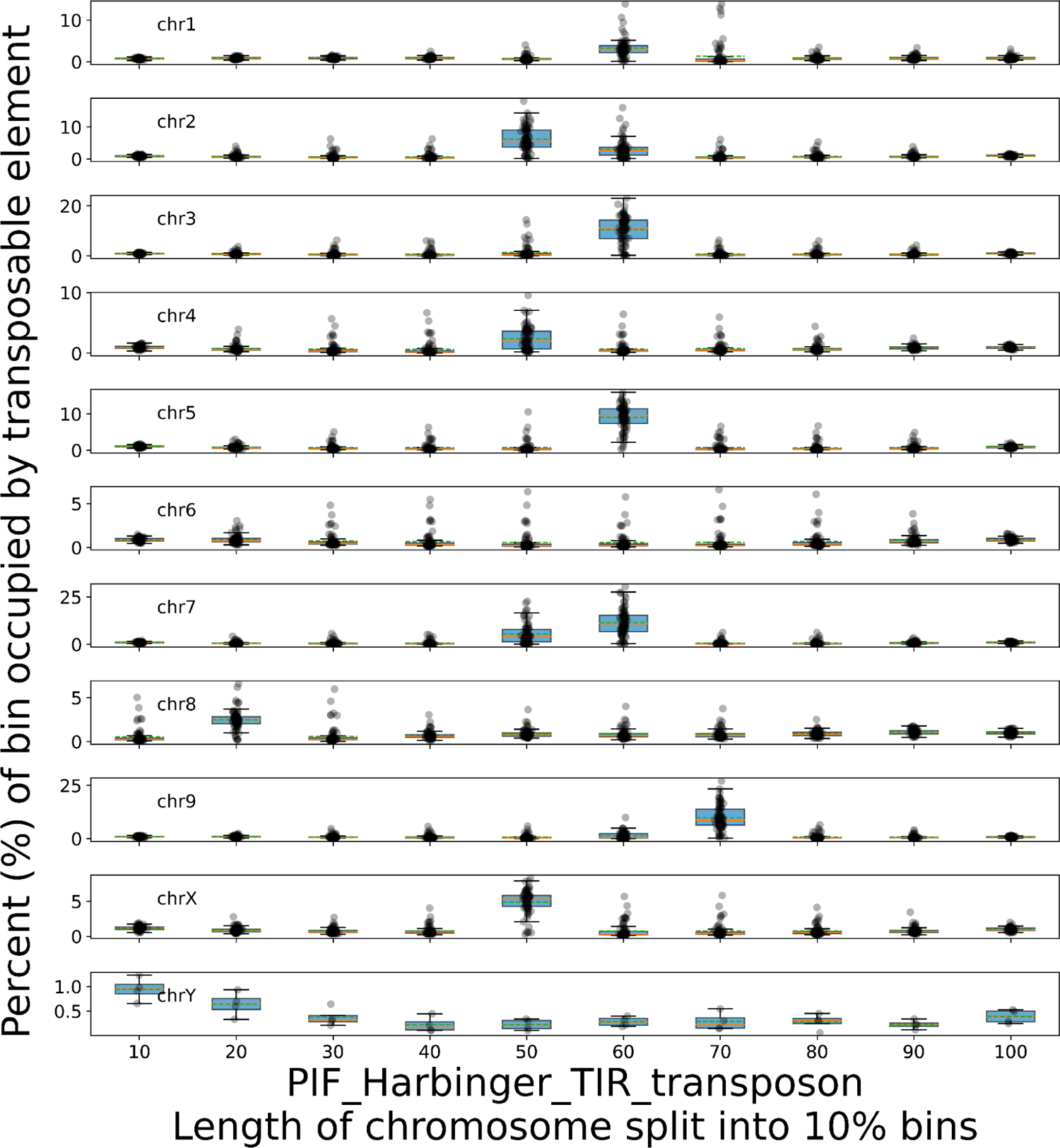
DNA transposable element Harbinger is a common feature of the cannabis centromeres and pericentromeres. Each of the chromosomes of the scaffolded assemblies is divided into sections/bins that each make up 10% of its sequence length, as a way to compare chromosomes with different lengths. Harbinger tends to occur in the center of chromosomes, corresponding to the putative centromere, except for chromosome 8, which is acrocentric.

**Supplemental Figure 15.**
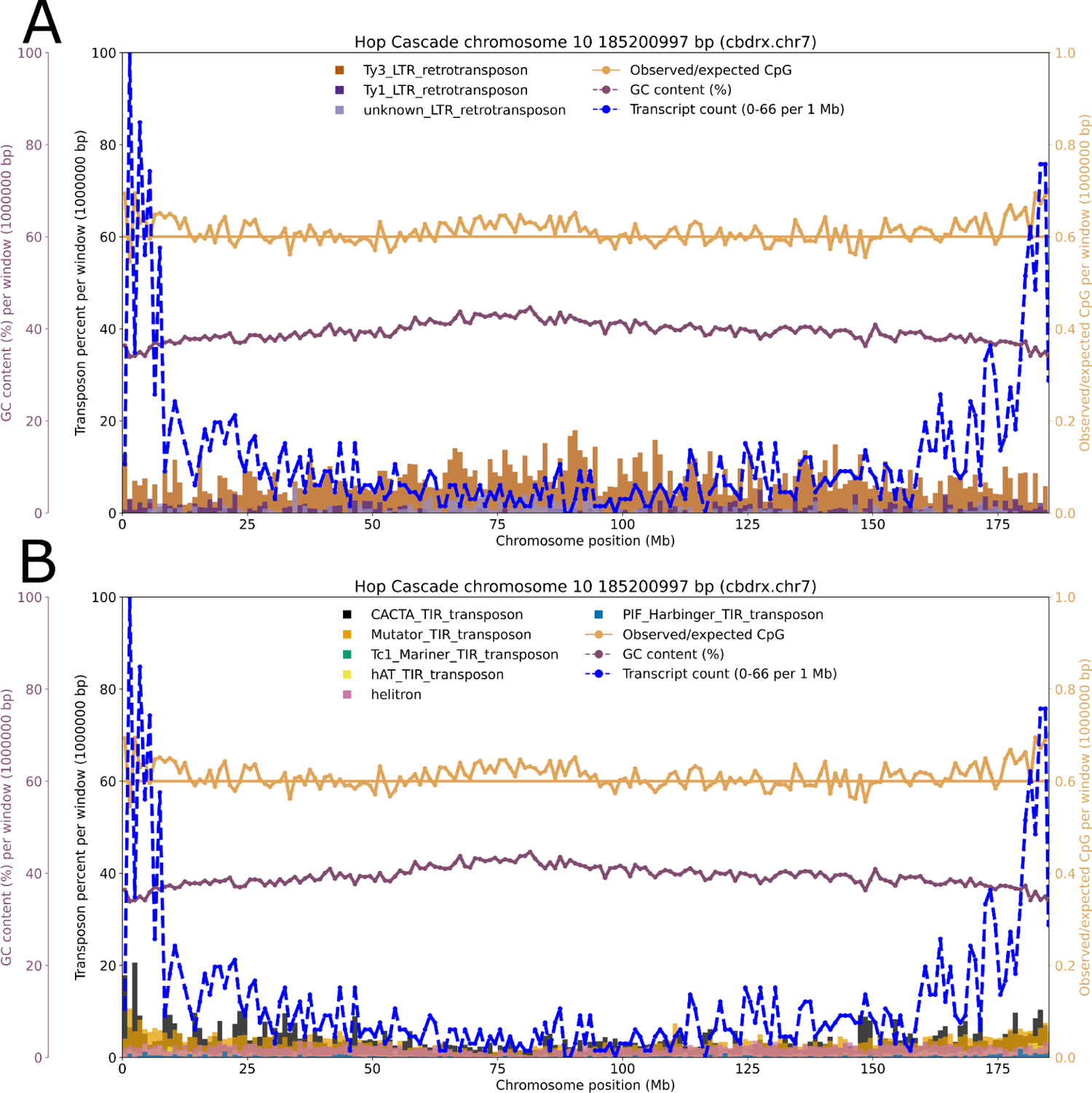
The hop genome, a close relative of cannabis, does not show evidence of a Harbinger-rich centromere. Chromosome/scaffold 10 of the hop genome (*Humulus lupulus*, cultivar “Cascade”) is the homologous chromosome to CBDRx chromosome 7. Hop has high gene content on the scaffold ends. A) Hop chromosome 10 is generally rich in Ty3 LTRs, with a slightly increased abundance in the center of the chromosome. B) Hop chromosome 10 has mostly low abundance of DNA TEs (1 Mb bins).

**Supplemental Figure 16.**
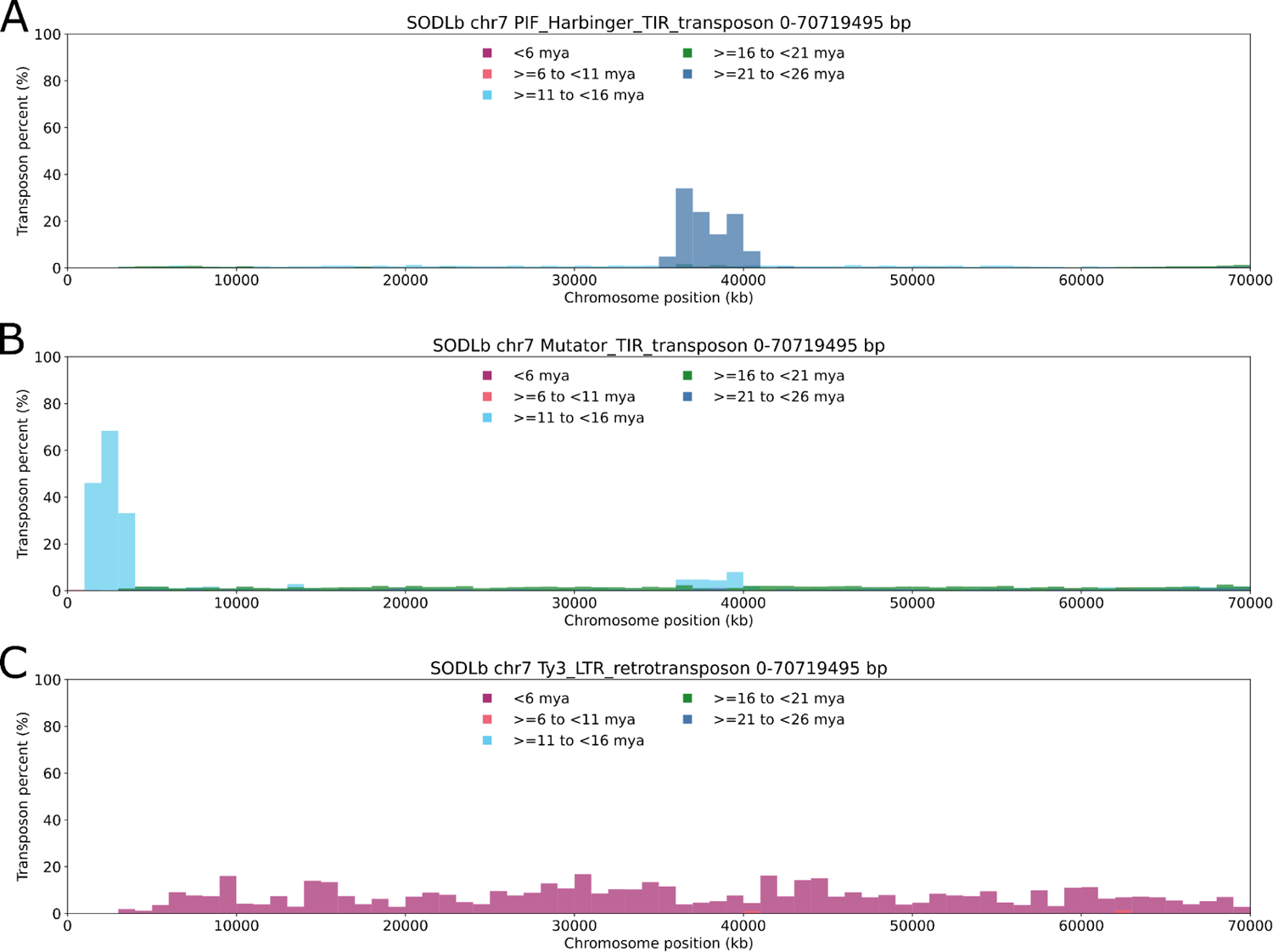
Distribution of TE divergence times across chromosome 7 of genome SODLb. DNA TEs have dynamic distributions that potentially reflect their role in shaping the genome over millions of years. DNA TE Harbinger occurs in the centromere region with a divergence date of ∼20 million years, while Mutator is especially abundant in gene-rich regions, and is more recent than Harbinger (Figure 3 F). In contrast, Ty3-LTR-RTs proliferated recently across the chromosome, with lower abundance in gene-rich regions, as seen in Figure 3 F.

**Supplemental Figure 17.**
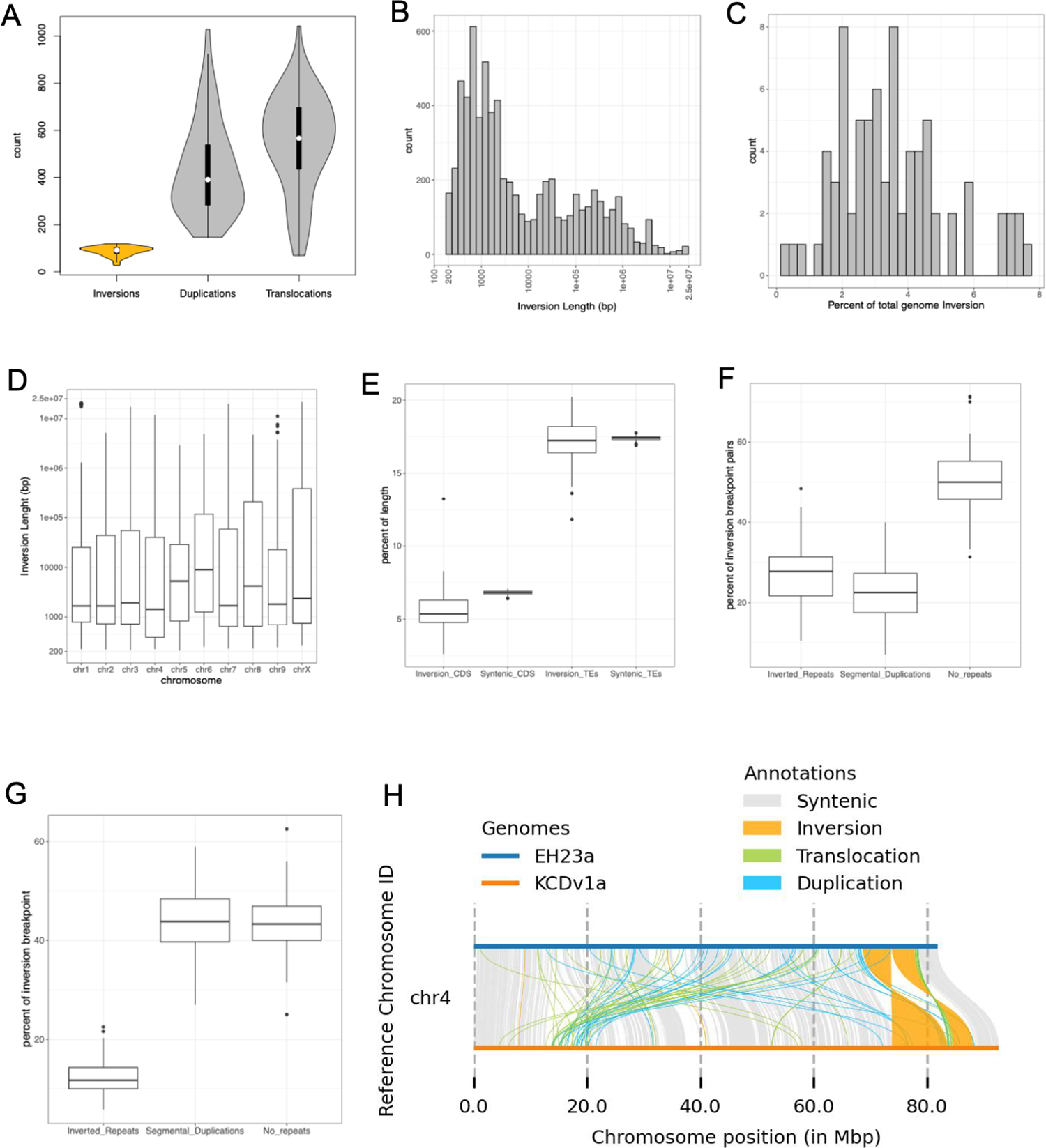
Cannabis pangenome reveals a wide range of structural variation (SV), on par with some of the values that have been reported for interspecies comparisons. A) Distributions of three types of SVs across the 78 scaffolded assemblies of the cannabis pangenome. Each sample assembly was aligned to the EH23a haplotype assembly for SV calling. B) Distribution of the total length of each sample genome assembly in inversions, as a percent of total genome length. C) Multi-modal distribution of inversion lengths, for all inversions from all samples. D) Distributions of inversion lengths, for all inversions from all samples. E) Distributions of coding sequence annotations (CDS) and intact Transposable Elements (TEs) within all inversions and syntenic regions from each sample. Inversions are significantly depleted of CDSs compared to syntenic regions, while on average, TEs are present at nearly an equal level within inversions and syntenic regions. F) Inversion breakpoints (BPs) pairs, defined as 8 kb windows centered at the start and end of each inversion larger than 10 kb, contain repetitive elements about 50% of the time. G) Inversion BPs show a higher rate of segmental duplications, but lower rate of inverted repeats, within self-to-self alignments of each 8 kb BP window, compared to the start-to-end pair alignments. F) Example alignment and SVs of a European hemp sample haplotype (KC Dora). The two mega base scale inversions are in a region of chromosome 4 that showed elevated F_st_ values for SNPs in prior work comparing feral US hemp to marijuana populations ^84^.

**Supplemental Figure 18.**
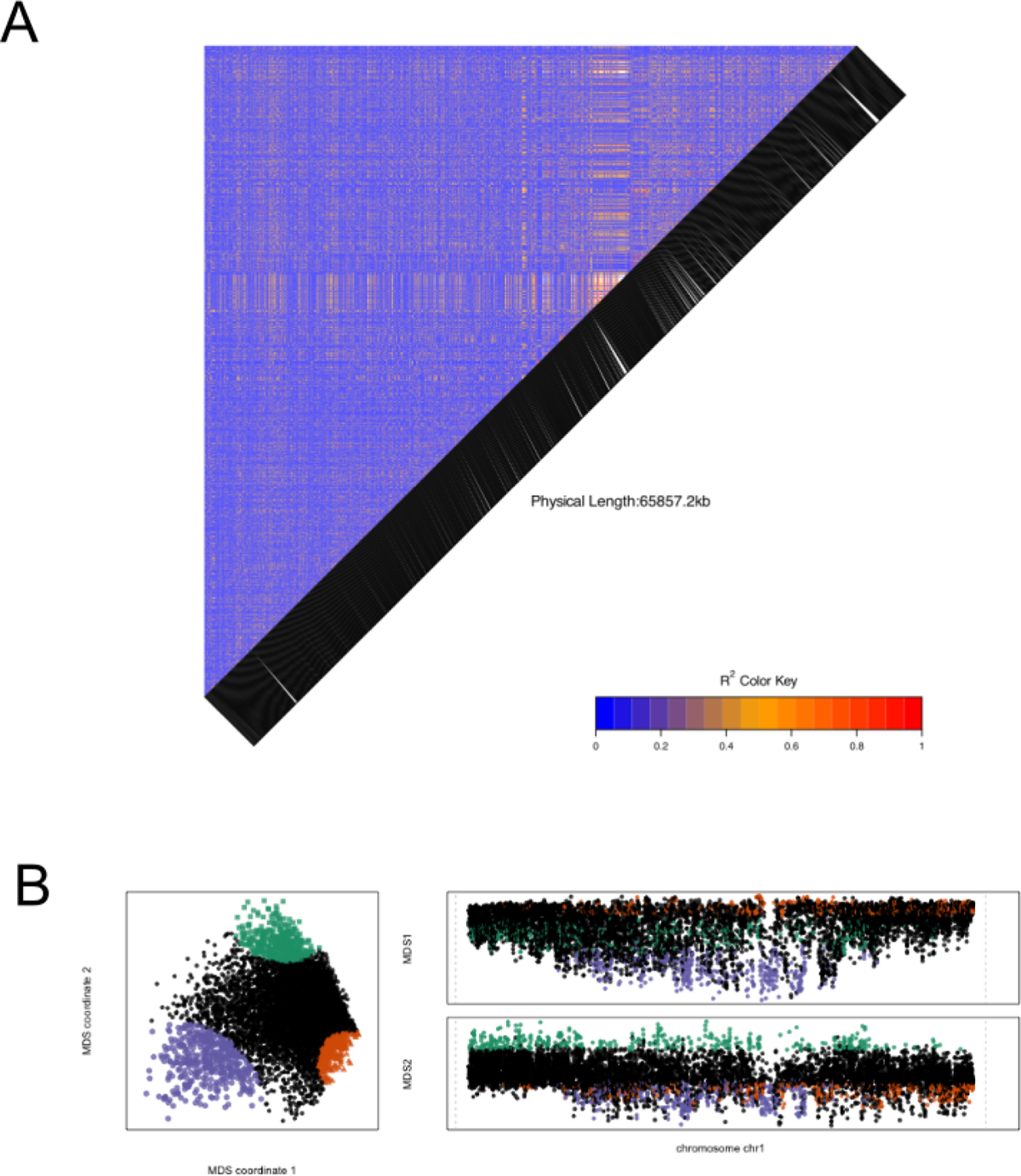
Linkage disequilibrium (LD) and local PCA identify a region on Chromosome 1 resulting in segregation distortion. A) Pairwise SNP R2 heatmap of chromosome 1 showing elevated linkage disequilibrium region that corresponds to the location of the interior breakpoint of the large (∼19.5 MB) inversion. B) Local PCA analysis highlighting an abrupt change in SNP frequencies in the region of the same interior inversion breakpoint.

**Supplemental Figure 19.**
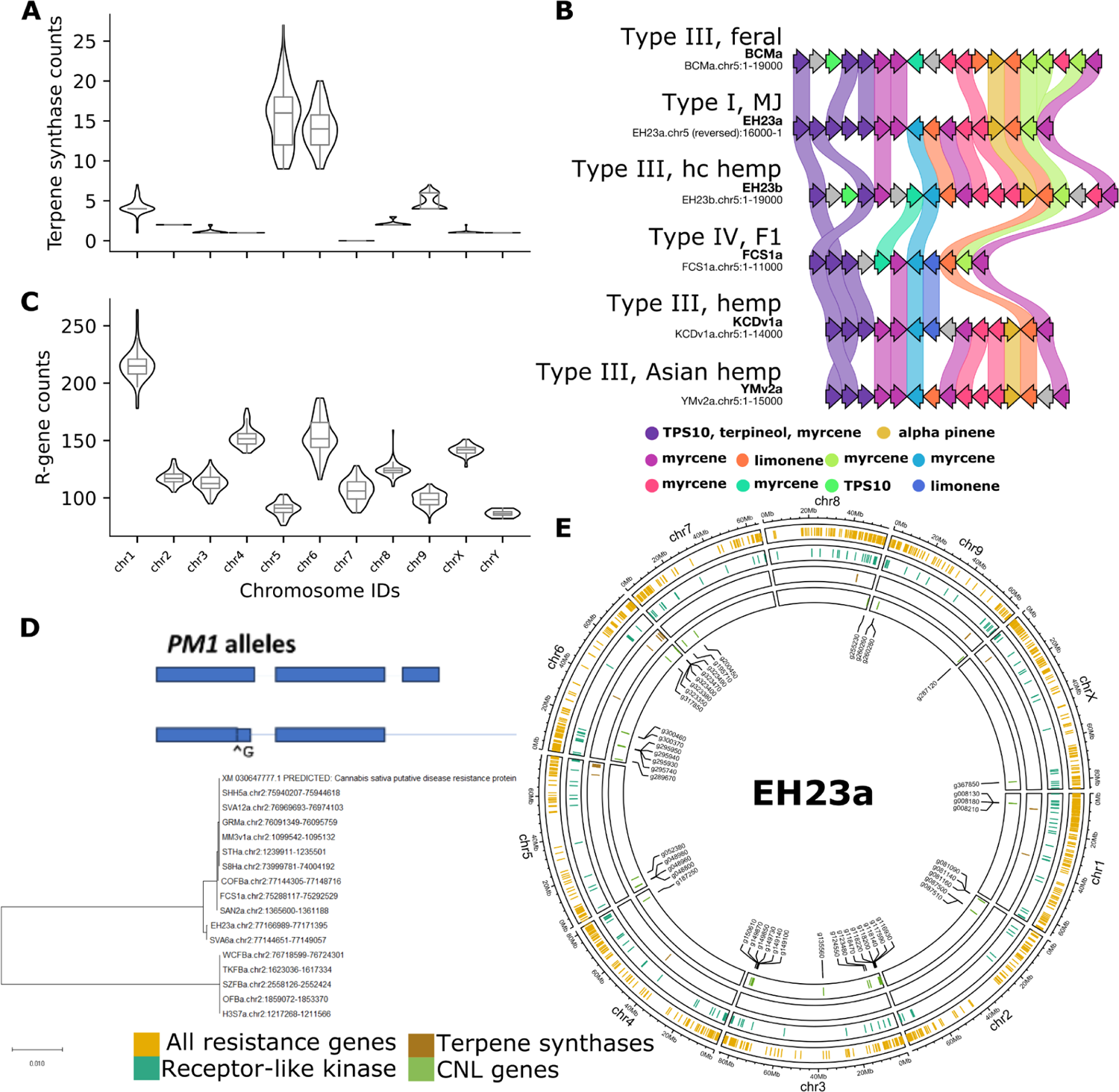
Terpene and disease resistance genes across the cannabis pangenome. A) Violin plot showing terpene synthase copy number in the cannabis pangenome. Chromosomes 5 and 6 are copy number “hotspots” in the cannabis pangenome. B) Organization of terpene synthases on chromosome 5 is consistent, with some variation (not drawn to scale). C) Violin plot showing numbers of resistance gene analogs per chromosome in scaffolded genomes. D) Maximum likelihood tree of CNL genes on chromosome 2 with similarity to a gene associated with powdery mildew resistance. E) Circos plot showing the EH23a genome as an example of the chromosomal distribution of disease resistance gene analogs. Outer track (gold)=all categories of RGAs identified by drago2; middle track (blue)=receptor-like kinases; interior track=coiled-coil nucleotide binding site leucine-rich repeat genes.

**Supplemental Figure 20.**
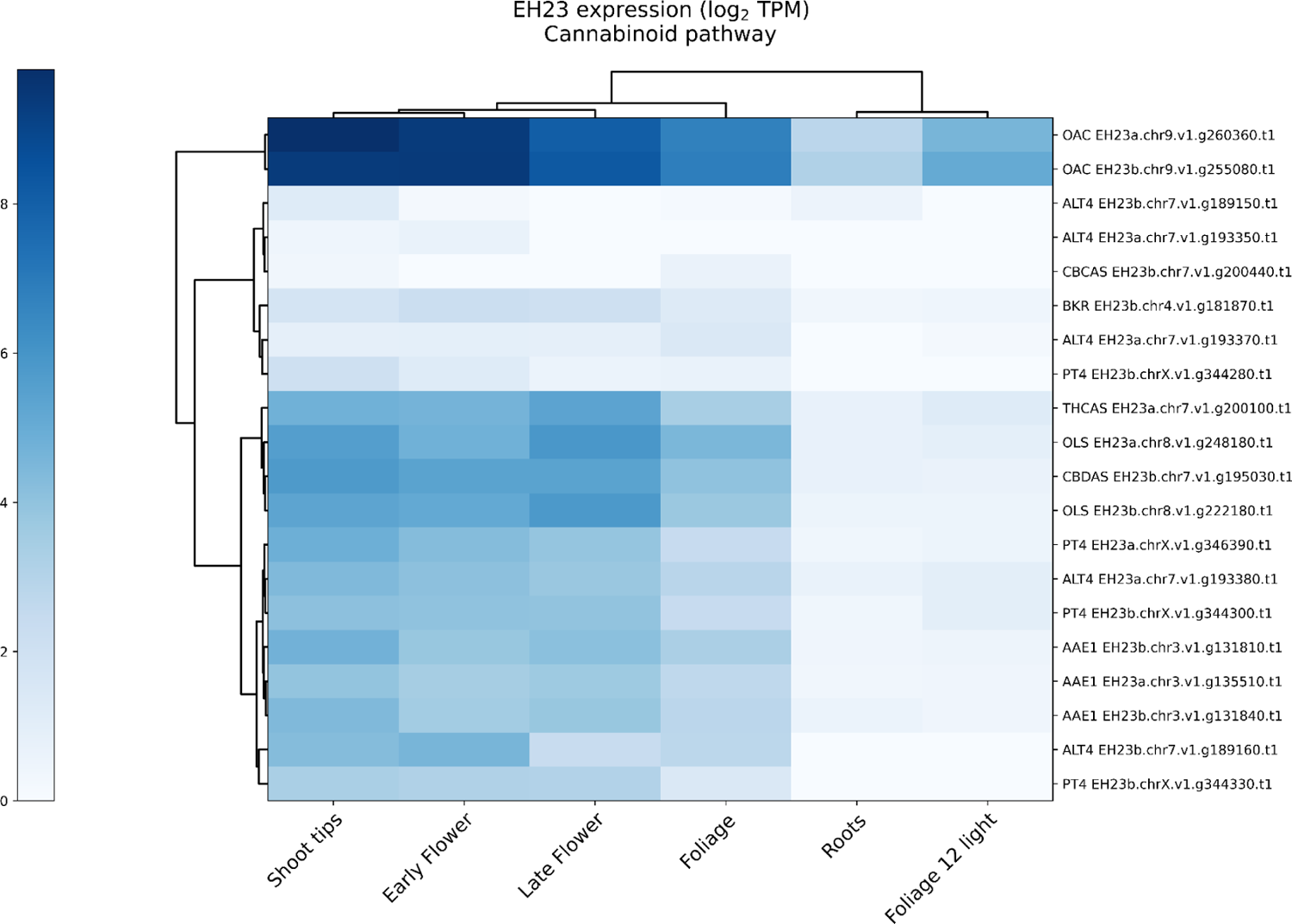
Expression of cannabinoid precursor pathway genes in EH23. Expression values for early and late-stage flowers are clustered together. OAC (olivetolic acid cyclase, UniProt ID I6WU39) is located on chromosome 9 and shows the highest expression. ALT4 and PT4 tend to occur together in the clustered expression heatmap, suggesting a pattern of co-expression and co-regulation. Pathway genes: acyl-activating enzyme (AAE1), olivetolic acid synthase (OLS), olivetolic acid cyclase (OAC), prenyltransferase (PT4; geranylpyrophosphate:olivetolate geranyltransferase), cannabidiolic acid (CBDA), tetrahydrocannabinolic acid (THCA), cannabichromenic acid (CBCA).

**Supplemental Figure 21.**
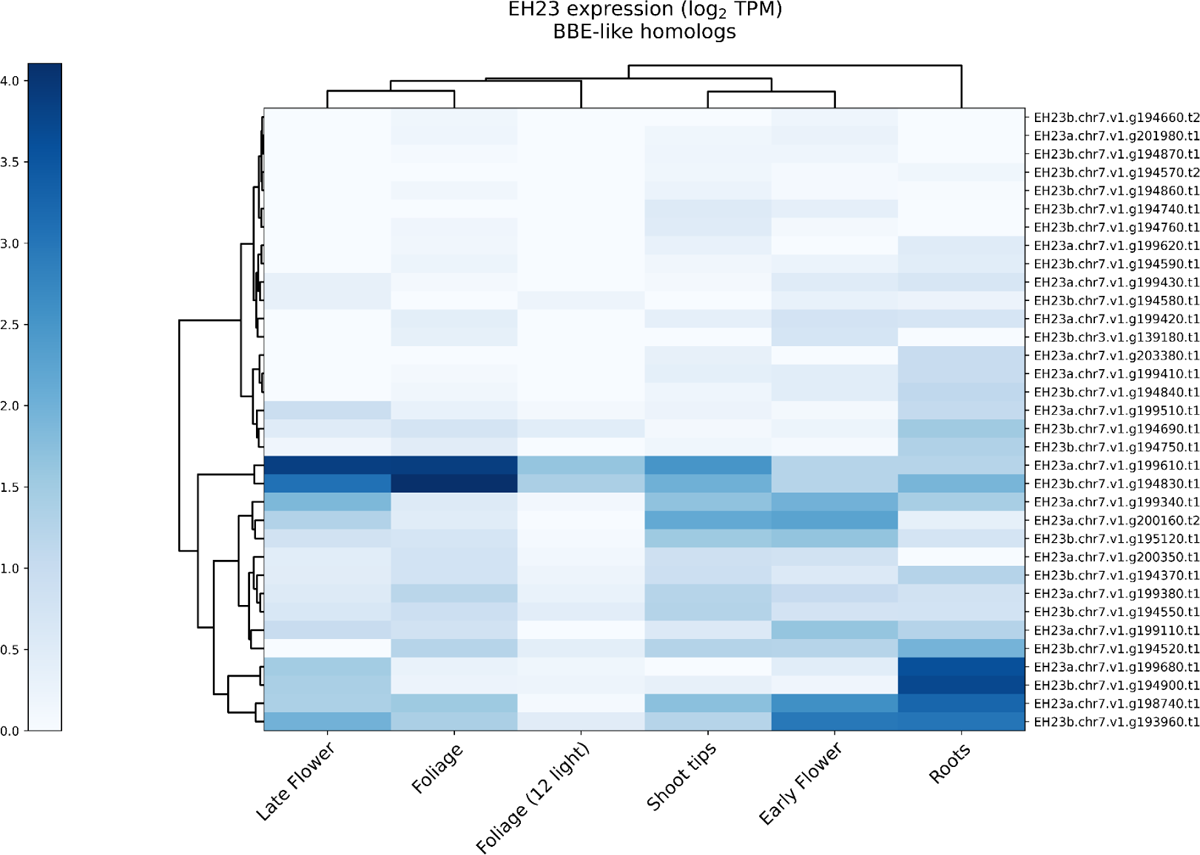
Expression of berberine bridge enzyme (BBE)-like genes in EH23. BBE-like genes are typically involved in alkaloid biosynthesis and are thought to be the ancestral gene of the cannabinoid synthases ^85^, both having an FAD-binding domain. BBE-like genes are almost exclusively expressed on chromosome 7, with foliage, late-stage flower, and roots showing highest expression.

**Supplemental Figure 22.** Cannabinoid synthase cassettes in all assemblies grouped by shape and organization. Overall cassette shape is consistent for each of the full length synthases (CBDAS, CBCAS, THCAS), with moderate variation in the number of partial synthase copies. CBCAS has the greatest variation in the number of full length synthases. However, full length CBDAS and THCAS occur as single copies in all examples. Among the type II genomes, which contain CBDA and THCA synthases in separate haplotypes, the cassette shapes follow type III and type I cassette organization, respectively. See Figshare for full resolution version.

**Supplemental Figure 23.**
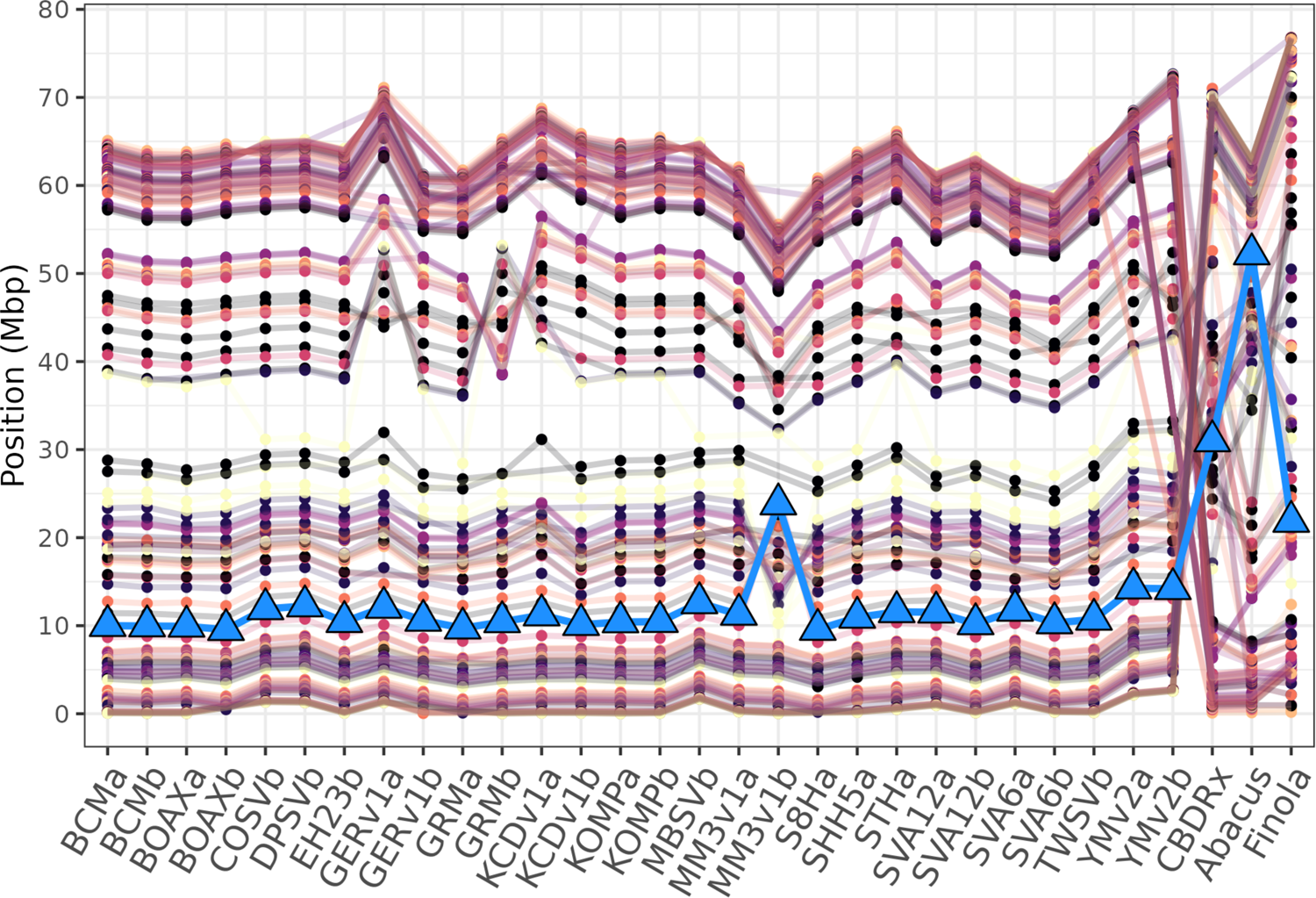
Comparison of BUSCO location and organization for a subset of scaffolded assemblies. Points and lines indicate the location of BUSCO genes (y-axis) on different chromosomes (x-axis). The location of CBDAS (>= 98% identity to AB292682.1) is denoted by the blue triangles and line. The position of CBDAS in CBDRx is approximately 30 Mb, in the pangenome assemblies this gene is consistently located at ∼10 Mb, which suggests that CBDAS is misplaced in CBDRx.

**Supplemental Figure 24.**
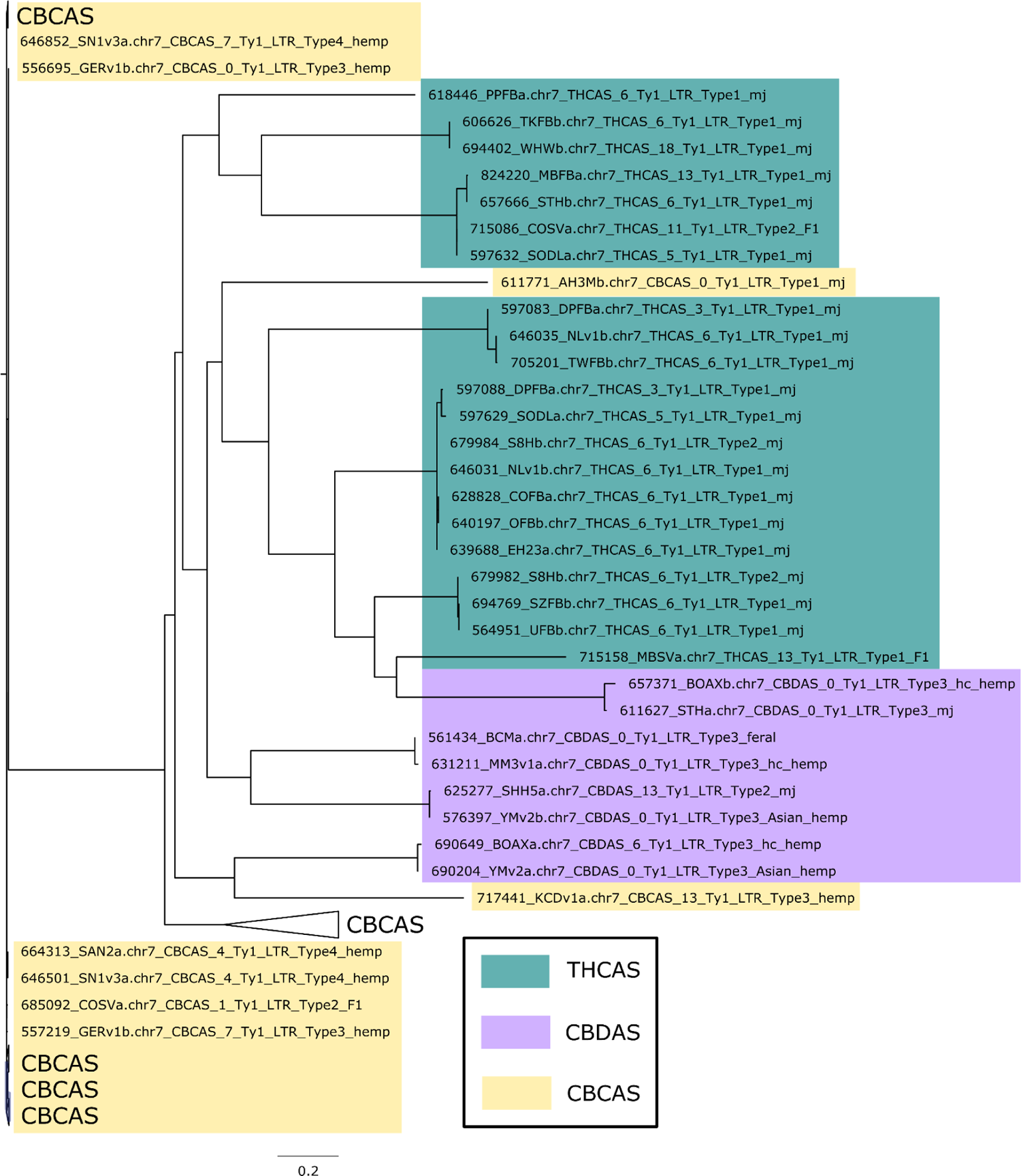
Maximum likelihood tree of Ty1 LTR-RT sequences flanking (2 kb upstream or downstream) cannabinoid synthases in the 78 scaffolded assemblies. Proximal LTR-RT sequences group largely according to full length synthase.

**Supplemental Figure 25.**
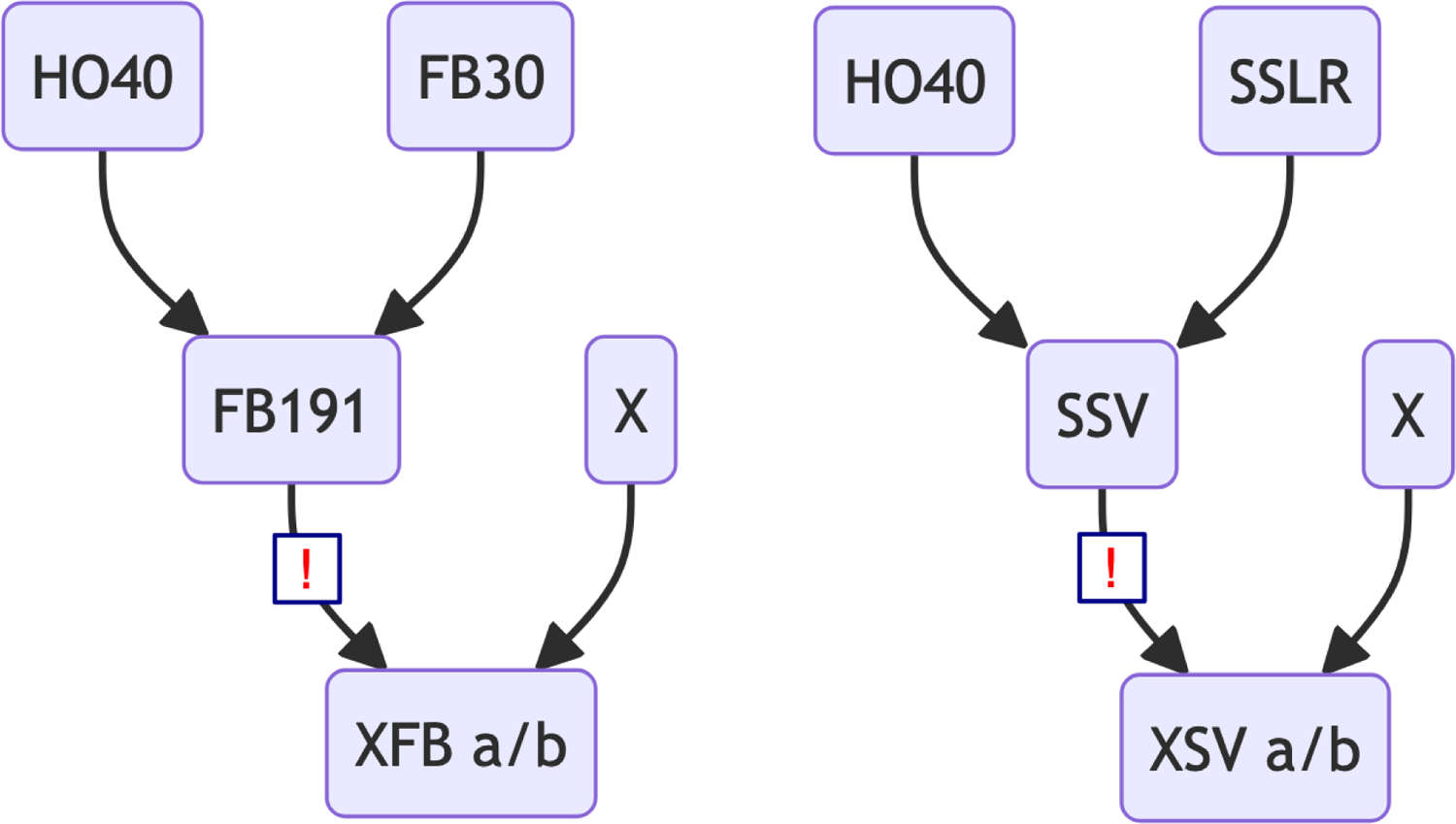
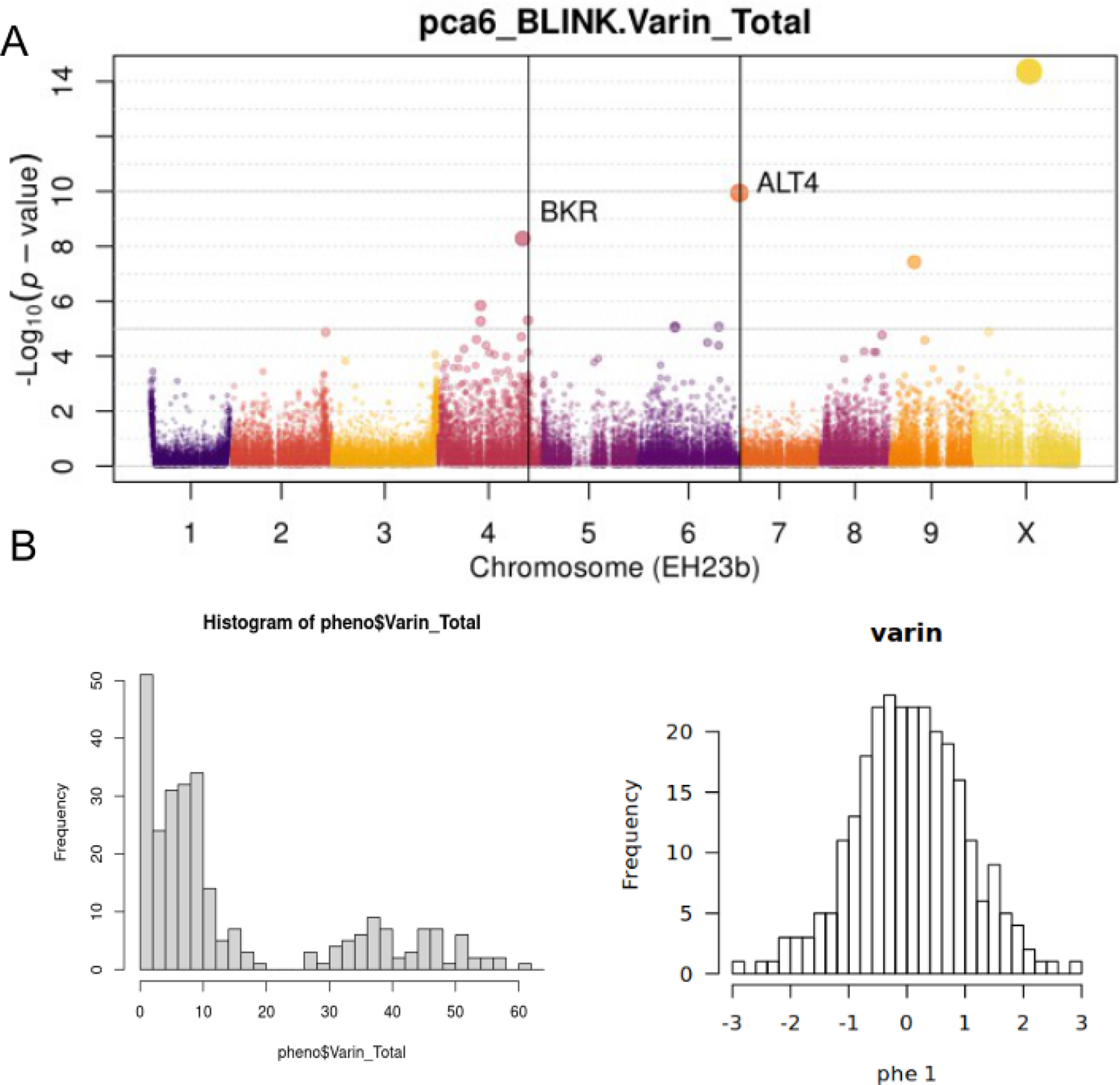
Varin (pentyl) cannabinoid ratio GWAS A) Significant GWA hits, using the BLINK model and normalized varin cannabinoid ratio data. B) Histogram of propyl:pentyl cannabinoid ratios for the F2 population (left), and arcsinh transformed propyl:pentyl cannabinoid ratios (right).

---

**Supplemental Table 1.** Pangenome sample and assembly details: https://figshare.com/articles/figure/Supplementary_Table_1/25869319

**Supplemental Table 2.**
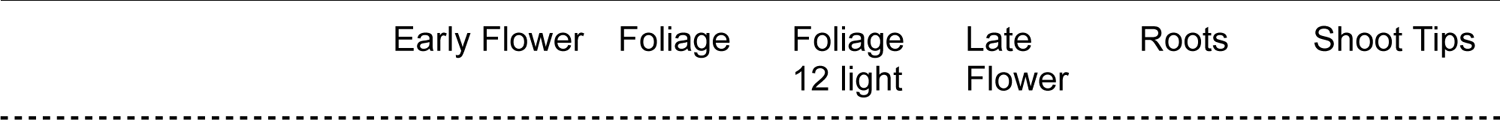
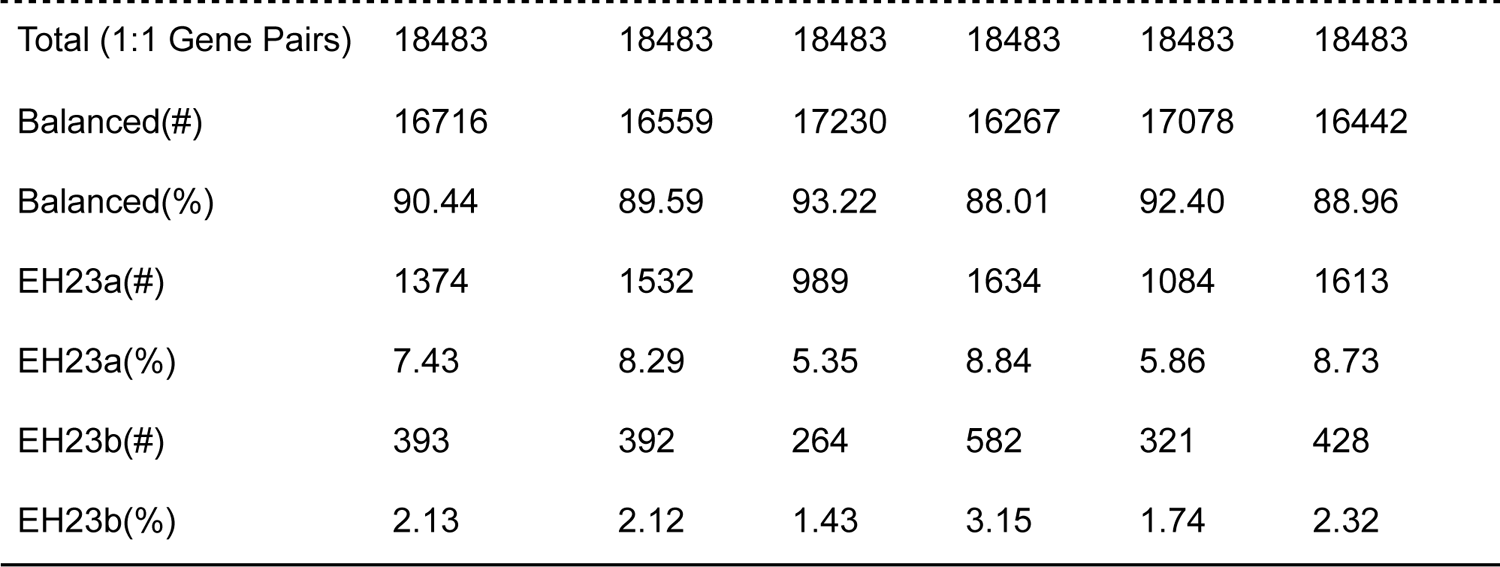
Balanced and biased gene expression in EH23.

| Early Flower | Foliage | Foliage<br>12 light | Late<br>Flower | Roots | Shoot Tips |
| --- | --- | --- | --- | --- | --- |

|  |  |  |  |  |  |  |
| --- | --- | --- | --- | --- | --- | --- |
| Total (1:1 Gene Pairs) | 18483 | 18483 | 18483 | 18483 | 18483 | 18483 |
| Balanced(#) | 16716 | 16559 | 17230 | 16267 | 17078 | 16442 |
| Balanced(%) | 90.44 | 89.59 | 93.22 | 88.01 | 92.40 | 88.96 |
| EH23a(#) | 1374 | 1532 | 989 | 1634 | 1084 | 1613 |
| EH23a(%) | 7.43 | 8.29 | 5.35 | 8.84 | 5.86 | 8.73 |
| EH23b(#) | 393 | 392 | 264 | 582 | 321 | 428 |
| EH23b(%) | 2.13 | 2.12 | 1.43 | 3.15 | 1.74 | 2.32 |

**Supplemental Table 3.** Pairwise Fst values for the populations based on SNPs Weir and Cockerham weighted Fst estimates:

| Comparison | weighted $F_{st}$ |
| --- | --- |
| mj vs feral | 0.21 |
| asian hemp vs mj | 0.21 |
| EU_hemp vs mj | 0.20 |
| asian hemp vs F1s | 0.15 |
| EU_hemp vs asian hemp | 0.14 |
| F1s vs feral | 0.13 |
| EU_hemp vs hc hemp | 0.10 |
| EU_hemp vs F1s | 0.10 |
| mj vs hc hemp | 0.07 |
| asian hemp vs hc hemp | 0.05 |
| EU_hemp vs feral | 0.04 |
| hc hemp vs F1s | 0.04 |
| mj vs F1s | 0.04 |
| hc hemp vs feral | 0.03 |

**Supplemental Table 4.** Average percent of scaffolded assemblies covered by transposable elements.

| Transposable element category | Average repeat content (%) | Standard deviation (%) | Standard deviation to repeat content ratio |
| --- | --- | --- | --- |

|  |  |  | (metric of variation in % repeat content) |
| --- | --- | --- | --- |
| Total repeat content | 67.89 | 1.19 | 0.02 |
| Ty1 LTR-RT | 16.27 | 1.68 | 0.1 |
| Ty3 LTR-RT | 19.7 | 1.32 | 0.07 |
| Unknown LTR-RT | 16.51 | 2.46 | 0.15 |
| CACTA | 3.12 | 1.59 | 0.51 |
| Harbinger | 1.09 | 0.32 | 0.29 |
| hAT | 1.95 | 1.21 | 0.62 |
| helitron | 2.84 | 1.68 | 0.59 |
| Mariner | 0.37 | 0.55 | 1.49 |
| Mutator | 6.03 | 1.45 | 0.24 |

**Supplemental Table 5.** Average distance between genes and transposable elements in scaffolded assemblies.

| Transposable element category | Median distance (bp) | Average distance (bp) | Standard deviation (bp) |
| --- | --- | --- | --- |
| Ty1 LTR-RT | 238 | 491.91 | 840.17 |
| Ty3 LTR-RT | 138 | 476.49 | 968.93 |
| Unknown LTR-RT | 279 | 538.85 | 1251.94 |
| CACTA | 103 | 443.83 | 940.63 |
| Harbinger | 335 | 551.38 | 916.6 |
| hAT | 359 | 613.82 | 1025.01 |
| helitron | 312 | 549.16 | 890.42 |
| Mariner | 142 | 506.79 | 1270.51 |
| Mutator | 274 | 517.64 | 897.37 |

**Supplemental Table 6.** Average methylation (%) across the genome.

| Genome | Population information | Mean methylation level (%) across the | Mean methylation level (%) across the |
| --- | --- | --- | --- |

|  |  | genome in haplotype A | genome in haplotype B |
| --- | --- | --- | --- |
| AH3M | MJ, type 1 | 84.37 | 87.47 |
| KCDv1 | Hemp, type 3 | 83.35 | 84.37 |
| NLv1 | MJ, type 1 | 84.36 | 87.10 |
| WHW | MJ, type 1 | 79.64 | 81.65 |
| SODL | MJ, type 1 | 84.45 | 85.73 |
| SN1v3 | Hemp, type 4 | 73.82 | 80.36 |
| SAN2 | Hemp, type 4 | 73.10 | 80.04 |
| FCS1 | F1, type 1 (FCS1a), type 4 (FCS1b) | 78.0 | 82.10 |
| H3S1 | F1, type 1 (H3S1a), type 4 (H3S1b) | 80.33 | 82.99 |

**Supplemental Table 7.**
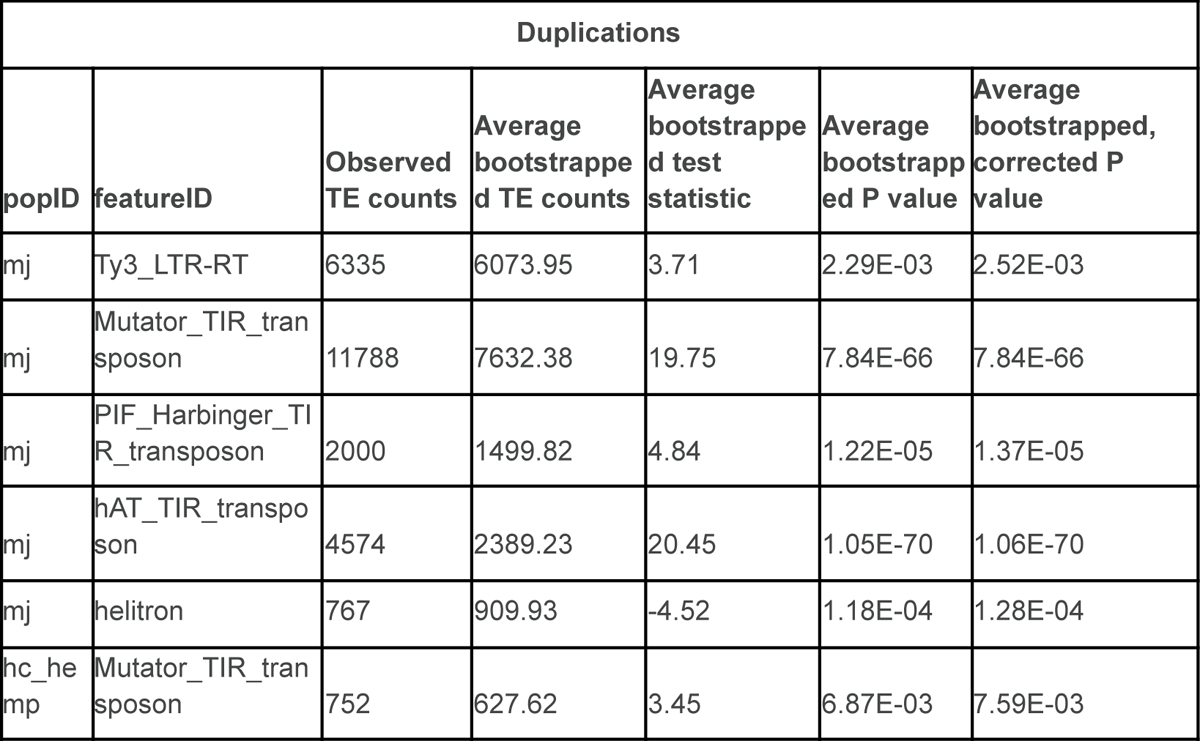
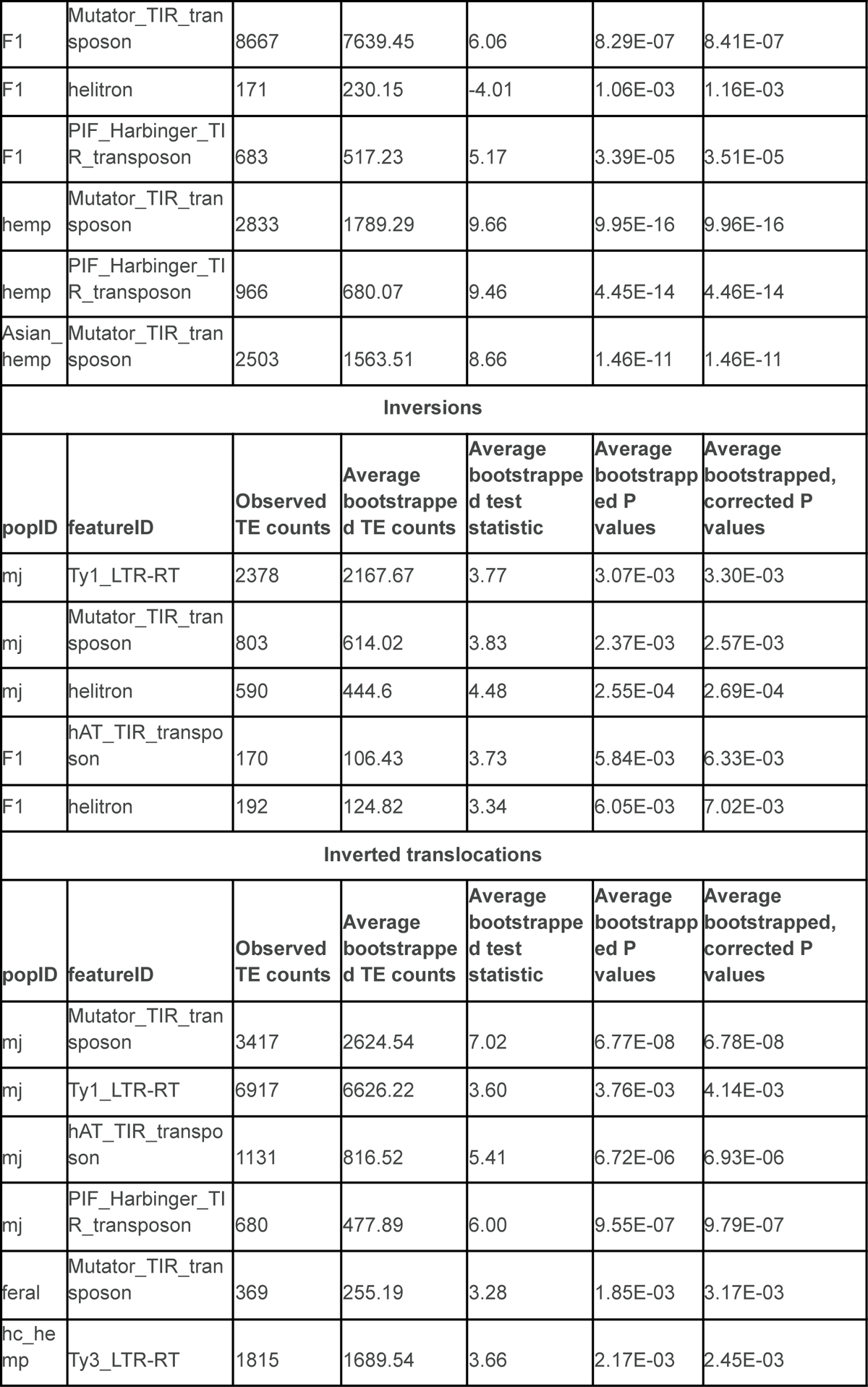
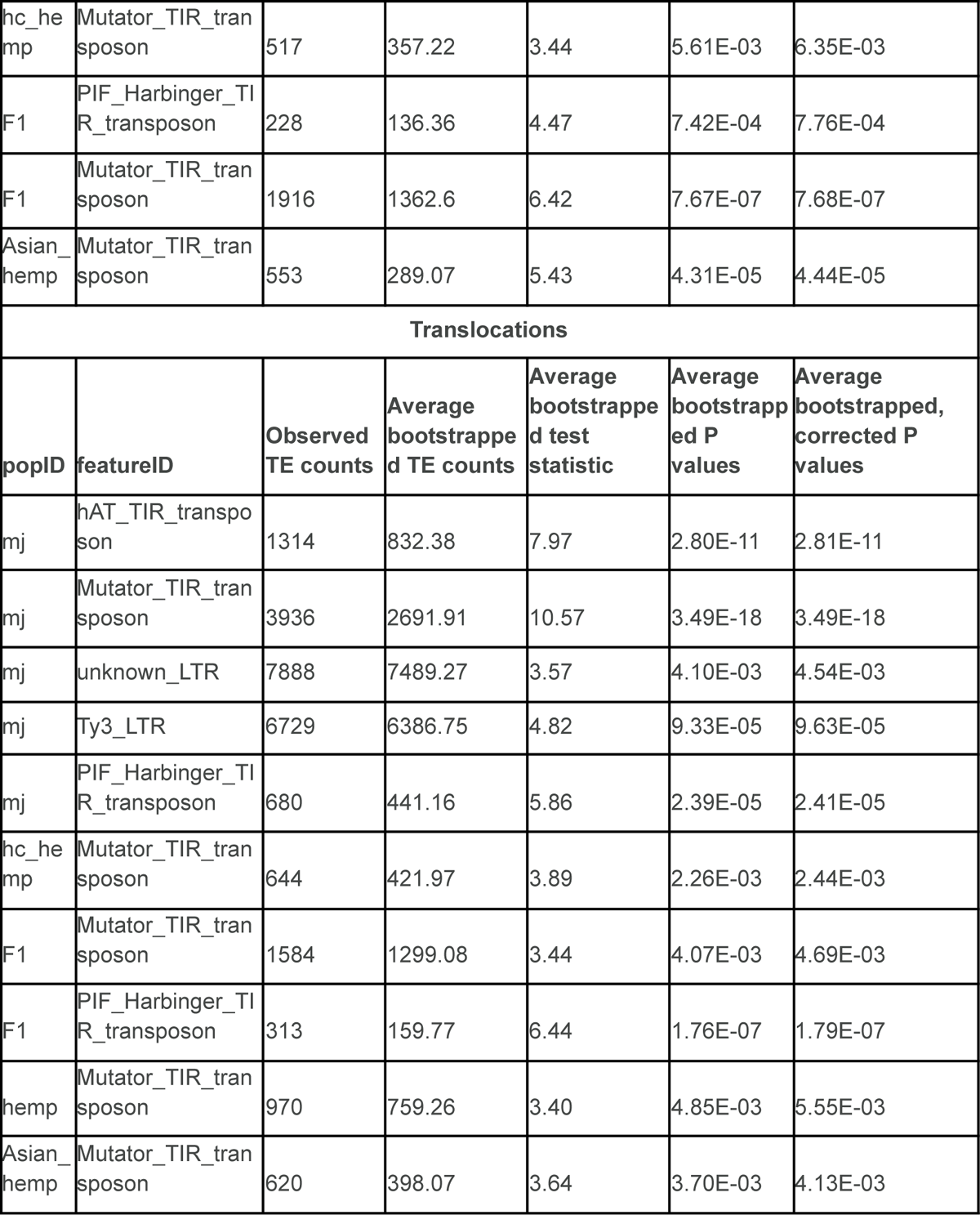
This table includes the number of TEs associated with breakpoints of SVs (duplications, inversions, inverted translocations, and translocations). The observed TE count is the total number of TEs associated with a given SV among the genomes in each population. The average bootstrapped TE count is the total number of TEs associated with a random region of the genome that has the same length and is from the same chromosome as an observed SV. A positive test statistic indicates TE enrichment and negative test statistic indicate TE depletion.

| Duplications |  |  |  |  |  |  |
| --- | --- | --- | --- | --- | --- | --- |
| popID | featureID | Observed TE counts | Average bootstrapped TE counts | Average bootstrapped test statistic | Average bootstrapped P value | Average bootstrapped, corrected P value |
| mj | Ty3_LTR-RT | 6335 | 6073.95 | 3.71 | 2.29E-03 | 2.52E-03 |
| mj | Mutator_TIR_transposon | 11788 | 7632.38 | 19.75 | 7.84E-66 | 7.84E-66 |
| mj | PIF_Harbinger_TIR_transposon | 2000 | 1499.82 | 4.84 | 1.22E-05 | 1.37E-05 |
| mj | hAT_TIR_transposon | 4574 | 2389.23 | 20.45 | 1.05E-70 | 1.06E-70 |
| mj | helitron | 767 | 909.93 | -4.52 | 1.18E-04 | 1.28E-04 |
| hc_hemp | Mutator_TIR_transposon | 752 | 627.62 | 3.45 | 6.87E-03 | 7.59E-03 |
| F1 | Mutator_TIR_transposon | 8667 | 7639.45 | 6.06 | 8.29E-07 | 8.41E-07 |
| F1 | helitron | 171 | 230.15 | -4.01 | 1.06E-03 | 1.16E-03 |
| F1 | PIF_Harbinger_TIR_transposon | 683 | 517.23 | 5.17 | 3.39E-05 | 3.51E-05 |
| hemp | Mutator_TIR_transposon | 2833 | 1789.29 | 9.66 | 9.95E-16 | 9.96E-16 |
| hemp | PIF_Harbinger_TIR_transposon | 966 | 680.07 | 9.46 | 4.45E-14 | 4.46E-14 |
| Asian_hemp | Mutator_TIR_transposon | 2503 | 1563.51 | 8.66 | 1.46E-11 | 1.46E-11 |
| **Inversions** |  |  |  |  |  |  |
| **popID** | **featureID** | **Observed TE counts** | **Average bootstrapped TE counts** | **Average bootstrapped test statistic** | **Average bootstrapped P values** | **Average bootstrapped, corrected P values** |
| mj | Ty1_LTR-RT | 2378 | 2167.67 | 3.77 | 3.07E-03 | 3.30E-03 |
| mj | Mutator_TIR_transposon | 803 | 614.02 | 3.83 | 2.37E-03 | 2.57E-03 |
| mj | helitron | 590 | 444.6 | 4.48 | 2.55E-04 | 2.69E-04 |
| F1 | hAT_TIR_transposon | 170 | 106.43 | 3.73 | 5.84E-03 | 6.33E-03 |
| F1 | helitron | 192 | 124.82 | 3.34 | 6.05E-03 | 7.02E-03 |
| **Inverted translocations** |  |  |  |  |  |  |
| **popID** | **featureID** | **Observed TE counts** | **Average bootstrapped TE counts** | **Average bootstrapped test statistic** | **Average bootstrapped P values** | **Average bootstrapped, corrected P values** |
| mj | Mutator_TIR_transposon | 3417 | 2624.54 | 7.02 | 6.77E-08 | 6.78E-08 |
| mj | Ty1_LTR-RT | 6917 | 6626.22 | 3.60 | 3.76E-03 | 4.14E-03 |
| mj | hAT_TIR_transposon | 1131 | 816.52 | 5.41 | 6.72E-06 | 6.93E-06 |
| mj | PIF_Harbinger_TIR_transposon | 680 | 477.89 | 6.00 | 9.55E-07 | 9.79E-07 |
| feral | Mutator_TIR_transposon | 369 | 255.19 | 3.28 | 1.85E-03 | 3.17E-03 |
| hc_hemp | Ty3_LTR-RT | 1815 | 1689.54 | 3.66 | 2.17E-03 | 2.45E-03 |

| hc_he<br>mp | Mutator_TIR_transposon | 517 | 357.22 | 3.44 | 5.61E-03 | 6.35E-03 |
| --- | --- | --- | --- | --- | --- | --- |
| F1 | PIF_Harbinger_TIR_transposon | 228 | 136.36 | 4.47 | 7.42E-04 | 7.76E-04 |
| F1 | Mutator_TIR_transposon | 1916 | 1362.6 | 6.42 | 7.67E-07 | 7.68E-07 |
| Asian_hemp | Mutator_TIR_transposon | 553 | 289.07 | 5.43 | 4.31E-05 | 4.44E-05 |
| **Translocations** |  |  |  |  |  |  |
| popID | featureID | Observed TE counts | Average bootstrapped TE counts | Average bootstrapped test statistic | Average bootstrapped P values | Average bootstrapped, corrected P values |
| mj | hAT_TIR_transposon | 1314 | 832.38 | 7.97 | 2.80E-11 | 2.81E-11 |
| mj | Mutator_TIR_transposon | 3936 | 2691.91 | 10.57 | 3.49E-18 | 3.49E-18 |
| mj | unknown_LTR | 7888 | 7489.27 | 3.57 | 4.10E-03 | 4.54E-03 |
| mj | Ty3_LTR | 6729 | 6386.75 | 4.82 | 9.33E-05 | 9.63E-05 |
| mj | PIF_Harbinger_TIR_transposon | 680 | 441.16 | 5.86 | 2.39E-05 | 2.41E-05 |
| hc_he<br>mp | Mutator_TIR_transposon | 644 | 421.97 | 3.89 | 2.26E-03 | 2.44E-03 |
| F1 | Mutator_TIR_transposon | 1584 | 1299.08 | 3.44 | 4.07E-03 | 4.69E-03 |
| F1 | PIF_Harbinger_TIR_transposon | 313 | 159.77 | 6.44 | 1.76E-07 | 1.79E-07 |
| hemp | Mutator_TIR_transposon | 970 | 759.26 | 3.40 | 4.85E-03 | 5.55E-03 |
| Asian_hemp | Mutator_TIR_transposon | 620 | 398.07 | 3.64 | 3.70E-03 | 4.13E-03 |

**Supplemental Table 8.** Support for gene models.

| Cultivar | Tissue | Sequencing method | NCBI BioProject |
| --- | --- | --- | --- |
| Sour Diesel, Canna Tsu, Black Lime, Valley Fire, White Cookies, Mama Thai, Terple, Cherry Chem, Blackberry Kush | Flower | Short-read Illumina | PRJNA498707 |
| ERB x H040 (EH23) | Roots, foliage, foliage collected at 12 hours<br>lights on/12 hours<br>lights off, shoot tips,<br>early-stage flower,<br>late-stage flower | Short-read Illumina | Pangenome release |
| Fortune Cookie x<br>Santhica 27<br>(FC6xSan27, FCS1) | Flower (collected at<br>5:00AM, 8:00AM,<br>3:00PM, 8:00PM),<br>Leaf (collected at<br>5:00AM, 8:00AM,<br>3:00PM, 8:00PM)<br>Stem (collected at<br>5:00AM, 8:00PM)<br>Small roots (collected<br>at 8:00AM)<br>Tap root (collected at<br>8:00AM) | Full-length Oxford<br>Nanopore cDNA | Pangenome release |
| Hercules x Santhica<br>27<br>(H35xSan27,<br>H3S1) | Flower (collected at<br>5:00AM, 8:00AM,<br>8:00PM)<br>Leaf (collected at<br>5:00AM, 8:00AM,<br>3:00PM, 8:00PM)<br>Small roots (collected<br>at 8:00AM)<br>Tap root (collected at<br>8:00AM) | Full-length Oxford<br>Nanopore cDNA | Pangenome release |
| Santhica 27 (SAN2) | Flower (collected at<br>8:00AM [2 reps],<br>8:00PM)<br>Leaf (collected at<br>5:00AM, 8:00PM)<br>Stem (collected at<br>8:00PM)<br>Small roots (collected<br>at 8:00AM) | Full-length Oxford<br>Nanopore cDNA | Pangenome release |
|  | Tap root (collected at 8:00AM) |  |  |

**Supplemental Table 9.**
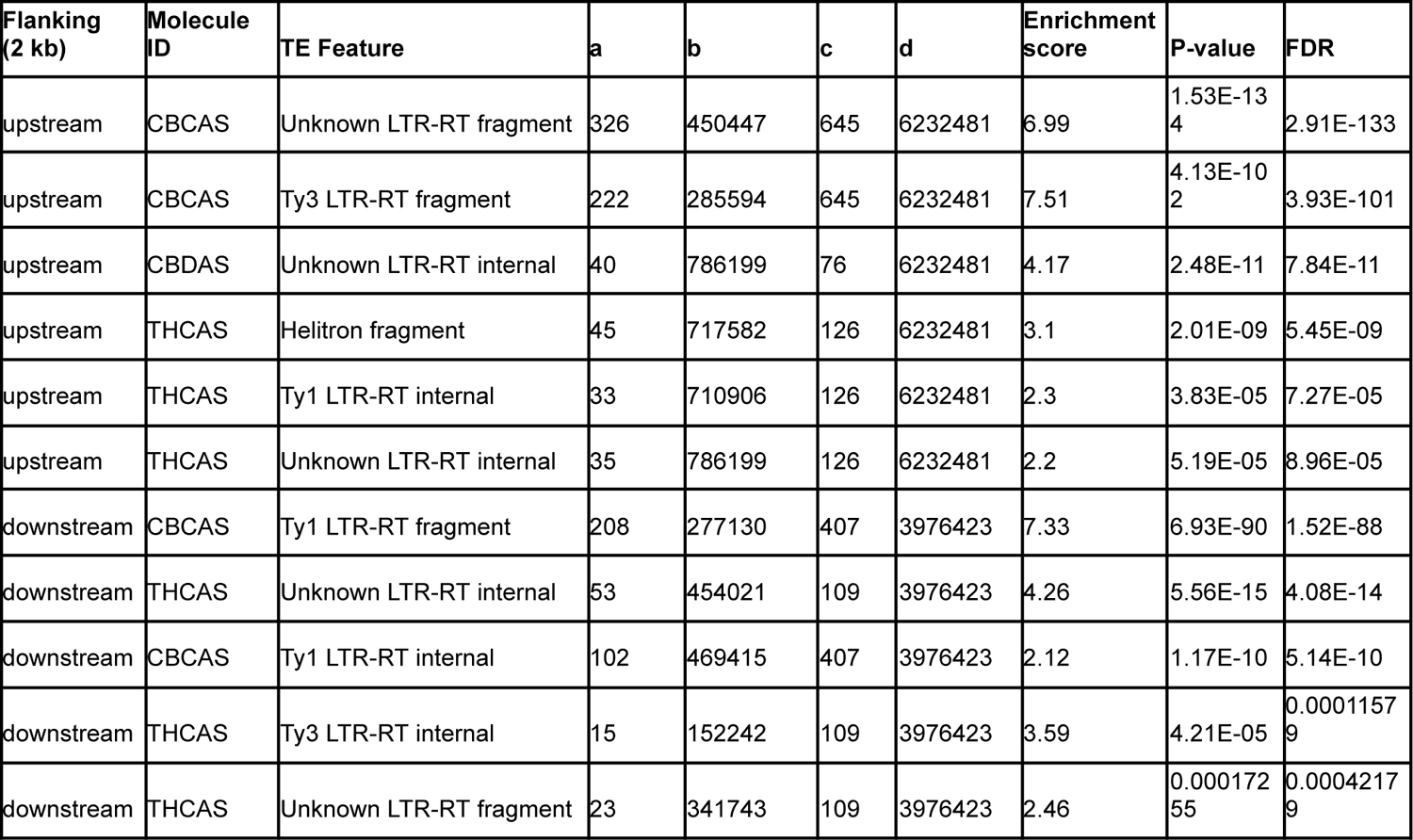
Statistically significant TEs upstream (2 kb) and downstream (2 kb) of cannabinoid synthases.

**Supplemental Table 10.** Trios used in crossover analysis of varin.

| Parent 1 | Parent 2 | Cross | Varin haplotype |
| --- | --- | --- | --- |
| FB191 |  | COFB | a |
| . |  | DPFB | b |
| . |  | MBFB | b |
| . |  | OFB | b |
| . |  | PPFB | b |
| . |  | SDFB | b |
| . |  | SKFB | b |
| . |  | TKFB | b |
| . |  | TWFB | b |
| . |  | UFB | b |
| . |  | WCFB | b |
| SSV |  | COSV | b |
| . |  | DPSV | b |
| . |  | MBSV | b |
| . |  | SVA6 | a |
| . |  | SVA12 | a |
|  |  | TWSVb | b |

**Supplemental Table 11.**
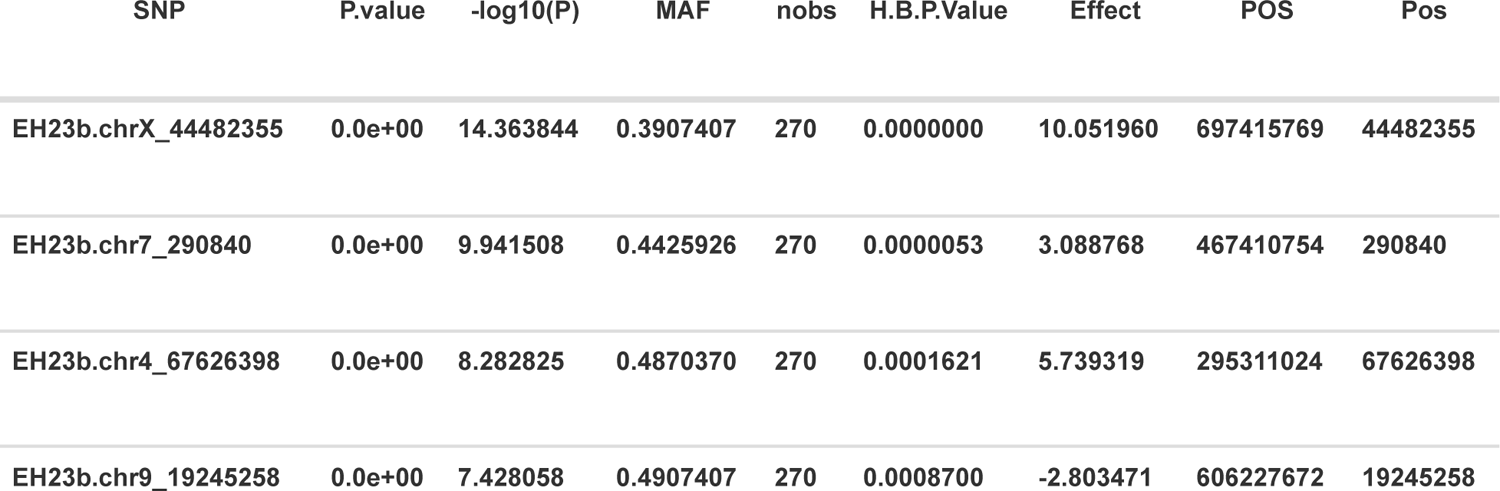
Significant GWA hits, using the BLINK model and normalized varin cannabinoid ratio data.

## Notes

https://figshare.com/projects/Cannabis_Pangenome/205555

https://resources.michael.salk.edu/root/home.html

https://github.com/ViningLab/CannabisPangenome

https://github.com/anthony-aylward/CannabisPangenomeShared/tree/main

